# Characterisation of cell-scale signalling by the core planar polarity pathway during *Drosophila* wing development

**DOI:** 10.1101/2024.10.14.618159

**Authors:** Alexandre Carayon, Helen Strutt, David Strutt

## Abstract

In developing epithelia, cells become planar polarised through asymmetric localisation of the core planar polarity proteins to opposite cell membranes, where they form stable intercellular complexes. Current models differ regarding the signalling mechanisms required for core protein polarisation. Here, we investigate the existence of cell-intrinsic cell-scale signalling *in vivo* in the *Drosophila* pupal wing. We use conditional and restrictive expression tools to spatiotemporally manipulate core protein activity, combined with quantitative measurement of core protein distribution, polarity and stability. Our results provide evidence for a robust cell-scale signal, while arguing against mechanisms that depend on depletion of a limited pool of a core protein or polarised transport of core proteins on microtubules. Furthermore, we show that polarity propagation across a tissue is hard, highlighting the strong intrinsic capacity of individual cells to establish and maintain planar polarity.

## Introduction

Planar polarity (also known as planar cell polarity [PCP]) describes the coordinated polarisation of cells within the tissue plane, and is established by molecular mechanisms that are conserved throughout the animal kingdom (Hale and Strutt, 2015). Examples of planar polarised structures are hair follicles in the skin and cilia on epithelia such as lung, while planar polarity also controls polarised cell movements during gastrulation that promote axis elongation and neural tube closure (Goodrich and Strutt, 2011; Devenport, 2016; Butler and Wallingford, 2017; Davey and Moens, 2017).

Planar polarity has been extensively studied in the *Drosophila* wing, where each cell is planar polarised and produces a distally oriented trichome (Adler, 2002). This depends on activity of the so-called ‘core proteins’: six proteins that form polarised intercellular complexes at the proximo-distal cell junctions (Goodrich and Strutt, 2011; Devenport, 2014; Harrison et al., 2020). Core protein complexes on the distal cell membrane are composed of the transmembrane protein Frizzled (Fz) and the cytoplasmic proteins Dishevelled (Dsh) and Diego (Dgo), whereas on proximal cell membranes they contain the transmembrane protein Strabismus (Stbm, also known as Van Gogh [Vang]) and the cytoplasmic protein Prickle (Pk), while the transmembrane protein Flamingo (Fmi, also known as Starry Night [Stan]) is localised on both apposing cell membranes (Fig.1A). This core protein distribution on opposite cell membranes in the hexagonal cells of the wing epithelium results in a characteristic zig-zag localisation pattern (Fig.1B). Disruption to activity of any single core protein affects this asymmetric pattern of localisation and trichomes no longer point distally (Harrison et al., 2020).

**Figure 1.**
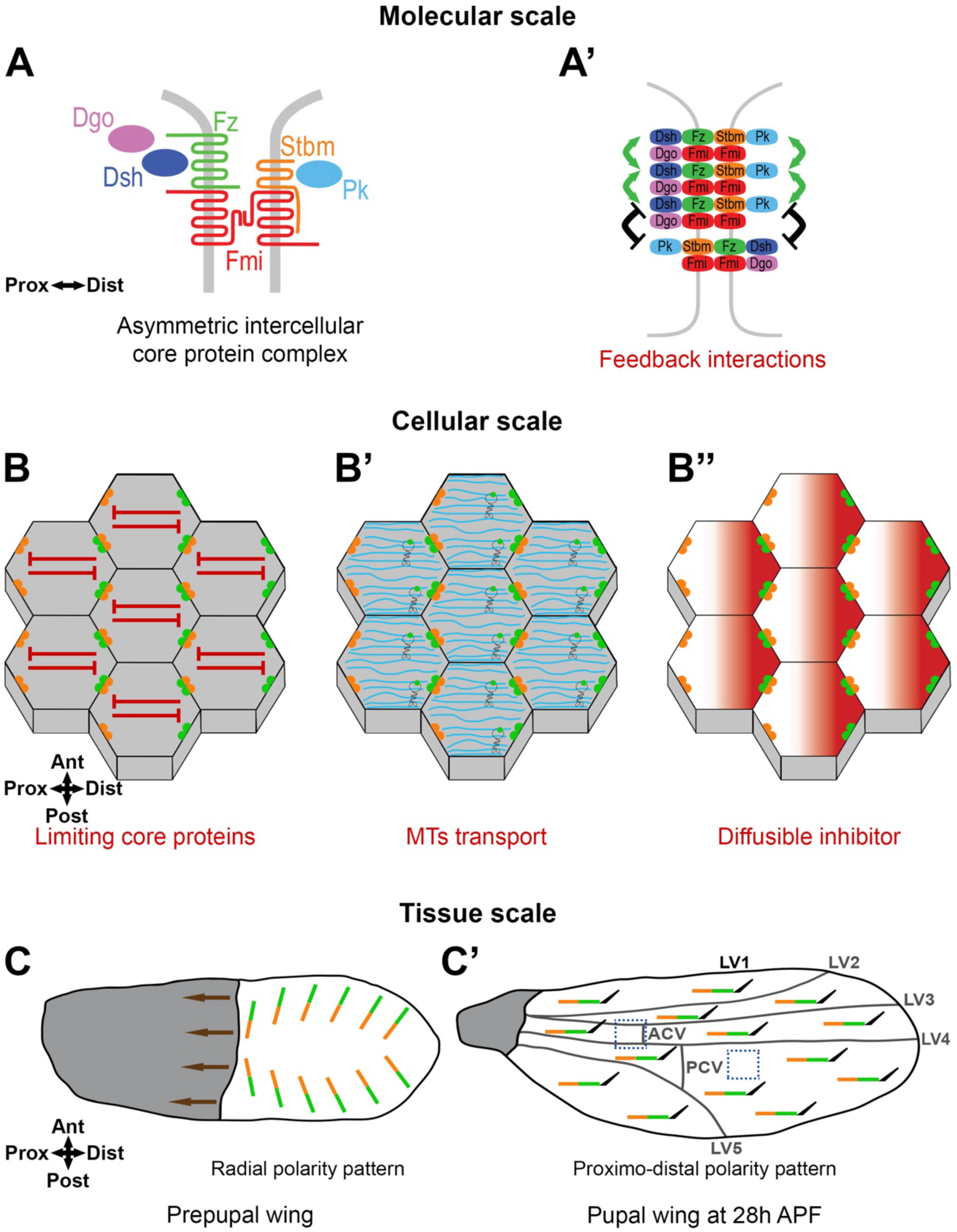
Planar Polarity in the *Drosophila* wing. **(A)** Core proteins form asymmetric intercellular protein complexes at apicolateral cell membranes mediating intercellular communication. Frizzled (Fz, green) a sevenpass transmembrane protein, and the cytoplasmic proteins Dishevelled (Dsh, dark blue) and Diego (Dgo, magenta) localise to distal cell membranes, while Strabismus (Stbm, orange, also known as Van Gogh [Vang]) a fourpass transmembrane protein and Prickle (Pk, light blue) a cytoplasmic protein, localise to proximal cell membranes. Flamingo (Fmi, red, also known as Starry Night [Stan]) an atypical sevenpass transmembrane cadherin, localises proximally and distally, forming a trans homodimer. **(A’)** Core protein complexes are thought to interact between themselves through feedback interactions locally on cell junctions (‘Molecular-scale’) to form stable clusters, with positive interactions stabilising complexes of the same orientation (green arrows) or negative interactions destabilising complexes of opposite orientation (black symbols). **(B-B’’)** Core protein complexes are segregated to opposite cell membranes generating the specific proximo-distal polarised zig-zag core protein localisation pattern which can be promoted by ‘Cell-scale signalling’. In this study we consider several hypotheses to identify such cell-scale signals, such as being mediated through depletion of a limiting pool of a core protein providing long-range inhibition (red symbols) **(B)**, through oriented MTs transport (MTs in cyan, with motor proteins) **(B’)**, or by a biochemical mechanism such as diffusion of an inhibitor **(B’’)**. **(C, C’)** In the 28h pupal wing, core protein polarity (represented by lines with Stbm in orange and Fz in green) and trichome (black hair) orientation is distal **(C’)**. It is suggested that this results from radial global cues generating an initial asymmetric bias across the tissue. During earlier polarity establishment in the prepupal wing, hinge contraction (brown arrows) induces cell rearrangements and a redistribution of the core proteins, from the radial polarity pattern in the prepupal wing **(C)** to aligned core protein polarity along the proximo-distal axis of the pupal wing by 28h APF **(C’)**.

The orientation of planar polarity in the developing *Drosophila* pupal wing is governed by processes occurring at different scales (Aw and Devenport, 2017; Harrison et al., 2020; Strutt and Strutt, 2021). At the tissue level there are ‘global cues’, which act to provide an overall direction to planar polarity (Aw and Devenport, 2017). The identities of the global cues in the wing are still under investigation, however experimental studies have revealed that planar polarity is initially present in a radial pattern pointing towards the pupal wing margin and is then re-ordered into a proximo-distal pattern as a result of mechanical tension from contraction of the wing hinge that induces polarised cell flows and cell rearrangements (Aigouy et al., 2010; Tan and Strutt, 2025) (reviewed in Eaton and Jülicher, 2011; Aw and Devenport, 2017; Butler and Wallingford, 2017) (Fig.1C,C’). The origin of the earlier radial pattern is under debate, with evidence being presented both for and against instructive roles for gradients of secreted Wnt ligands or Fat-Dachsous cadherin expression (e.g. Ma et al., 2003; Matakatsu and Blair, 2004; Casal et al., 2006; Matakatsu and Blair, 2006; Sagner et al., 2012; Wu et al., 2013; Ewen-Campen et al., 2020; Yu et al., 2020). In addition, experiments in which core pathway activity is only activated at timepoints later than hinge contraction, suggest an absence of any proximo-distal polarity cues in the developing wing other than the tissue tension induced by the hinge (Strutt and Strutt, 2002; Strutt and Strutt, 2007; Brittle et al., 2022). Specifically such ‘*de novo* induction’ of pathway activity results in strong swirling patterns of polarity across the surface of the wing that do not respect any specific tissue axis.

The initial polarity bias produced by global cues is then thought to be locally amplified by feedback interactions between core protein complexes on specific cell junctions (‘Molecular-scale’ [Fig.1A’]) (Usui et al., 1999; Axelrod, 2001; Strutt, 2001; Brittle et al., 2022). Complexes of the same orientation are suggested to be stabilised by positive feedback interactions whereas complexes of the opposite orientation are destabilised by negative feedback interactions. This leads to sorting of stable complexes into a uniform orientation on proximal and distal cell membranes (Harrison et al., 2020; Strutt and Strutt, 2021).

Computational studies support the idea that global cues combined with local feedback interactions are sufficient to establish coordinated planar polarity across a tissue (Amonlirdviman et al., 2005; Le Garrec et al., 2006; Fischer et al., 2013). However, they also predict that in the absence of a global cue, local feedback interactions are insufficient to polarise cells. This stands in contrast to the *in vivo* studies showing that *de novo* induction of core pathway activity after the onset of hinge contraction results in swirling patterns of polarity that appear uncoupled from global cues (Strutt and Strutt, 2002; Strutt and Strutt, 2007; Brittle et al., 2022). To explain the production of such *de novo* patterns of swirling polarity in the absence of global cues, the existence of ‘non-local’ cell-scale signalling has been suggested (Meinhardt, 2007; Burak and Shraiman, 2009; Abley et al., 2013; Shadkhoo and Mani, 2019) (Fig.1B). In this context, ‘cell-scale signalling’ describes possible intracellular mechanisms acting to promote core protein segregation to opposite cell membranes. For example, this could be a signal from ‘distal complexes’ at one side of the cell leading to segregation of ‘proximal complexes’ to the opposite cell edge, or *vice versa*.

Despite the theoretical and experimental evidence for its existence, there has so far been no systematic investigation into cell-scale signalling in planar polarity. In this manuscript we use the establishment of proximo-distal planar polarity in the *Drosophila* wing as a model system to explore possible mechanisms. Based on one suggested scenario (Meinhardt, 2007), we tested a ‘depletion’ model whereby one or more core proteins is present in limiting amounts for cell for polarisation, thus acting as a non-local inhibitory signal (Fig.1B). To do this, we set up a system in which we can induce core pathway *de novo* polarisation while varying core pathway gene dosage. We coupled this with use of tagged Fz that acts as a fluorescent timer to discriminate different Fz populations over time. As an alternative possibility, we assessed whether the previously reported transport of core proteins on microtubules (MTs) (Shimada et al., 2006; Harumoto et al., 2010; Matis et al., 2014) could be acting as a cell-scale signal during polarity establishment (Fig.1B’). Finally, we studied polarity propagation across the tissue caused by induction of a boundary of Fz expression, in an attempt to characterise the properties of the putative cell-scale signal.

Our results argue against roles in mediation of a cell-scale signal for either depletion of a limited pool of a core protein or polarised transport of core proteins on microtubules. Nevertheless, our studies of polarity propagation from boundaries reveal that such propagation is hard to induce even over timescales of hours. This suggests that cells possess an intrinsic capacity to polarise and maintain their polarity, implying the existence of a robust cell-scale signal (Fig.1B’’).

## Results

### Dynamics of *de novo* planar polarity establishment

To understand mechanisms of core pathway planar polarity establishment, we first followed the process of cell polarisation over time, measuring three parameters: (i) cell polarity orientation (polarity angle), (ii) cell polarity magnitude (polarity strength), and (iii) core protein complex stability. Cell polarity orientation and magnitude were determined using the PCA method in QuantifyPolarity as described previously (Tan et al., 2021). Complex stability was assessed by measuring the dynamics of turnover of the core protein Fz. Fz interacts with Fmi to form the asymmetric backbone of core protein complexes (Strutt and Strutt, 2007; Chen et al., 2008; Strutt and Strutt, 2008; Strutt et al., 2023) and thus its stability is a proxy for overall complex stability. In our experiments Fz is tagged with two distinct fluorescent proteins whose maturation timings are different: sfGFP (superfolder GFP) with a very rapid maturation, showing the total population of Fz, and mKate2 with a slower maturation revealing older (presumed stable) populations of Fz, to form a fluorescent timer (Khmelinskii et al., 2012; Barry et al., 2016; Ressurreicao et al., 2018) (Fig.S1A). The ratio of mKate2/sfGFP is used as a proxy for the proportion of stable Fz.

In wild-type pupal wings, the pattern of core pathway planar polarity depends upon (i) pre-existing polarity established at earlier stages of development, (ii) cell flows and cell re-arrangements that occur during pupal wing morphogenesis (Aigouy et al., 2010; Tan and Strutt, 2025). To attempt to bypass effects of these upstream polarity cues, we studied planar polarisation in the *de novo* polarisation context (Strutt and Strutt, 2002; Strutt and Strutt, 2007; Brittle et al., 2022), generated by induction of Fz expression in a *fz* null mutant background at stages of pupal wing development when cell flows and cell re-arrangements are largely complete (see Introduction).

We used the FLP/FRT recombination system (*FRT-STOP-FRT* cassette excision in the presence of *hsFLP*) (Theodosiou and Xu, 1998) combined with a 1h heat shock (38°C) (Fig.S1B) to induce Fz::mKate2-sfGFP expression under the direct control of the *Actin5C* promoter ubiquitously throughout the wing in a *fz* null mutant background. To confirm that Fz::mKate2-sfGFP had reached steady state levels at our chosen experimental timepoints, we monitored Fz protein by western blot analysis at multiple time points after *de novo* induction. At 6h, 8h and 10h after induction, levels were not significantly different from each other, or from our control condition of constitutive Fz::mKate2-sfGFP levels (Fig.S1C-D). Fz::mKate2-sfGFP levels were several fold lower than endogenous Fz, however, this does not affect the ability of Fz::mKate2-sfGFP to achieve a normal polarity magnitude (see below). We initially analysed planar polarity establishment in a flat part of the wing, below longitudinal vein 4 (LV4) and distal to the posterior cross vein (PCV) (Fig.1C’).

To characterise the rate of polarisation and establishment of stable polarity complexes during *de novo* polarity establishment, we first needed reference measurements for polarised and unpolarised cells. For fully polarised cells, we examined wings that constitutively expressed Act-Fz::mKate2-sfGFP throughout development (Fig.2D). Live imaging, with observation of native sfGFP and mKate2 fluorescence, showed Fz asymmetric localisation at cell junctions in the pupal wing, leading to the specific zig-zag core protein localisation pattern (Fig.2I’-I’’), with a proximo-distal polarity orientation of Fz (Fig.2I’’’) and a uniform proximo-distal orientation of trichomes in the adult wing (Fig.2L), in agreement with previous work (Strutt, 2001). Moreover, the magnitude of Fz polarity achieved as measured by the PCA method was similarly to that seen previously for endogenously-tagged Fz-EGFP (Fig.2J) (Tan et al., 2021). Hence mKate2-sfGFP tagged Fz expressed under the *Actin5C* promoter recapitulates endogenous Fz function.

**Figure 2.**
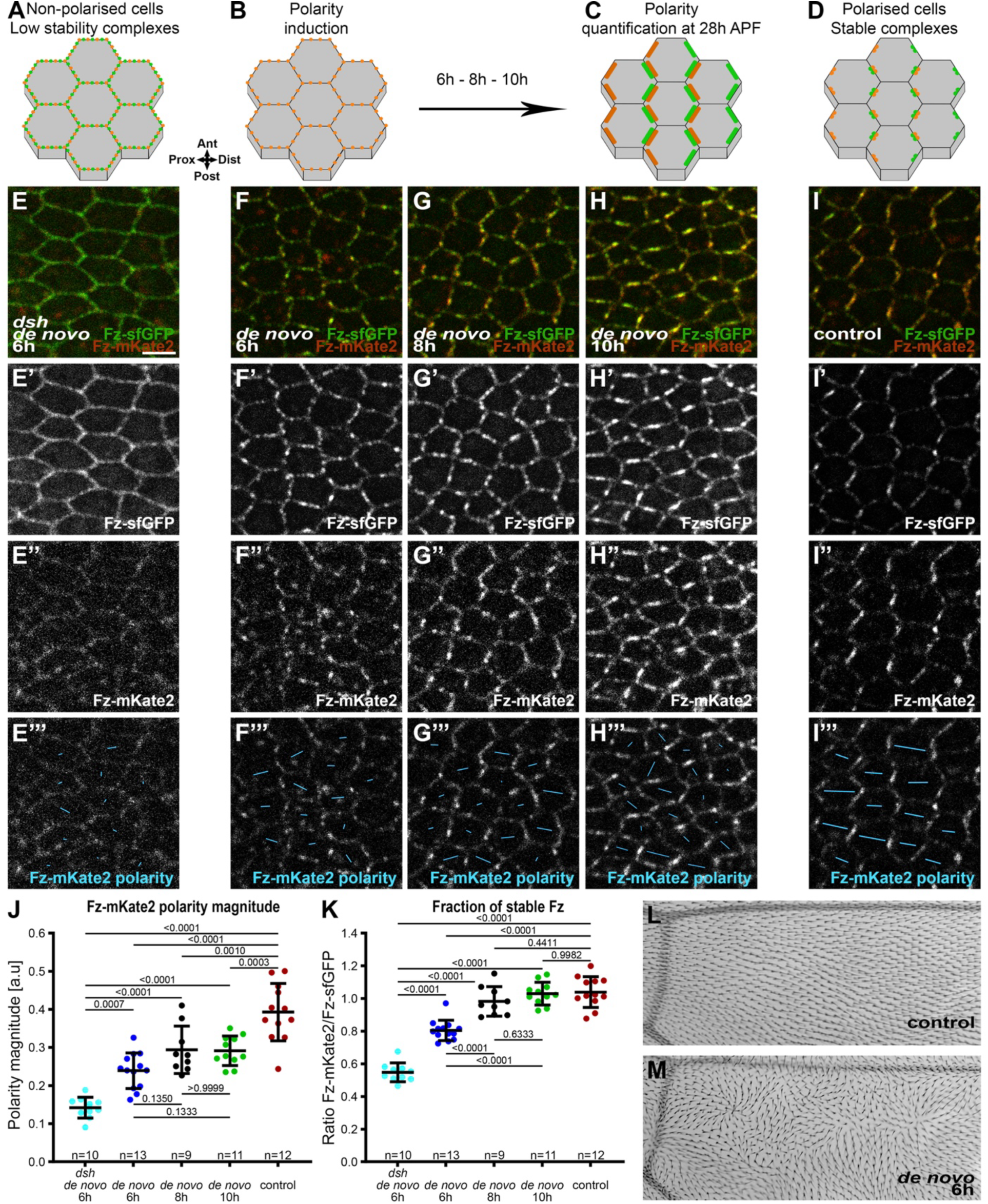
Dynamics of planar polarity establishment. **(A-D)** Schematics of localisation of core planar polarity pathway protein complexes in cells of the *Drosophila* pupal wing at 28h APF, in non-polarised cells **(A)**, before polarity induction **(B)**, after polarity induction (6h, 8h or 10h) **(C)**, or in the control condition with constitutive expression of Fz::mKate2-sfGFP **(D),** with localisation of Stbm (orange) and Fz (green) at apico-lateral cell junctions indicated. See Fig.**S1B** for induction timings. **(E-I)** Live confocal images of 28h APF pupal wing epithelia expressing Fz::mKate2-sfGFP taken below longitudinal vein 4, see **Fig.1C’** right-hand box. See **Table S1** for full genotypes. Scale bar, 4 µm and the same hereafter. Native fluorescence for sfGFP (green, **E’-I’**) and mKate2 (red, **E’’-I’’**), with *de novo* condition with 6h induction in a *dsh^1^* mutant background **(E)**, with *de novo* condition with induction for 6h **(F)**, 8h **(G)**, 10h **(H)** to establish core protein planar polarity; and in the control condition with constitutive expression of Fz::mKate2-sfGFP which gives a normal core protein proximo-distal polarity **(I)**. See **Fig.S1A** for details of Fz tagged to form a fluorescent timer. **(E’’’-I’’’)** Cell-by-cell polarity pattern of mKate2 fluorescence in pupal wings expressing Fz::mKate2-sfGFP at 28h APF. The length and orientation of cyan bars denote the polarity magnitude and angle respectively for a given cell. **(J, K)** Quantified polarity magnitude based on mKate2 fluorescence **(J)** or fraction of stable Fz as determined by ratio of mKate2/sfGFP fluorescence **(K)**, in live pupal wings at 28h APF, in the conditions described in **(E-I)**. Error bars are standard deviation (SD); n, number of wings. ANOVA with Tukey’s multiple comparisons test was used to compare all genotypes, p values as indicated. **(L, M)** Images of dorsal surface of mounted adult *Drosophila* wings, below longitudinal vein 4. **(L)** Control condition of constitutive Fz::mKate2-sfGFP expression, showing uniform distal orientation of trichomes. **(M)** *de novo* condition after induction at 22h APF, at the equivalent time to the 6h induction when imaging at 28h APF, showing swirled trichome pattern. Proximal is left, and anterior is up.

As an unpolarised condition we used flies deficient for *dsh* planar polarity activity, in which Fz::mKate2-sfGFP was induced *de novo* for 6h in a *fz* null mutant background (polarity induction at 22h after prepupa formation [APF] and polarity quantification at 28h APF). We characterised this situation as the lowest polarity condition for our study and defined it as ‘non-polarised’ cells and as having low stability core protein complexes (Fig.2A,J,K) consistent with the requirement for Dsh activity for Fz polarisation and stability (Strutt, 2001; Strutt et al., 2011; Warrington et al., 2017). In this context the specific zig-zag core protein localisation pattern was lost (Fig.2E’-E’’) and Fz polarity as measured by the PCA method was low (Fig.2E’’,J).

We then analysed *de novo* polarity induction in a wild-type background, where expression of Fz::mKate2-sfGFP was induced in a *fz* null background at 6h, 8h or 10h prior to observation at 28h APF (Fig.2B,F-H). Induction of Fz::mKate2 for 6h revealed detectable planar polarity compared to unpolarised cells (*dsh de novo* 6h vs *de novo* 6h; p=0.0007) (Fig.2J). This polarity increased with induction for 8h and 10h to reach a plateau (Fig.2J), with a lower peak value than in the control (constitutive Fz::mKate2-sfGFP) condition (*de novo* 10h vs control; p=0.0003). These three *de novo* polarity induction conditions gave similar swirling patterns of polarity (Fig.2F’’’,G’’’,H’’’,M), as expected for pathway activation in the absence of proximo-distal global cues (Strutt and Strutt, 2002; Strutt and Strutt, 2007). See Fig.S5B-E’’’ for further evidence that *de novo* induction polarity patterns are also not influenced by global cues in the proximal wing.

Quantification of the stable fraction of Fz (mKate2/sfGFP ratio) revealed that core protein complexes from 6h after polarity induction are more stable than those in unpolarised cells (*dsh de novo* 6h vs *de novo* 6hr; p<0.0001), reaching a similar level of stability as the normal condition 8h after polarity induction (*de novo* 8h vs control; p=0.4411) (Fig.2K). In summary, our findings reveal that following *de novo* polarity induction in the pupal wing, planar polarity magnitude and complex stability reach a plateau by 8h.

### Planar polarity establishment is not highly sensitive to variation in levels of core complex components

After characterisation of *de novo* polarisation induction in our system, we went on to ask whether *de novo* polarisation is sensitive to levels of individual core proteins. Based on theoretical considerations suggesting that cell polarisation depends on a cell-scale inhibitory signal (Meinhardt, 2007), we hypothesised that the levels of an individual core protein might be limiting for polarisation. This could be due either to levels being limiting for complex formation – thus placing an upper bound on the number of complexes that can form at cell junctions (representing a ‘depletion model’) (Fig.1B) – or due to a dosage sensitive role in inhibitory interactions that sort proteins within the cell (Fig.1A’,B’’).

We therefore looked at the effects of reducing gene dosage by 50% (i.e. the heterozygous condition) during Fz *de novo* polarisation (6h after induction) and once Fz *de novo* polarisation reaches a plateau (8h after induction), and also compared to polarisation in otherwise normal conditions. For *fmi*, *stbm*, *dgo* and *pk* we used molecularly characterised ‘null’ alleles (see Materials and Methods). In the case of *dsh* we used the planar polarity specific *dsh^1^* allele to avoid any potential confounding effects of compromising canonical Wg signalling due to Dsh functioning in both pathways (Axelrod et al., 1998; Boutros et al., 1998). In *dsh^1^*mutant tissue, Dsh protein is not seen localised to core protein complexes at cell junctions (Axelrod, 2001; Shimada et al., 2001) and quantitation of protein distributions supports *dsh^1^* being null for planar polarity function (Warrington et al., 2017). We have previously shown that the level of Stbm is reduced in the heterozygous condition (Strutt et al., 2016). We confirmed that the same was true for Fmi, Pk and Dgo by carrying out western blot analysis (Fig.S2A-D). The *dsh^1^* allele gives rise to a non-functional protein (Axelrod, 2001), however our previous western blot analysis supports Dsh levels being sensitive to gene dosage (Strutt et al., 2016).

As we measured Fz stability and polarity using our inducible Fz::mKate2-sfGFP expressing transgene, we were unable to vary Fz dosage in these experiments. Nevertheless, we previously showed that halving *fz* gene dosage did not significantly affect the overall levels of Fz protein (Strutt et al., 2016), consistent with Fz production not normally being limiting for core pathway function. Moreover, our western blot analysis indicates that levels of Fz::mKate2-sfGFP are several fold lower than endogenous Fz levels (Fig.S1C,D), suggesting that our *de novo* induction conditions most likely represent a ‘sensitised’ condition for analysing core pathway function.

Notably, we found no significant difference in the degree of polarisation (polarity magnitude) of Fz in polarity complexes at either 6h (Fig.3A-H) or 8h (Fig.S3A-H) after induction, when halving the dosage of *dgo*, *pk*, *dsh*, *stbm* or *fmi*. Halving the dosage of *pk*, *stbm* or *fmi* also had no significant effect on the stable fraction of Fz in polarity complexes at either 6h (Fig. 3J) or 8h (Fig.S3I), although a weak decrease was seen in Fz stability in *dsh^1^/+* at 6h and 8h after polarity induction and in *dgo/+* at 8h after polarity induction (Fig.3J,S3I). This is consistent with the known roles of Dsh and Dgo in stabilising Fz at cell junctions (Strutt et al., 2011; Warrington et al., 2017). Wings homozygous mutant for *dsh* showed the expected low polarity (*dsh/dsh de novo* 6h vs *de novo* 6h; p<0.0001) (Fig.3H) and low stability (*dsh/dsh de novo* 6h vs *de novo* 6h; p<0.0001) (Fig.3J). Interestingly, we did see an effect of core protein gene dosage on the magnitude of Fz polarity achieved (Fig.3I), but not on Fz stability (Fig.3K) in the control condition of constitutive Fz::mKate2-sfGFP expression. Specifically, halving the gene dosage of *dgo*, *pk*, *dsh* and *stbm* resulted in a reduction in polarity magnitude. This is in contrast to previous results when an effect on asymmetry of endogenous Fmi was not detected after halving gene dosages (Strutt et al., 2016). We surmise that this is due to levels of Fz::mKate2-sfGFP expressed constitutively under the *Actin5C* promoter being lower than endogenous Fz levels (Fig.S1C,D), resulting in enhanced sensitivity to reductions in levels of other core proteins.

**Figure 3.**
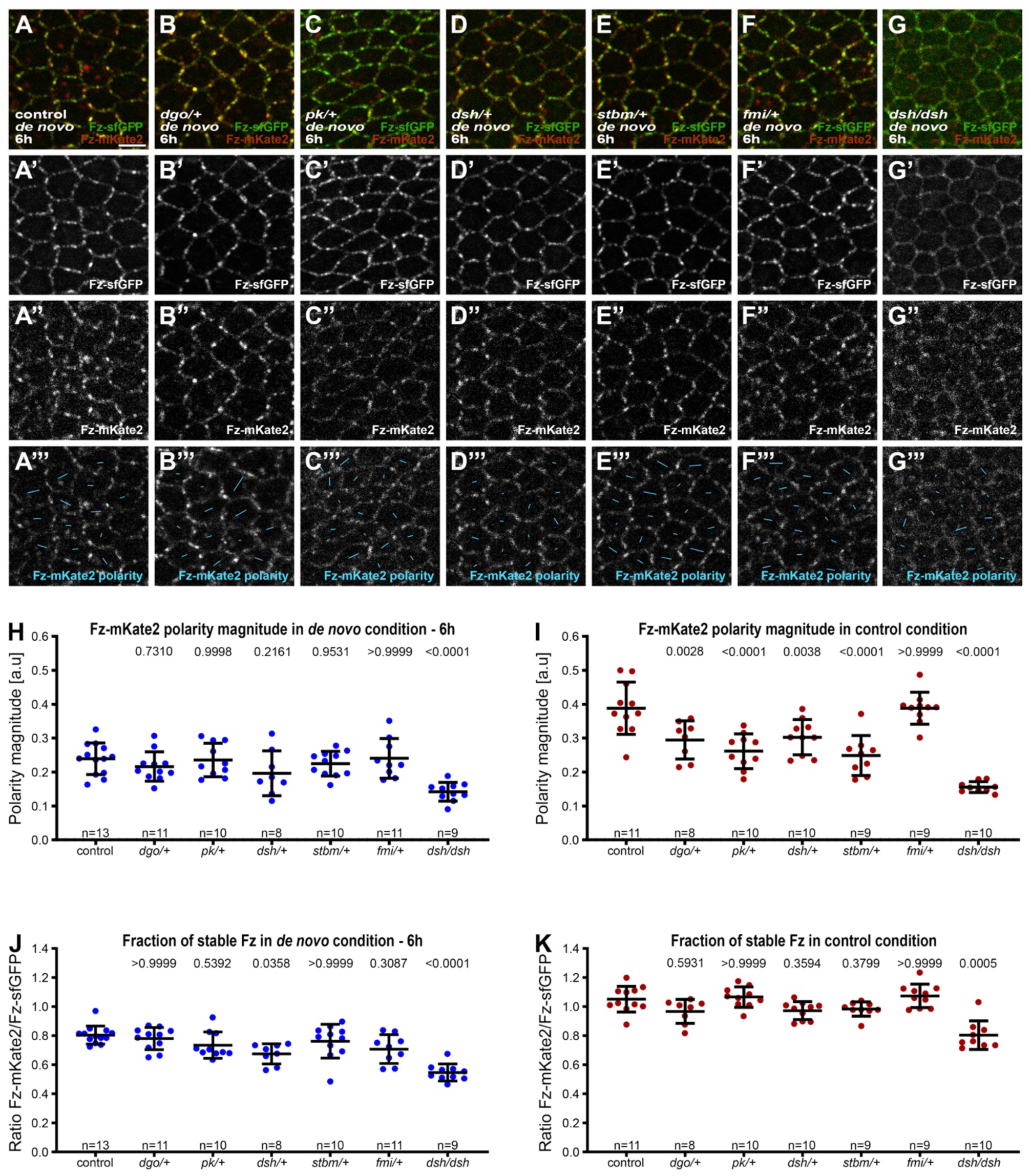
Effect of core protein dosage during *de novo* planar polarity establishment. **(A-G)** Live confocal images of 28h APF pupal wing epithelia expressing Fz::mKate2-sfGFP, in *de novo* condition taken below longitudinal vein 4. Native fluorescence for sfGFP (green, **A’-G’**) and mKate2 (red, **A’’-G’’**), with *de novo* condition with 6h induction to establish core protein planar polarity in an otherwise wild-type background **(A)**, heterozygous mutant backgrounds for *dgo* **(B)**, *pk* **(C)**, *dsh^1^* **(D)**, *stbm* **(E)**, *fmi* **(F)** and hemizygous mutant background for *dsh^1^* **(G)**. See **Table S1** for the full genotypes. See **Fig.S3** for live confocal images of equivalent 28h APF pupal wing epithelia with constitutive Fz::mKate2-sfGFP expression. **(A’’’-G’’’)** Cell-by-cell polarity pattern of mKate2 fluorescence in pupal wings expressing Fz::mKate2-sfGFP at 28h APF. The length and orientation of cyan bars denote the polarity magnitude and angle for a given cell respectively. **(H-K)** Quantified Fz-mKate2 polarity magnitude based on mKate2 fluorescence **(H, I)** or fraction of stable Fz as determined by ratio of mKate2/sfGFP **(J, K)** in live pupal wings at 28h APF, in the conditions described in **(A-G**). Error bars are SD; n, number of wings. ANOVA with Dunnett’s multiple comparisons tests for quantified Fz-mKate2 polarity and ANOVA with Kruskal-Wallis multiple comparisons for fraction of stable Fz, were used to compare wild-type to mutant backgrounds, p values as indicated. See also **Fig.S3**.

Overall, our results show that reductions in core protein dosage do not compromise the ability to establish planar polarity during the initial ‘fast’ phase of *de novo* polarisation (up to 8-10h, Fig.2J), despite the lower than normal Fz levels. Fz stability shows a weak sensitivity to levels of functional Dsh and Dgo during early phases of polarisation, but this is not sufficient to affect levels of polarity achieved. However, in wings continuously expressing Fz::mKate2-sfGFP, we do see sensitivity to Dgo, Pk, Dsh and Stbm levels, consistent with mature polarity being limited when protein levels are reduced.

### Microtubules do not provide a cell-scale signal for core protein localisation

Our results so far fail to provide evidence that the establishment of ‘cell-scale’ polarity can be explained by depletion of a limiting pool of a core protein. We therefore decided to explore another possibility, which is that core protein polarity might depend on polarised microtubule (MT) transport (Fig.1B’). A role for MTs in mediating an upstream ‘global cue’ via polarised transport or core proteins has been previously investigated (e.g. Shimada et al., 2006; Harumoto et al., 2010; Matis et al., 2014), but direct functions in core pathway cell-scale polarisation have not been studied. Loss of core protein function has been reported to not alter proximo-distal MT polarity during stages of pupal wing development when core polarity is normally proximo-distal (Harumoto et al., 2010), however, this does not rule out the possibility that core proteins help to orient MTs, possibly in parallel to other cues such as cell shape. Such core protein-dependent MT orientation could then amplify cell-scale core protein polarity through polarised core protein trafficking.

To investigate, we used our system of establishing *de novo* core protein polarity (induction at 22h APF and observation at 28h APF, in this case expressing Fz-eYFP under control of the *Actin5C* promoter), where core proteins polarise in swirling patterns (Fig.2F,M). To follow core protein and MT polarity and cell shape, we fixed and immunolabelled wings and acquired deconvolved images from a flat region of the wing below longitudinal vein 4 (Fig.1C’). In wild-type wings, cells were elongated (cell eccentricity close to 0.6) (Fig.4A, Fig.S4B) and were aligned following the proximo-distal axis (cell orientation close to 0°) (Fig.4D, Fig.S4A). In these cells, both endogenous Stbm protein and MTs were polarised following the same proximo-distal pattern (Fig.4A’’-A’’’’,G,J, Fig.S4C,E). Without *de novo* polarity induction (non-induced Fz-eYFP in a *fz* mutant background), there was no change in cell orientation and eccentricity compared to the wild-type condition (Fig.4B,E, Fig.S4A,B). However, Stbm was located all around the cell perimeter (Fig.4B’’,B’’’’,H, Fig.S4C,D), while MTs were still aligned following the proximo-distal axis (Fig.4B’’’K, Fig.S4E).

**Figure 4.**
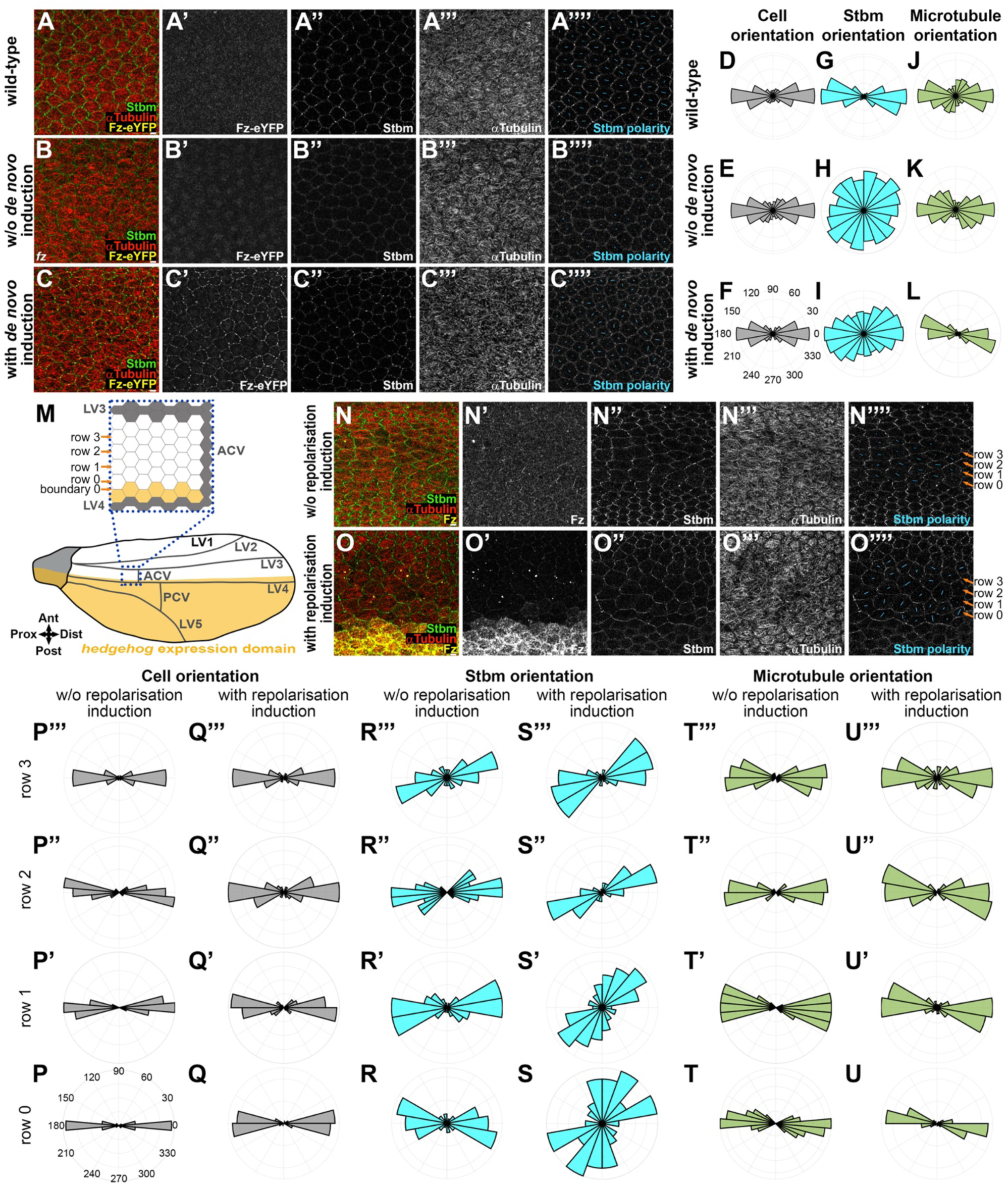
Apical microtubules and core proteins are oriented independently of each other. **(A-C)** Apical view of cells with deconvolved confocal images of 28h APF fixed pupal wing epithelia, distal to the posterior cross vein, below longitudinal vein 4, in a wild-type background **(A)**, in *de novo* Fz polarity establishment condition in a *fz* mutant background without polarity induction **(B)** and with polarity induction for 6h **(C)**. Native Fz-eYFP fluorescence (yellow, **A’**, **B’** and **C’**), immunolabelling for Stbm (green, **A’’**, **B’’** and **C’’)** and ⍺Tubulin (red, **A’’’**, **B’’’** and **C’’’)**. Note there is no Fz-eYFP fluorescence at cell junctions detected in (**A’** and **B’**). Cell-by-cell polarity pattern of pupal wings in respective conditions **(A’’’’**, **B’’’’** and **C’’’’**). The length and orientation of cyan bars denote the polarity magnitude and angle respectively for a given cell. See **Table S1** for the full genotypes. See **Fig.1C’** for vein locations and imaged wing area. **(D-L)** Polar histograms depicting binned cell orientation relative to the horizontal axis **(D-F)**, Stbm polarity orientation **(G-I)** and microtubule orientation **(J-L)** relative to the average cell orientation, in wild-type wings **(D**, **G**, and **J)**, in *de novo* condition without **(E**, **H**, and **K)** or with **(F**, **I**, and **L)** 6h Fz-eYFP polarity induction in fixed pupal wings at 28h APF. n=7-9 wings for each condition. See also **Fig.S4A-E**. **(M)** Cartoon of *hedgehog* expression domain (yellow) in the posterior part of the *Drosophila* pupal wing at 28h APF with a zoom in on the proximal region to the anterior cross vein between longitudinal vein 3 and longitudinal 4 (blue dash square). Grey cells are vein cells, white cells are intervein cells and yellow cells are intervein cells with *hedgehog* expression. Boundary 0 is between intervein cells expressing or not expressing *hedgehog* and the first row of intervein cells in contact with cells expressing *hedgehog* across boundary 0 is cell row 0 with anteriorly row 1, 2 and 3. **(N-O)** Apical view of cells from deconvolved confocal images of 28h APF fixed pupal wings, in the region proximal to the anterior cross vein between longitudinal vein 3 and longitudinal 4, in an otherwise wild-type background without Fz repolarisation induction **(N)** and with Fz repolarisation induction for 6h **(O)**. For Fz repolarisation conditions, induced Fz is over-expressed in the *hedgehog* expression domain (posterior wing part) with the UAS-GAL4 system (Fig.4M). Immunolabelling as described in **(A-C)**, against Fz (yellow, **N’**, **O’)**, Stbm (green, **N’’**, **O’’)** and ⍺Tubulin (red, **N’’’**, **O’’’)**. Cell-by-cell polarity pattern of pupal wings in respective conditions **(N’’’’**, **O’’’’**). The length and orientation of cyan bars denote the polarity magnitude and angle respectively for a given cell. See **Table S1** for the full genotypes. See Fig.4M and **Fig.1C’** for vein locations and imaged wing areas. **(P-U)** Polar histograms depicting binned cell orientation relative to the horizontal axis **(P-Q)**, Stbm polarity orientation **(R-S)** and microtubule orientation **(T-U)** relative to the average cell orientation, without **(P**, **R** and **T)** or with **(Q**, **S** and **U)** Fz repolarisation induction for 6h in fixed pupal wings at 28h APF. Cells are grouped in rows relative to their location relative to Fz overexpression (*hedgehog* expression domain) in Fz repolarisation condition or relative to longitudinal vein 4 without Fz repolarisation with row 0 in contact with the *hh-GAL4* overexpression boundary and row 3 furthest away. n=8 wings for both conditions. See also **Fig.S4F-J**.

Induction of *de novo* polarity did not change cell shape and orientation (Fig.4C,F, Fig.S4A,B) but induced a swirling core protein polarity with a partial proximo-distal bias in the studied region below vein 4 (Fig.4C’,C’’’’,I, Fig.S4C,D), in accordance with the orientation of swirled trichomes in the adult wing (Fig.2M). The difference in core protein localisation was not associated with a shift in MT orientation; MTs always followed a proximo-distal orientation (Fig.4C’’’,L, Fig.S4E).

To further explore the relationship between core protein polarisation and MT polarity, we next induced core protein repolarisation along the antero-posterior axis. We created an ectopic Fz boundary along horizontal cell membranes (Fig.4M) by over-expressing Fz in the *hedgehog* (*hh*) expression domain (posterior compartment) with an inducible UAS/GAL4 system (induction at 22h APF and observation at 28h APF) comprising *hh-GAL4* and a *UAS-FRT-STOP-FRT-fz* transgene in the presence of *hs-FLP*. We then analysed the region of the wing above this area of protein induction, in the region proximal to the anterior cross vein between longitudinal vein 3 and longitudinal vein 4 where we found that there is the strongest and sharpest boundary of Hedgehog expression (see Fig.4O’).

Without induction of core protein repolarisation, in this region of the wing endogenous Fz and Stbm were normally localised along medio-lateral cell junctions (Fig.4N-N’’), with a proximo-distal polarity orientation (Fig.4N’’’’,R-R’’’, Fig.S4H) as expected. Cells were also oriented along the proximo-distal axis (Fig.4P-P’’’, Fig.S4F) with MTs oriented in the same direction (Fig.4N’’’,T-T’’’, Fig.S4J).

After induction of core protein repolarisation, a Fz over-expression boundary was observable, defining the boundary 0 with cell row 0 (Fig.4M,O’). This repolarisation resulted in a relocalisation of Stbm along horizontal cell membranes (Fig.4O’’), leading to antero-posterior Stbm repolarisation over about 1-2 cell rows (Fig.4O’’’’,S-S’’, Fig.S4H) and an enrichment of Stbm along boundary 0 due to the Fz over-expression on the opposing boundary (Fig.4O’’, Fig.S4I). Despite the reorientation of polarity revealed by the change in Stbm localisation, MT orientation was unchanged, with a proximo-distal orientation in all cell rows (Fig.4O’’’,U-U’’’, Fig.S4J) and cells oriented along the same axis (Fig.4Q-Q’’’, Fig.S4F). We did observe an unexpected slight aspect change towards rounder cells in rows 2 and 3, which are away from the repolarisation boundary (Fig.S4G).

In summary, our results show that core pathway polarity can be established on an orthogonal (antero-posterior) axis with no corresponding change in MT polarity, consistent with the core proteins being able to segregate to opposite cell ends independently of MTs.

### Induced core protein relocalisation on a boundary does not lead to a wave of repolarisation

So far we have presented evidence against cell-scale polarity relying on a ‘depletion’ mechanism (i.e. not highly sensitive to core protein levels) or a MT transport mechanism. We have also observed that at the cellular level, over a period of 6h an antero-posterior boundary of Fz overexpression can only repolarise core protein localisation for about 1-2 cell rows (Fig.4O,S, Fig.S4H). This short range of repolarisation was unexpected, especially as the same induction regime results in repolarisation of trichomes in the adult wing over at least 5 rows (Fig.5C), consistent with previous reports (Wu and Mlodzik, 2008). To investigate further, we examined the sites of trichome emergence in pupal wings at 33h APF under similar induction conditions. With this longer induction time, we observed trichomes being repolarised for about 3-4 cell rows (Fig.S5A), but still less far than observed in adult wings (see Discussion).

**Figure 5.**
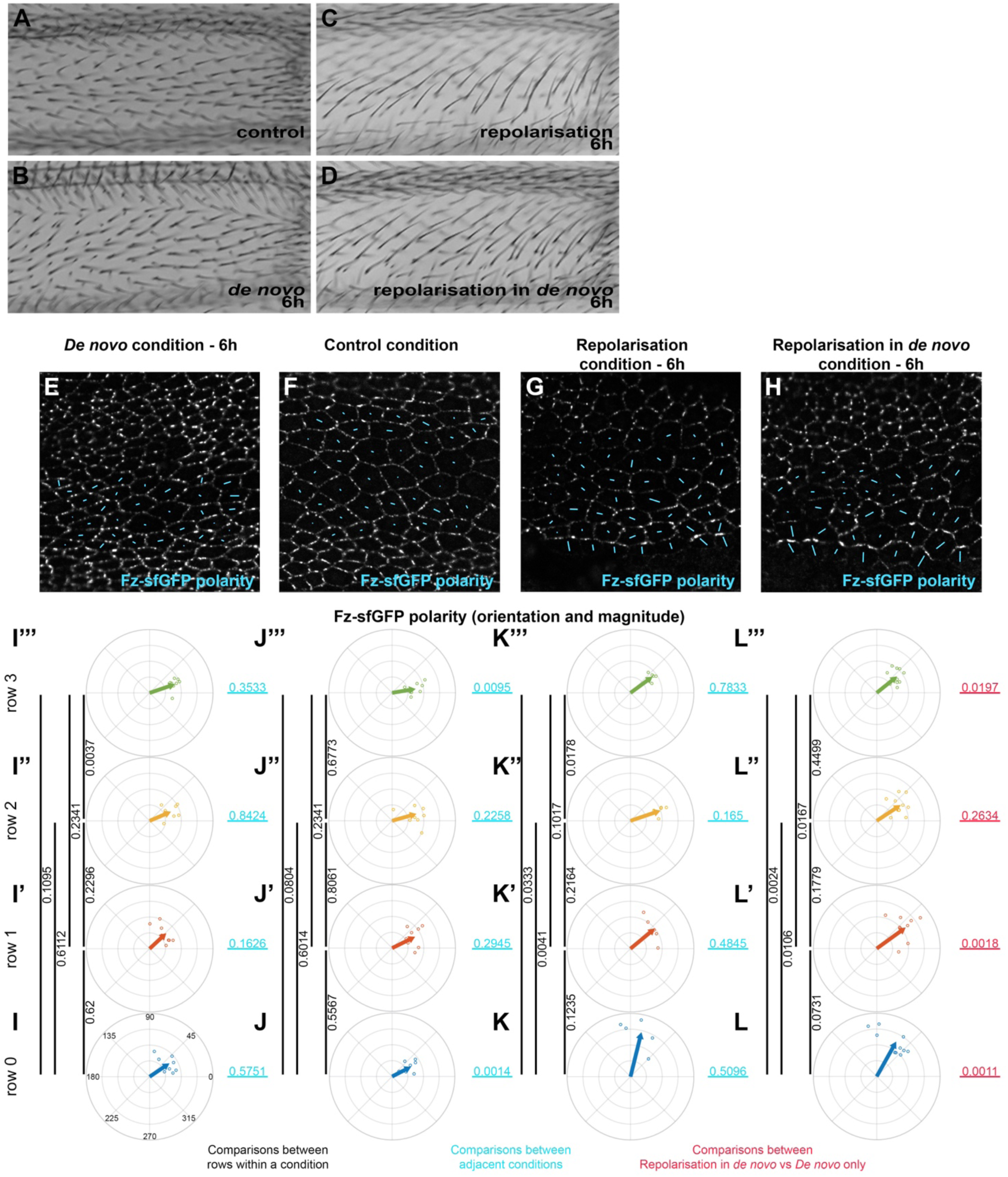
Reorientation of planar polarity from a boundary under control and *de novo* conditions. **(A-D)** Images of dorsal surface of adult *Drosophila* wings, taken proximal to the anterior cross vein between longitudinal vein 3 and longitudinal vein 4, showing trichome orientation in control condition with constitutive Fz::mKate2-sfGFP expression **(A)**, in *de novo* Fz::mKate2-sfGFP induction condition **(B)**, in repolarisation from the *hh-GAL4* boundary with constitutive Fz::mKate2-sfGFP expression condition **(C)**, and in repolarisation from the *hh-GAL4* boundary under *de novo* Fz::mKate2-sfGFP induction condition **(D)**. Repolarisation conditions are under UAS-GAL4 control for Fz over-expression in the posterior wing (*hedgehog* expression domain), with induced Fz expression from *UAS-FRT-STOP-FRT-Fz* in the presence of *hsFLP*. Note, the swirling pattern of polarity generated by *de novo* induction generates a proximo-distal trichome orientation in this analysed wing area **(B)** but not in surrounding wing regions **(Fig.S5B-D)**. Trichome polarity is reoriented in the antero-posterior direction in both repolarisation conditions. See Fig.4M and **Fig.1C’** for vein locations and imaged wing area. See **Table S1** for the full genotypes. **(E-H)** Planar polarity measurement at the cellular scale in the region proximal to the anterior cross vein, between longitudinal vein 3 and longitudinal vein 4 in fixed pupal wings at 28h APF. The length and orientation of cyan bars denote the polarity magnitude and angle for a given cell respectively. *De novo* condition with 6h induction of Fz::mKate2-sfGFP **(E)**, constitutive expression of Fz::mKate2-sfGFP **(F)**, repolarisation condition with constitutive expression of Fz::mKate2-sfGFP **(G)**, and repolarisation condition with 6h *de novo* induction of Fz::mKate2-sfGFP **(H)**. In repolarisation conditions, induced Fz is over-expressed in the posterior wing, whereas in anterior wing only Fz::mKate2-sfGFP is expressed. In the posterior wing the two Fz populations (over-expressed Fz and Fz::mKate2-sfGFP) compete for the same membrane locations and Fz::mKate2-sfGFP signal is not evident. **(I-L)** Circular plots of quantified total Fz (Fz::sfGFP) magnitude and orientation relative to horizontal axis in the region proximal to the anterior cross vein, between longitudinal vein 3 and longitudinal vein 4 at 28h APF in fixed pupal wings. Small dots show polarity angle and magnitude for individual wings, arrows show average polarity and magnitude across all wings. Cells are grouped in rows relative to their location relative to the Fz overexpression domain in Fz repolarisation condition or relative to longitudinal vein 4 without Fz repolarisation, with row 0 in contact with Fz overexpression boundary and row 3 furthest away. **(I-I’’’)** *de novo* condition with 6h to establish core protein polarity, **(J-J’’’)** control condition with constitutive Fz::mKate2-sfGFP expression, **(K-K’’’)** repolarisation condition for 6h to re-orient core protein polarity, and **(L-L’’’)** repolarisation under *de novo* condition for 6h induced Fz::mKate2-sfGFP expression); in row 0 **(I**, **J**, **K** and **L)**, in row 1 **(I’**, **J’**, **K’** and **L’)**, in row 2 **(I’’**, **J’’**, **K’’** and **L’’)** and in row 3 **(I’’’**, **J’’’**, **K’’’** and **L’’’)**. Vertical black lines associated with p values on right of each column represent comparisons between different rows of the same polarisation condition. Horizontal pale blue lines associated with p values represent comparison between the two adjacent polarisation conditions for the same cell row. On the far right, horizontal red underlined p values represent comparison between repolarisation in *de novo* condition versus *de novo* condition (far left) for the respective rows of cells. Hotelling’s T-square tests were used to compare total Fz polarity (orientation and magnitude).

We hypothesised that the limited ability of a Fz overexpression boundary to repolarise may be due to three factors. First, there might be a strong and persistent influence of a global cue specifying proximo-distally oriented polarity. Second, there might be strong cell-cell coupling of polarity between neighbours. Hence, a local perturbation, e.g. over-expression of Fz in a neighbouring row of cells, has little effect on the polarity of neighbouring cells, as their polarity is already strongly coupled to their neighbours in the adjacent region of non-overexpression. Third, cells might have a strong intrinsic ability to polarise, while cell-cell coupling of polarity with neighbours might be weak. In this case, once established, polarity is highly robust to the effects of Fz overexpression in neighbouring cells due to relatively weak effects of cell-cell coupling.

In terms of examining these possibilities, we have already attempted to find conditions that weaken (but do not break) the cell-intrinsic polarisation (Fig.3) by altering core protein levels, but this was unsuccessful. Moreover, there is no reported way to weaken cell-cell coupling of polarity. Interfering with global cues also presents a challenge, given that they are poorly characterised. However, current evidence suggests that *de novo* induction at times following the onset of hinge contraction results in a swirling polarity pattern that is not influenced by global cues (see Introduction), a possibility that we investigate further below.

Hence, we decided to compare repolarisation, induced as previously with *hh-GAL4*, in a control condition with constitutive Fz::mKate2-sfGFP expression (where global cues set up proximo-distal polarity), to repolarisation in the *de novo* condition produced by Fz::mKate2-sfGFP induction at 22h APF, where cells are in the process of polarising. Our prediction was that *de novo* polarity would be easy to orient/repolarise by a boundary of Fz overexpression, as cells (and their neighbours) should not have a pre-existing polarity.

We first looked at the polarity of trichomes in the adult wing. In the control condition of constitutive Fz::mKate2-sfGFP expression, proximal to the anterior cross vein between longitudinal vein 3 and longitudinal vein, as expected trichomes are oriented along the proximo-distal axis (Fig.5A). We found that in this wing region, trichomes also point proximo-distally after *de novo* polarisation induction of Fz::mKate2-sfGFP expression at 22h APF (Fig.5B). To investigate the possibility that this proximo-distal polarity in the *de novo* condition might be due to the presence of a previously unappreciated proximo-distal global cue, we examined the trichome polarity in surrounding regions of the wing. Notably, regardless of whether Fz::mKate2-sfGFP expression was induced at 18h, 20h or 22h APF, we observed a similar trichome swirling pattern in the proximal wing, with regions proximal, distal, anterior and posterior to the experimental region showing non-proximo-distal adult trichome polarity (Fig.S5B-D). Moreover, quantification of polarity at 28h APF in the experimental region showed no difference in the degree of proximo-distal polarisation with longer induction times (Fig.S5E). These observations argue against a proximo-distal global cue being active in this region of the wing at this developmental stage. Instead, we surmise that the proximo-distal polarity in the experimental region is part of the normal trichome swirling pattern in this wing region. Moreover, we can expect to see repolarisation of this proximo-distal polarity to antero-posterior polarity if we overexpress Fz in the posterior compartment.

Interestingly, induction of repolarisation in both the control and *de novo* conditions resulted in similar degrees of partial trichome repolarisation on the antero-posterior axis (Fig.5C,D). In neither case was trichome repolarisation seen beyond longitudinal vein 3. This result does not fit our prediction of longer-range repolarisation in *de novo*.

We then compared Fz distribution in this region of pupal wings at 28h APF in the same four conditions: control constitutive Fz::mKate2-sfGFP expression, 6h of *de novo* Fz::mKate2-sfGFP polarity establishment, 6h of repolarisation caused by Fz overexpression in the posterior compartment, and 6h of *de novo* polarity with 6h of repolarisation. As we saw in the adult wing, *de novo* polarity was broadly proximo-distal and similar in magnitude to the control polarity establishment in this region (Fig.5E,F,I-J’’’).

Surprisingly, in both the repolarisation (Fig.5G,K-K’’’) and repolarisation in *de novo* conditions (Fig.5H,L-L’’’), only row 0 was strongly repolarised (Fig.5J vs 5K p=0.0014, Fig.5L vs 5I p=0.0011). In the repolarisation only condition, there was no significant change in polarity in rows 1 or 2 (Fig.5J’ vs 5K’ p=0.2945, 5J’’ vs 5K’’ p=0.2258) although unexpectedly row 3 showed a significant displacement from proximo-distal polarity (Fig.5J’’’ vs 5K’’’ p=0.0095). In the repolarisation in *de novo* condition, row 1 was modestly repolarised towards the antero-posterior axis (Fig.5L’ vs Fig.5I’ p=0.0018), but again row 2 showed no detectable repolarisation (Fig.5L’’ vs Fig.5I’’ p=0.2634) and unexpectedly row 3 also showed antero-posterior displacement (Fig.5L’’’ vs Fig.5I’’’ p=0.0197).

The strong repolarisation of row 0 (in cells touching Fz overexpressing cells) is expected, as Stbm is known to be strongly recruited to cell boundaries with high apposing levels of Fz (Bastock et al., 2003). However, the variable pattern of weak repolarisation between rows 1-3 indicates a failure of this strong repolarisation to propagate from cell to cell, and we suspect the variation may be simply due to sampling noise. In particular, these data indicate that there is no dramatic increase in repolarisation from a boundary in *de novo* as compared to repolarisation in the control condition, with only a modest difference in polarity in row 1 in the *de novo* condition.

Overall, our results show that it is hard to repolarise from a boundary of Fz overexpression in both control and *de novo* polarity conditions, consistent with *de novo* polarity being rapidly and robustly established and not easily perturbed. This provides evidence in favour of an effective cell-intrinsic polarisation mechanism.

### Cell-intrinsic polarity is robust against local perturbation

Given the different conditions, the similarity of degree of repolarisation in control vs *de novo* conditions was puzzling. We reasoned that there might be differences in these conditions that were masked by simply using measures of overall cell polarity. We therefore carried out a more detailed analysis, measuring levels of proteins on individual cell junctions (Fig.6A).

**Figure 6.**
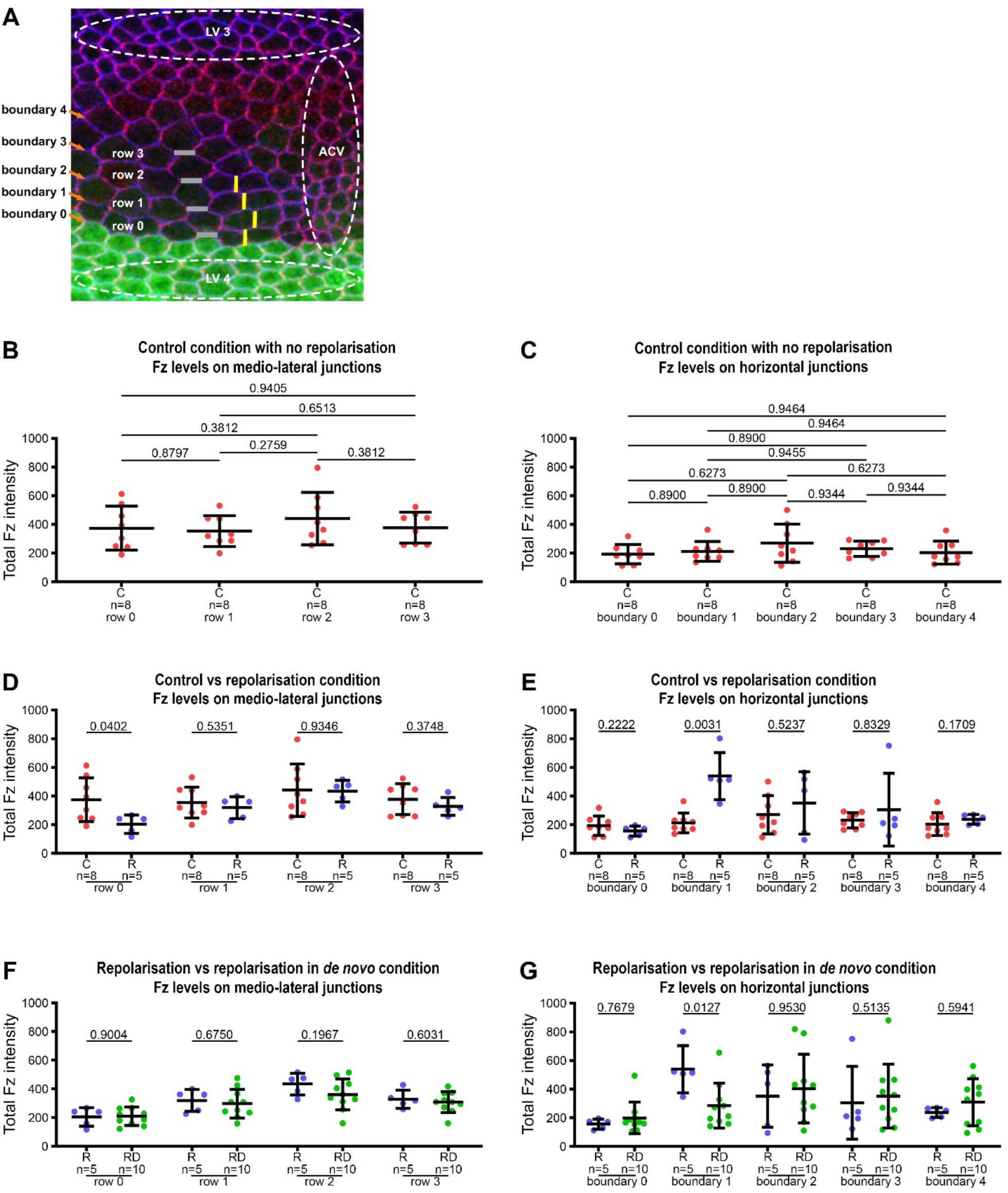
Distribution of Fz along medio-lateral and horizontal junctions in Fz repolarisation and repolarisation under *de novo* conditions. **(A)** Explanatory diagram for wing region, cell rows and cell boundaries. Visualisation of the anterior part of the anterior cross vein (ACV) between the longitudinal vein 3 (LV 3) and the longitudinal vein 4 (LV 4) in a fixed pupal wing at 28h APF. Green area indicates the expression domain of the *hh-GAL4* driver used to over-express Fz. E-Cadherin staining is in blue and Stbm staining in magenta. Junctions with horizontal orientation are schematised with grey lines, forming horizontal boundaries with boundary 0 between last row of cells over expressing Fz in LV 4 and the first row of cells without over expression of Fz in the intervein area (row 0). Medio-lateral junctions are schematised in yellow. As in Fig.5, repolarisation conditions are under UAS-GAL4 control for Fz over-expression in the posterior wing (*hedgehog* expression domain), with induced Fz expression from *UAS-FRT-STOP-FRT-Fz* in the presence of *hsFLP*. **(B-G)** Quantitation of Fz intensity in fixed pupal wings at 28h APF, binned by row of cells relative to their location relative to Fz overexpression in Fz repolarisation and Fz repolarisation under *de novo* conditions, or relative to longitudinal vein 4 without Fz repolarisation induction, with row 0 in contact with Fz overexpression boundary and row 3 furthest away. Boundary 0 is the posterior horizontal junction of row 0 in contact with Fz overexpression area and boundary 1 is the anterior horizontal junction shared between cells in row 0 and row 1. See **Table S1** for the full genotypes. Error bars are SD; n, number of wings, p values as indicated. **(B, C)** Total Fz intensity in control condition with constitutive Fz::mKate2-sfGFP expression, along medio-lateral junctions **(B)** and along horizontal junctions **(C)**. ANOVA with Holm-Sidak’s multiple comparisons test was used to compare all conditions. **(D, E)** Comparison of Fz distribution in control condition with constitutive Fz::mKate2-sfGFP expression versus repolarisation condition with 6h induction, along medio-lateral junctions **(D)** and along horizontal junctions **(E)**. Unpaired t test **(D)** and Mann-Whitney test **(E)** were used to compare fluorescence intensities. **(F, G)** Comparison of Fz distribution in repolarisation condition versus repolarisation in *de novo* condition with 6h induction, along medio-lateral junctions **(F)** and along horizontal junctions **(G)**. Unpaired t test **(F)** and Mann-Whitney test **(G)** were used to compare fluorescence intensities. See also **Fig.S6**.

In control conditions, as expected, all cells in rows 0 to 3 have similar protein levels on their medio-lateral junctions (Fig.6B, yellow lines in Fig.6A), and on their horizontal junctions (Fig.6C, grey lines in Fig.6A), with ∼2x higher levels on the medio-lateral junctions, consistent with the observed proximo-distal polarity.

After 6h repolarisation in the control condition, Fz distribution is only significantly altered on the row 0 medio-lateral junction (Fig.6D) and the boundary 1 horizontal junction between row 0 and row 1 (Fig.6E). Specifically, Fz is lost from the medio-lateral junctions of cells in row 0 and increased on the horizontal junctions along boundary 1. This is consistent with the changes in cell polarity seen in rows 0 and 1 (Fig.5G,K-K’’’), and supports the view that polarity changes only propagate 1-2 rows from a boundary of Fz overexpression.

We then investigated 6h repolarisation under *de novo* conditions. Interestingly, Fz levels along junctions in repolarisation in *de novo* do not exactly mirror those in repolarisation in the control condition. While Fz levels are similarly low on medio-lateral junctions in row 0 (Fig.6F), they are less high on the horizontal boundary 1 between rows 0 and 1 than in repolarisation in normal condition (Fig.6G). We questioned whether this difference might be due to the short period of polarisation, but obtained the same result after 10h induction (Fig.S6A-D).

Taken together, our findings reveal that induction of repolarisation from a boundary leads to a new core protein distribution along cell membranes, with relocalisation from medio-lateral junctions to horizontal junctions. However, these core protein relocalisations do not induce polarity rearrangement in neighbouring cells. This failure of repolarisation to propagate beyond 1-2 rows of cells is consistent with the presence of an effective cell-intrinsic polarisation machinery, which is only weakly affected by cell-cell coupling of polarity to neighbours.

## Discussion

The establishment of coordinated planar polarity across a tissue depends on processes acting at different scales (Devenport, 2014; Aw and Devenport, 2017; Butler and Wallingford, 2017; Harrison et al., 2020; Strutt and Strutt, 2021; Brittle et al., 2022). These include global cues at the tissue level, cell-intrinsic polarisation mechanisms that can amplify polarity either on cell-cell boundaries or at the level of whole cells, and cell-cell coupling of polarity between neighbouring cells (see Fig.1). An important question is the relative influence of each of these processes on the final polarity pattern.

In this work we have focused on understanding how cell-intrinsic ‘cell-scale signalling’ might occur in the *Drosophila* pupal wing, promoting segregation of the core proteins to opposite cell ends. We used several tools including: (i) *de novo* induction of core pathway planar polarity, to bypass earlier effects of global cues; (ii) manipulation of core protein gene dosage to test ‘depletion’ models for cell-scale patterning; (iii) induced repolarisation of planar polarity from a boundary to challenge the effects of cell-intrinsic polarisation mechanisms; and (iv) use of a fluorescent timer to monitor Fz stability simultaneously at multiple cell junctions over time.

Based on experiments in which we attempted to repolarise tissue from a boundary of Fz overexpression, we find support for the existence of a robust cell-scale signalling mechanism that is able to establish and maintain polarity in the face of local perturbations. However, our results fail to provide evidence that such cell-scale signalling is mediated by either a depletion of a limiting pool of an individual core protein, or by reorientation of the apical MT network by core pathway activity.

While we have yet to elucidate the underlying mechanism of cell-scale signalling, we believe that such signalling underpins the reported ability of both insect and mammalian epithelial cells to spontaneously planar polarise with only locally coordinated polarity that does not align with a global tissue axis (Strutt and Strutt, 2002; Strutt and Strutt, 2007; Vladar et al., 2012; Aw et al., 2016). Moreover, the conservation across species suggests that there may be a single universal mechanism. As discussed in previous work (Meinhardt, 2007; Burak and Shraiman, 2009; Abley et al., 2013; Shadkhoo and Mani, 2019), such a signal may be mediated by diffusion of a cytoplasmic protein (Fig.1B’’) and might be generated by (for instance) post-translation modification of a core complex component. In this context, it is interesting to note that multiple post-translational modifications of core proteins have been reported (reviewed in Harrison et al., 2020), which represent potential candidate mechanisms.

A caveat to our conclusion that cell-scale signalling does not depend on depletion of a limited pool of one of the core proteins is that we only lowered protein levels by 50%. However, we believe that such conditions would be sufficient to reveal a role for two reasons. First, we assayed the rate of *de novo* polarisation, rather than simply the final polarity achieved. Our reasoning was that if a core protein were limiting, halving the levels would minimally be expected to reduce the rate of the polarisation process. Secondly, the experiments were carried out under conditions where Fz levels were several fold lower than normal, which we would expect to further sensitise the system if one of the other core proteins were limiting. However, we note that we cannot eliminate the possibility that Fz itself is the limiting factor.

Interestingly, repolarisation from a boundary in the control and *de novo* conditions exhibited different effects on Fz distribution. In the control condition (constitutive expression of Fz::mKate2-sfGFP), repolarisation in row 0 occurs via loss of Fz from medio-lateral junctions and accumulation of Fz on the horizontal junction away from the region of overexpression (Fig.7B). This could be either a redistribution of existing protein via endocytosis and trafficking, or removal and degradation of protein from medio-lateral junctions in parallel with delivery of new protein to the horizontal junction (boundary 1), as previously suggested (Butler and Wallingford, 2017; Warrington et al., 2017). These results are consistent with a cell-scale signal possibly mediated by ectopic recruitment of Stbm to boundary 0, that promotes Fz removal from medio-lateral junctions and localisation to the opposite horizontal junction. In contrast, under *de novo* conditions, a Fz expression boundary results in repolarisation in row 0 that is characterised by low Fz on medio-lateral boundaries, but no strong Fz accumulation on boundary 1 (Fig.7C). Failure to see high Fz on any of the boundaries in row 0 was unexpected, but again supports the presence of a cell-scale signal coming from boundary 0 preventing Fz from accumulating on medio-lateral junctions.

**Figure 7.**
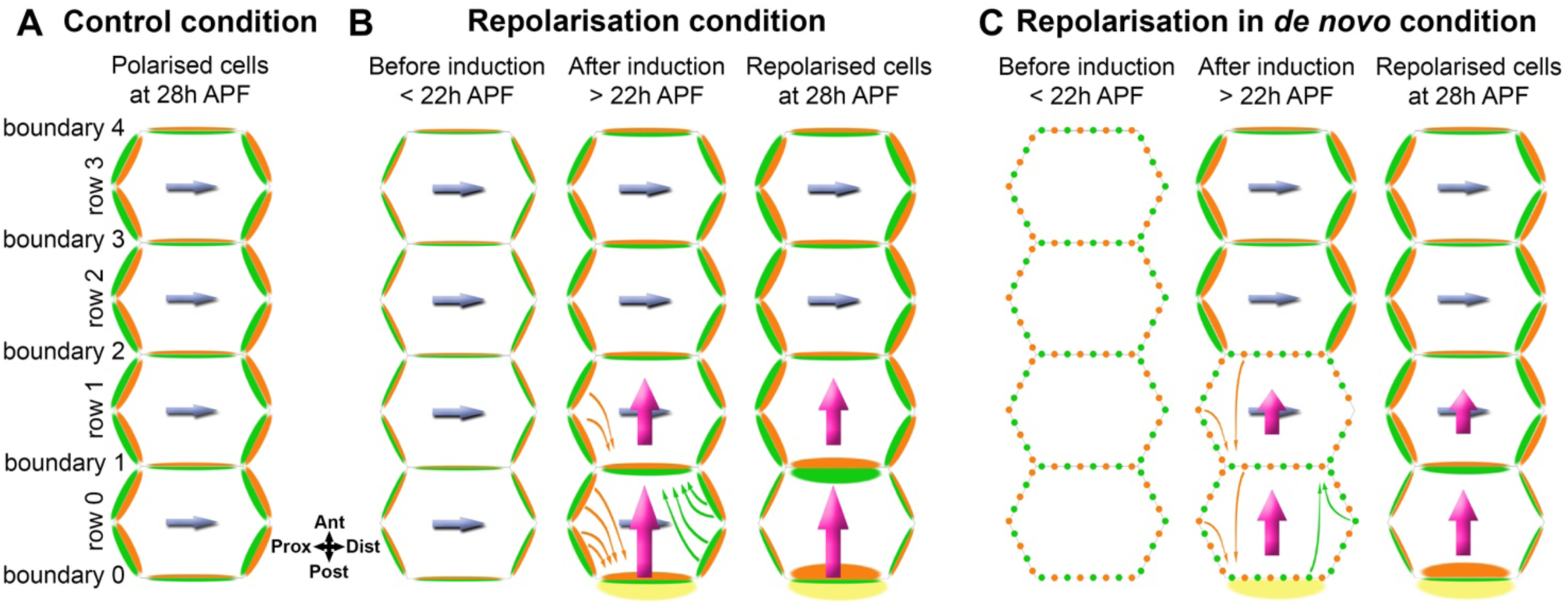
Summary model of polarity propagation confronted with cell-scale polarity. **(A)** Core protein polarisation in control (wild-type) conditions at 28h APF. The level of total Fz (green) does not vary significantly on different medio-lateral junctions or on different horizontal junctions. Stbm (orange) levels mirror Fz levels on the same cell junctions. Planar polarity is directed by the integrated effects of an upstream extrinsic global cue and a cell intrinsic cell-scale polarisation system (blue arrow) with a proximo-distal orientation. **(B)** Repolarisation of core protein polarity from a boundary. Before repolarisation induction, Fz (green) and Stbm (orange) become polarised following the proximo-distal axis due to the normal extrinsic and intrinsic planar polarity systems (blue arrow). After repolarisation induction by Fz over-expression in the posterior part of the wing, over-expressed Fz (yellow) accumulates on posterior boundary 0. In row 0, Fz is seen to be lost from medio-lateral junctions and to accumulate on the horizontal junction away from the region of overexpression (boundary 1). This could be due to redistribution of existing protein via endocytosis and trafficking (green arrows) or removal and degradation of protein from medio-lateral junctions and net delivery of new protein to horizontal junctions. Stbm redistribution also occurs (orange arrows) with a loss of Stbm from medio-lateral junctions and accumulation on anterior boundary 0. We surmise that Stbm redistribution to boundary 0 is primarily a result of Stbm being sequestered by forming asymmetric complexes with the high levels of over-expressed Fz on the other side of the boundary. In row 1, it is likely that a loss of Stbm located on medio-lateral junctions, and relocation along boundary 1, again occurs via the same potential mechanisms with sequestration into boundary 1 due to accumulation of Fz on the other side of this boundary. We hypothesise that Stbm redistribution to boundaries 0 and 1 results in overall cell repolarisation of rows 0 and 1 through generation of a cell-scale cell intrinsic polarity signal (magenta arrows) that overrides the original cell polarity cues (blue arrows). Notably, repolarisation does not propagate into row 2, presumably due to weak cell-cell coupling of polarity and the strong cell-intrinsic polarity mechanisms in these cells (blue arrows). **(C)** Repolarisation of core protein polarity from a boundary in *de novo* conditions. As in repolarisation of core protein polarity in the control conditions, we surmise that Stbm accumulates on boundary 0 in row 0, due to high Fz levels on the other side of the boundary causing Stbm to be sequestered into asymmetric complexes. This could then result in a cell-scale polarity signal (magenta arrows) on the antero-posterior axis that promotes Fz accumulation on boundary 1 in row 0, rather than on mediolateral boundaries. Concurrently, cells in rows 1-3 spontaneously polarise and adopt a proximo-distal polarity due to the presence of a strong cell intrinsic polarisation system (blue arrows). Weak cell-cell coupling of polarity means that the altered polarity in row 0 does not propagate significantly.

An unexplained observation is that in our repolarisation from a boundary experiments, when observing trichome initiation in the 33h pupal wing we only detect significant repolarisation over about 3-4 cell rows, but in the adult wing repolarisation extends at least 5 cell rows. We propose that there may exist some degree of coupling of polarity between mature trichomes that is not yet present when pre-hairs emerge at the cell edge (Wong and Adler, 1993), which might be mechanical in nature, or be mediated for instance by septate junction proteins (Venema et al., 2004).

In conclusion, this work represents the first attempt to systematically study cell-scale signalling in core pathway planar polarity. We provide evidence for and against different possible mechanisms, providing the basis for future work.

## Acknowledgments

Larra Trinidad, Samantha Warrington and Carl Harrison are thanked for useful discussions and the fly room staff for excellent technical support. We are grateful to Natalia Bulgakova and Miguel Ramirez-Morena for advice on microtubule angle quantification. Imaging was performed in the Wolfson Light Microscopy Facility. The work was funded by a Wellcome Trust Senior Fellowship (210630/Z/18/Z) to DS.

## Author contributions

Conceptualisation, A.C. and D.S.; methodology, A.C. and D.S.; formal analysis, A.C.; investigation, A.C. and H.S.; resources, A.C. and D.S.; writing – original draft, A.C. and D.S.; writing – review & editing, A.C., H.S. and D.S.; supervision, D.S.; funding acquisition, D.S.

## Materials and Methods

### Resources Table

| Antibody | Source | Identifier |
| --- | --- | --- |
| For immunolabelling: |  |  |
| Mouse monoclonal anti-α-Tubulin | Sigma | T9026 |
| Rabbit anti-Stbm DS17 | Ref (Warrington et al., 2013) | N/A |
| Rat anti-Stbm DS18 | Ref (Strutt and Strutt, 2008) | N/A |
| Rabbit anti-Fz | Ref (Bastock and Strutt, 2007) | N/A |
| Rabbit anti-GFP | Abcam | Ab6556 |
| For western blotting: |  |  |
| Mouse monoclonal anti-α-Tubulin | Sigma | T9026 |
| Mouse monoclonal anti-Fmi 74 | Ref (Usui et al., 1999) | DSHB |
| Rabbit anti-Fz | Ref (Bastock and Strutt, 2007) | N/A |
| Rabbit anti-GFP | Abcam | Ab6556 |
| Rat anti-Pk | Ref (Strutt et al., 2013) | N/A |

| Fly | Source | Identifier |
| --- | --- | --- |
| *stbm*<sup>6</sup> | Ref (Wolff and Rubin, 1998) | FlyBase: FBal0062423 |
| *dsh*<sup>1</sup> | Ref (Perrimon and Mahowald, 1987) | FlyBase: FBal0003138 |
| *dgo*<sup>380</sup> | Ref (Feiguin et al., 2001) | FlyBase: FBal0141190 |
| *pk*<sup>pk-sple13</sup> | Ref (Gubb et al., 1999) | N/A |
| *fmi*<sup>E59</sup> | Ref (Usui et al., 1999) | FlyBase: FBal0101421 |
| *fz*<sup>P21</sup> | Ref (Jones et al., 1996) | FlyBase: FBal0004937 |
| *P[w+, Actin5C-FRT-PolyA-FRT-fz-eYFP]* | Ref (Strutt, 2001) | FlyBase: FBtp0017633 |
| *ActP-FRT-polyA-FRT-fz-mKate2-sfGFP* | Ref (Ressurreicao et al., 2018) | FlyBase: FBal0361851 |
| *P[w+, Act>STOP>fz-mKate2-sfGFP]* | Ref (Ressurreicao et al., 2018) | FlyBase: FBal0361851 |
| *hsFLP[attP2]* | Bloomington Drosophila Stock Center | FlyBase: FBti0160508 |
| *hsFLP1* | Bloomington Drosophila Stock Center | FlyBase: FBti0002044 |
| *hsFLP22* | Bloomington Drosophila Stock Center | FlyBase: FBti0000785 |
| *P[w+, hh-GAL4]* | Ref (Tanimoto et al., 2000) | FlyBase: FBal0121962 |
| *P[w+, UAS>STOP>FzNL]* | This work (on III) |  |
| *w*<sup>1118</sup> | Bloomington Drosophila Stock Center | FlyBase: FBal0018186 |
| *P[acman]-EGFP-Dgo* | Ref (Ressurreicao et al., 2018) | FlyBase: FBrf0237330 |
| Chemical | Source | Identifier |
| 16% paraformaldehyde solution (methanol free) | Agar Scientific | cat#R1026 |
| Triton X-100 | VWR | cat#28817.295; CAS: 9002-93-1 |
| Tween-20 | VWR | cat#437082Q; CAS: 9005-64-5 |
| Glycerol | VWR | cat#284546F; CAS: 56-81-5 |
| DABCO | Fluka | cat#33480; CAS: 280-57-9 |
| Vectashield | Vector Labs | cat#H-1000-10 |
| Normal Goat Serum | Jackson Labs | cat#005-000-121 |
| Alexa568-Phalloidin | Thermo | cat#A12380 |

| Software |  |
| --- | --- |
| ImageJ 1,54f | https://imagej.nih.gov/ij/<br>Ref (Schneider et al., 2012) |
| MatLab R2019b | https://uk.mathworks.com/products/matlab.html |
| Puncta selection script (MatLab) | Ref (Strutt et al., 2016) |
| Microtubules orientation script (MatLab) | Ref (Ramirez-Moreno et al., 2023) |
| GraphPad Prism v9 | https://www.graphpad.com |
| QuantifyPolarity | Ref (Tan et al., 2021) |
| NIS Elements AR 4.60 | Nikon |
| Tissue Analyzer | https://grr.gred-clermont.fr/labmirouse/software/WebPA/<br>Ref (Etournay et al., 2016) |

### Fly genetics and husbandry

*Drosophila melanogaster* were raised on standard cornmeal/agar/molasses media at 25°C unless otherwise specified. To over-express constructs of interest, the UAS/GAL4 system was used (Brand and Perrimon, 1993), with the *hedgehog-GAL4* (*hh-GAL4*) driver. Prepupa of the appropriate genotype were selected and aged at 25°C as indicated.

For induction and recombination experiments *hsFLP* was used to excise an *FRT-Stop-FRT* cassette to allow ubiquitous expression of *Fz-eYFP (P [w+, Actin>STOP>fz-eYFP])* or *Fz-mKate2-sfGFP* (*P[w+, Actin>STOP>fz-mKate2-sfGFP]attP2 fz^P21^*). Heat shocks were performed for two hours at 37°C at 24h APF for microtubule experiments, or for one hour at 37°C at the specified time for other experiments. On the basis of when trichomes subsequently emerge, we observe that development is halted for the period of the heat shock. Pupa were left to age at 25°C before dissecting and were fixed at the same developmental time of 28h APF unless otherwise stated.

Mutant alleles are described in FlyBase; *stbm^6^*, *pk-sple^13^*, *fmi^E59^*, *fz^P21^* and *dgo^380^* are putative null alleles and unable to give rise to functional proteins, while *dsh^1^* gives rise to a mutant protein defective for planar polarity but which functions normally in Wingless signalling (Axelrod et al., 1998; Boutros et al., 1998; Warrington et al., 2017). Importantly, by the criterion of abolishing detectable asymmetric localisation of the other core proteins or their downstream effectors, the *stbm^6^*, *pk-sple^13^*, *fmi^E59^*, *fz^P21^* and *dsh^1^* alleles are completely lacking in planar polarity function (Wong and Adler, 1993; Usui et al., 1999; Axelrod, 2001; Strutt, 2001; Tree et al., 2002; Bastock et al., 2003; Strutt and Warrington, 2008; Lu et al., 2010; Warrington et al., 2017).

Full genotypes for each experiment are provided in Supplemental Table 1.

### Dissection and immunolabelling of pupal wings

Pupal wings were dissected at 28h APF at room temperature. Briefly, pupae were removed from their pupal case and fixed for 40 min in 4% paraformaldehyde in PBS, or 50 min in 10% paraformaldehyde in PBS for microtubule immunolabelling. Wings were then dissected and the outer cuticle removed, fixed for 10 min in 4% paraformaldehyde in PBS or in 10% paraformaldehyde in PBS for microtubule immunolabelling and were blocked for 1h in PBS containing 0.2% Triton X100 (PTX) and 10% normal goat serum. Primary and secondary antibodies were incubated overnight at 4°C in PTX with 10% normal goat serum, and all washes were in PTX. After immunolabelling, wings were post-fixed in 4% paraformaldehyde in PBS for 10 min. Wings were mounted in 12.5 μl of Vectashield or in 25 μl of PBS containing 10% glycerol and 2.5% DABCO, pH7.5 for microtubule experiments.

### Imaging

Pupal wings were oriented along the proximodistal axis using longitudinal vein 4 as the reference and were imaged on a Nikon A1R GaAsP confocal microscope using a 60x NA1.4 apochromatic lens. Imaging of Fz::mKate2-sfGFP was done on live pupae, where a small piece of cuticle was removed from above the pupal wing, and the exposed wing was mounted on strips of double-sided tape in a glass-bottomed dish. Wings were imaged with a pixel size of 80 nm, and the pinhole was set to 1.2 AU. 9 Z-slices separated by 150 nm were imaged, and then the 3 brightest slices around junctions were selected and averaged for each channel in ImageJ.

For microtubule imaging, fixed pupal wings were imaged with a pixel size of 40 nm, and the pinhole was set to 0.4 AU. 19 Z-slices separated by 70 nm were captured. Microtubule images were then deconvolved following the Landweber’s method with point scan confocal modality with 10 iterations. The 3 brightest slices around junctions were selected from the deconvolved image stack and averaged for each channel in ImageJ.

### Prehairs

For prehair visualisation, prepupa were aged for 22h at 27°C. Heat shocks were performed for one hour at 37°C. Pupa were left to age at 27°C before dissecting and were fixed at the same developmental time of 33h APF. Wings were processed for immunofluorescence as described for core protein detection, except to improve labelling of actin structures, wings were fixed for 50 min in supplemented fix with 1% Triton X-100; GFP protein was detected using 1:4000 rabbit anti-GFP (Abcam) and Actin was visualised using 1:1000 Alexa568-Phalloidin. Wings were oriented along the proximodistal axis using longitudinal vein 4 as the reference and were imaged on a Nikon A1R GaAsP confocal microscope using a 60x NA1.4 apochromatic lens. 9 Z-slices separated by 150 nm were imaged, and then the 3 brightest slices around junctions were selected and averaged for each channel in ImageJ.

### Western blotting

For pupal wing western blots, 28 h APF pupal wings were dissected directly into sample buffer (141 mM Tris base, 2% lithium dodecyl sulphate,10% glycerol, 0.51 mM EDTA, 100 mM dithiothreitol, pH 8.5) and the equivalent of the indicated number of wings per lane were run. A Bio-Rad ChemiDoc XRS+ was used for imaging, and band intensities from four biological replicates were quantified using ImageJ. Data were compared used unpaired t tests or ANOVA for multiple comparisons. Western blots were probed with mouse monoclonal anti-Fmi 74 (DSHB, Usui et al., 1999), affinity-purified rabbit anti-Fz (Bastock and Strutt, 2007), affinity-purified rat anti-Pk (Strutt et al., 2013), affinity-purified rabbit anti-GFP (Abcam ab6556) and Mouse monoclonal anti-Tubulin DM1A (Sigma-Aldrich T9026). We do not have an anti-Dgo antibody suitable for western blotting.

### Adult wings

Adult wings were dehydrated in isopropanol and mounted in GMM (50% methylsalicylate, 50% Canada Balsam), and incubated overnight on a 60°C hot plate to clear. Wings were photographed under brightfield illumination on a Leica DMR compound at 20x magnification, with longitudinal vein 4 as the horizontal reference.

### Quantification and statistical analysis

#### Fluorescence detection and quantitation

Membrane masks were generated using Tissue Analyzer (Etournay et al., 2016), and the mean intensity of fluorescence in membranes was measured using an automated MATLAB script (see **Puncta_defined_area** in MATLAB scripts in Supplemental Material and (Strutt et al., 2016)).

For Fig.6 cell boundary quantitations, fluorescence on medio-lateral oriented junctions or on horizontal oriented junctions was measured by manual drawing of ROIs in ImageJ. All boundaries were classified following their orientation compared to the Fz over-expression boundary generated with the UAS-GAL4 system using *hedgehog-GAL4*, with the *hedgehog* expression domain in the posterior part of the pupal wing. Horizontal junctions were defined as parallel to the Fz over-expression boundary (between 0 and 45 degrees) and medio-lateral junctions as junctions linking two horizontal boundaries (between 45 and 90 degrees).

For live imaging, background due to autofluorescence was subtracted, and mean fluorescence intensity was averaged across wings.

The fraction of stable Fz was determined by ratio of mKate2/sfGFP fluorescence intensity. When all genotypes were compared, an ANOVA with Tukey’s multiple comparisons test was used, or ANOVAs with Dunnett’s or Kruskal-Wallis multiple comparisons tests were used to compare wild-type to mutant backgrounds.

Total Fz intensities were compared with an ANOVA with Holm-Sidak’s multiple comparisons test to weigh all conditions or with unpaired t test and Mann-Whitney test to compare fluorescence intensities from two conditions.

#### Planar polarity

For polarity magnitude and cell morphological properties analysis QuantifyPolarity software was used (Tan et al., 2021). Wing images were oriented along the proximo distal wing axis based on longitudinal wing vein orientation corresponding to the x axis, and border masks were generated using Tissue Analyzer (Aigouy et al., 2010) to delimit cells. A cell-by-cell analysis was performed, with polarity magnitude and orientation calculated using the Principle Component Analysis (PCA) option in QuantifyPolarity. Results are given for individual cell and average per wing. In this algorithm, a cell is first transformed onto an ellipse shape to obtain cell orientation and eccentricity. Then, for polarity measurement, intensities of individual pixels are normalised and the polarity angle is calculated as the angle that produces the largest variance of normalised intensities. Finally, the normalised intensities are converted into pseudo-XY-coordinates and the eigenvalues of the covariation matrix of these transformed coordinates are used to compute the polarity magnitude. Planar polarity quantification is independent of cell geometry. The full description of the algorithm is available in (Tan et al., 2021).

For Figure 5, custom MATLAB scripts were used to combine polarity magnitude and orientation (see **Combined results for polarity** in MATLAB scripts in Supplemental Material), and these were visualised on polar plots (see **Polar plot for polarity** for details in MATLAB scripts in Supplemental Material). Statistical tests are described in legend text for each figure. For polarity magnitude comparisons, when all genotypes were compared between them, an ANOVA with Tukey’s multiple comparisons test was used, or an ANOVA with Dunnett’s multiple comparisons tests was used to compare wild-type to mutant backgrounds, or unpaired Mann-Whitney tests were used to compare with or without repolarisation induction. To compare two conditions for total Fz polarity (orientation and magnitude), Hotelling’s T-square tests were used.

#### Cell and microtubule orientation

Analysis of cell orientation and apical microtubule orientation used custom automated MATLAB image analysis scripts, based on an initial protocol developed by (Gomez et al., 2016; Płochocka et al., 2021) and improved by (Ramirez-Moreno et al., 2023) (see **Microtubule_orientation** in MATLAB scripts in Supplemental Material). When all genotypes were compared between them, an ANOVA with Tukey’s multiple comparisons test was used, or unpaired Mann-Whitney tests were used to compare with or without repolarisation induction.

### Description of MATLAB scripts

#### Puncta_defined_area

This is a MATLAB script that measures membrane and puncta intensities on segmented images (Strutt et al., 2016) generated using Tissue Analyzer (Etournay et al., 2016), and was used here to measure total membrane intensity (Mmean).

#### Microtubule_orientation

The average projections of images with tubulin signals were adjusted so that 0.5% of the pixels with the lowest intensities were set to black and the 0.5% of the pixels with the highest intensities were set to white to normalise the variability in signal between images and increase the contrast. Masks generated using Tissue Analyzer (Etournay et al., 2016) were used to isolate tubulin signals in individual cells. Then, the magnitude of the tubulin signal according to its direction (gradient of the signal) in each cell was calculated by convolving the tubulin signal using two 5 × 5 Sobel operators (Gomez et al., 2016). The resulting distributions of tubulin signals were aligned to their maxima at 0 and averaged for cells with specified eccentricities (0.750 ± 0.025 and 0.800 ± 0.025) to produce average profiles of microtubule angle distributions in cells (Płochocka et al., 2021). At the same time, the unaltered distributions were fitted with the Von Mises distribution and the estimated mean and standard deviation of the fitted curve in each cell. The mean was used as the main direction of the apical microtubule network in this cell and the standard deviation as the measure of the microtubule alignment with each other (Microtubule Standard Deviation, MTSD). Finally, the cell directions were calculated by fitting the pixel coordinates of each cell isolated using masks to an ellipse and obtaining the direction of the long axis of the best-fit ellipse. MTSD was plotted against cell eccentricity for cells with a MTSD <90 excluding cells with unfittable distributions of tubulin signal. This MATLAB code is available at https://github.com/nbul/Cytoskeleton/tree/master/PCP-MT. Angle visualisation and quantification data about directions of cell elongation (the direction of the long axis of the best-fit ellipse), overall directions of microtubule networks and Stbm polarity angle (transferred from QuantifyPolarity) were plotted on image masks produced using Tissue Analyzer (Etournay et al., 2016). Here, the length of plotted lines does not reflect polarity magnitudes or length of elongation but is fixed at half of the average long cell axis. Average cell orientation within each image was used to normalise the Stbm polarity angle, individual cell orientations and microtubule angles for each cell in polar histograms with 9 bins in 0-180° format. Data were mirrored to ease visualisation. Normalised angle data was also used to separate angles into quadrants: between −45°, 45° and between 45°, 135°. The plotting and calculations were performed within the script for the microtubule organisation analysis (https://github.com/nbul/Cytoskeleton/tree/master/PCP-MT).

#### Combined results for polarity

This is a MATLAB script suitable for combining polarity magnitude and angle results generated by QuantifyPolarity software (see Supplemental Material).

#### Polar plot for polarity

This is a MATLAB script suitable for generating polar plots showing polarity magnitude and angle, with results from ‘Combined results for polarity (magnitude + angle)’ MatLab script (see Supplemental Material).

---

**Supplemental Figure 1.**
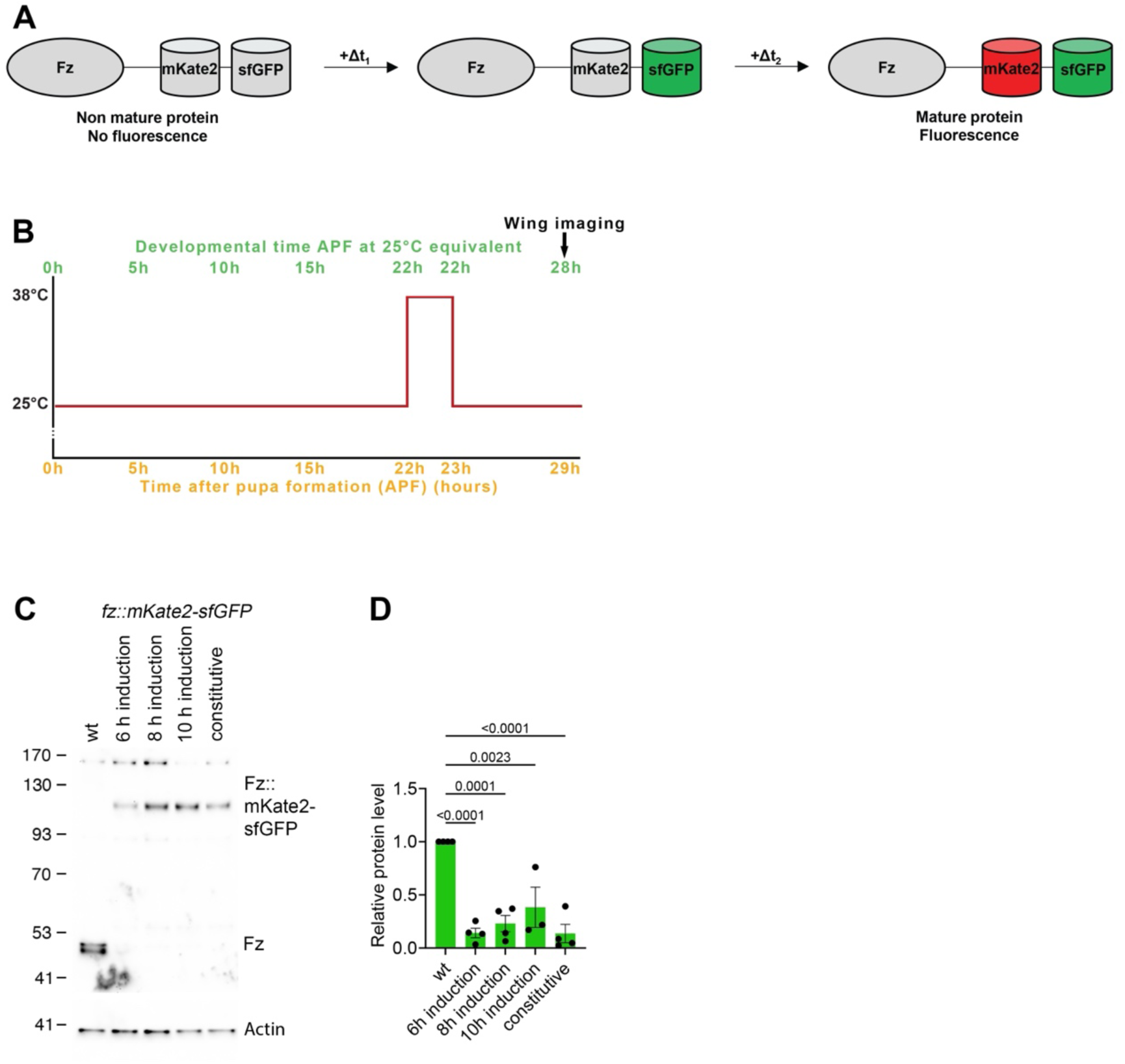
Tools for Fz induction and measurement of stable fractions. **(A)** Schematic illustration of evolution over time of fluorescent timer (mKate2 and sfGFP) fused with Fz. **(B)** Scheme of induced Fz tagged with fluorescent timer expression by heat shock induction, with Fz observation at 28h APF. Here is represented the condition with 6h between tagged Fz expression induction and its observation. With 8h between tagged Fz expression induction and its observation at 28h APF, heat shock is performed at 20h APF and for 10h time lapse, at 18h APF. **(C)** Western blot probed for Fz protein with Actin for loading control. Extracts are from pupal wings at 28h APF with wild-type Fz or constitutive or *de novo* induced (6h, 8h and 10h induction) Fz::mKate2-sfGFP expression, and 2 wings were loaded for each sample. **(D)** Quantification from four biological replicates of Fz and Fz::mKate2-sfGFP levels in conditions described in **(C),** normalised with wild-type Fz level. Error bars are SEM. ANOVA with Tukey’s multiple comparisons test was used to compare all genotypes, only significant p values are indicated.

**Supplemental Figure 2.**
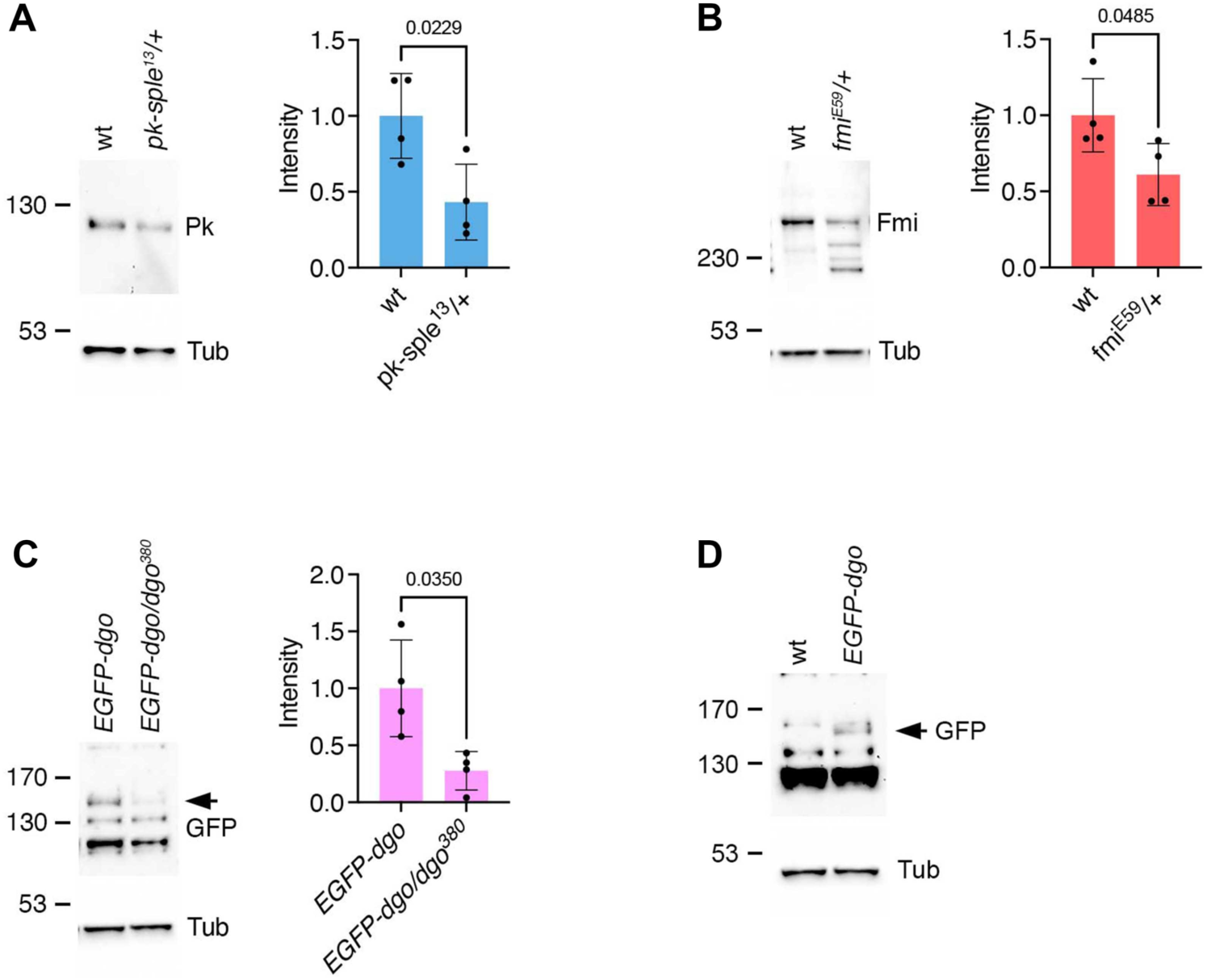
Decreasing *prickle, flamingo* and *diego* gene dosage results in lower protein levels. **(A-C)** Western blots and quantifications from pupal wings, comparing levels of Pk from *pk-sple^13^/+* heterozygous flies **(A)**, or levels of Fmi from *fmi^E59^/+* heterogygous flies to wild-type **(B)**, or comparing levels of EGFP reflecting Dgo levels from *dgo^380^/EGFP-dgo, dgo^380^* heterozygous flies to *EGFP-dgo, dgo^380^/EGFP-dgo, dgo^380^* flies **(C)**. EGFP-Dgo is expressed from a rescue transgene in a *dgo^380^* null mutant background. We used this as a proxy for endogenous Dgo levels, as we do not have an antibody against Dgo that works on western blots. Quantifications are from four biological replicates, unpaired t-test. 2 pupal wings were loaded for Pk and Fmi and 8 for the EGFP-Dgo Western blot. **(D**) Western blot showing EGFP band (arrow) in EGFP-Dgo compared to wild-type pupal wings to demonstrate that the correct EGFP band was identified, as the anti-EGFP antibody used also gives several cross-reacting bands. 8 pupal wings were loaded for this blot.

**Supplemental Figure 3.**
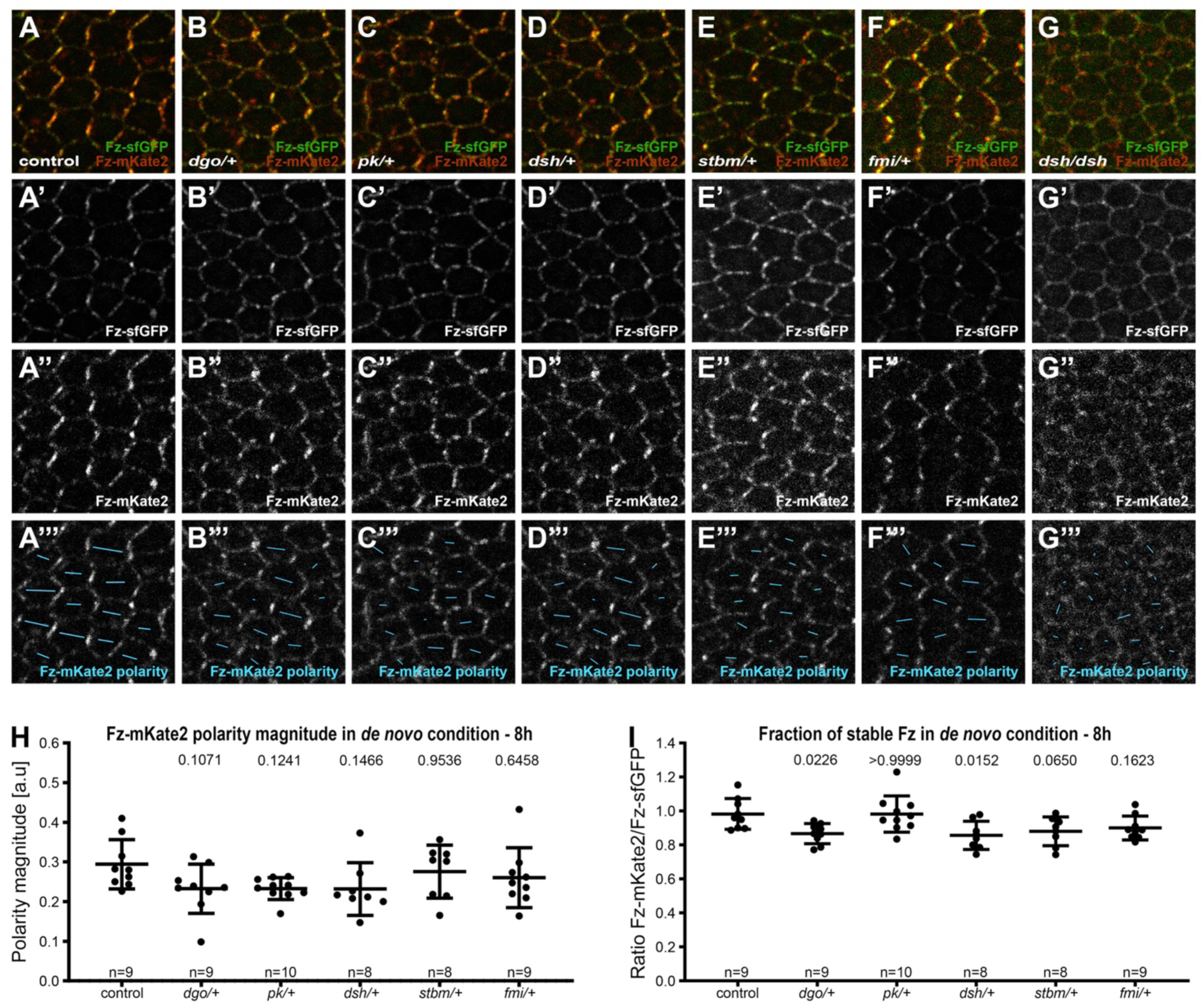
Core protein polarity patterns under control conditions and in *de novo* condition with 8h induction to establish planar polarity. **(A-G)** Live confocal images of 28h APF pupal wing epithelia expressing Fz::mKate2-sfGFP taken below longitudinal vein 4, see **Fig. 1C’**. See **Table S1** for the full genotypes. Native fluorescence for sfGFP (green, **A’-G’)** and mKate2 (red, **A’’-G’’)**, with constitutive expression of Fz::mKate2-sfGFP in an otherwise wild-type background **(A)**, or in heterozygous mutant backgrounds for *dgo* **(B)**, *pk* **(C)**, *dsh^1^* **(D)**, *stbm* **(E)**, *fmi* **(F)**, and hemizygous mutant background for *dsh^1^* **(G)**. See **Fig.S1B** for details of Fz tagged with fluorescent timer. **(A’’’-G’’’)** Cell-by-cell polarity pattern of mKate2 fluorescence in pupal wings expressing Fz::mKate2-sfGFP at 28h APF. The length and orientation of cyan bars denote the polarity magnitude and angle respectively for a given cell. **(H, I)** Quantified polarity magnitude based on mKate2 fluorescence **(H)** or fraction of stable Fz as determined by ratio of mKate2/sfGFP **(I)**, in live pupal wings at 28h APF, in wild-type or core protein heterozygous mutant backgrounds in *de novo* condition with 8h induction of Fz:mKate2-sfGFP to establish core protein planar polarity. Error bars are SD; n, number of wings. ANOVA with Dunnett’s multiple comparisons tests were used to compare wild-type to mutant backgrounds, p values as indicated.

**Supplemental Figure 4.**
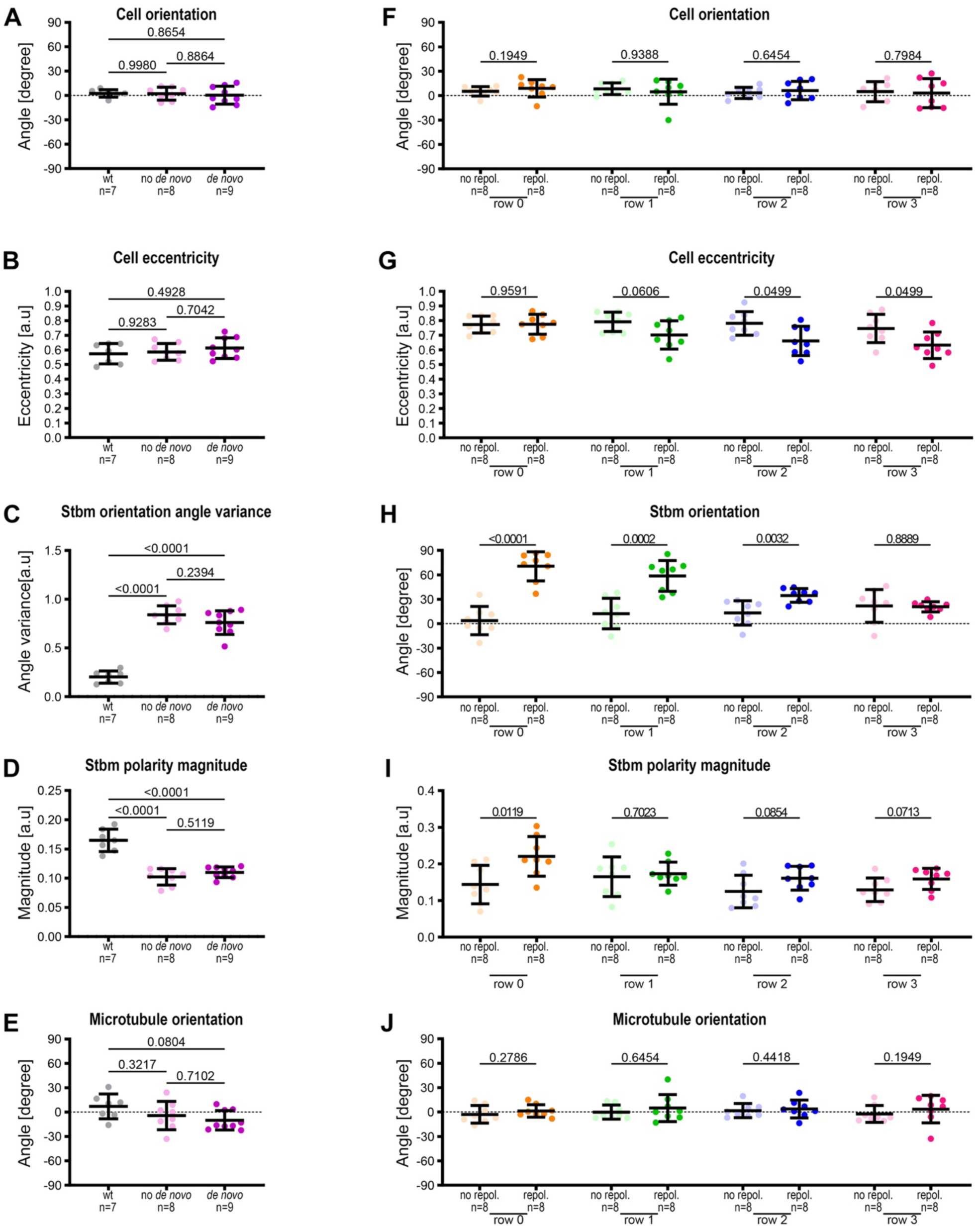
Quantification of cell shape, Stbm polarity and microtubule orientation in wild-type and *de novo* and repolarisation conditions at 28h APF. **(A-E)** Quantifications from conditions in **Fig.4A-C** of cell orientation relative to horizontal axis **(A)**, cell eccentricity **(B)**, Stbm orientation angle variance **(C)**, Stbm polarity magnitude **(D)** and microtubule orientation **(E)** relative to the average cell orientation, in wild-type wings, in *de novo* condition without induction (no *de novo*) or with (*de novo*) Fz-eYFP polarisation induction for 6h, in fixed pupal wings at 28h APF, in the region distal to the posterior cross vein, below longitudinal vein 4. Error bars are SD; n, number of wings. ANOVA with Tukey’s multiple comparisons tests were used to compare all genotypes, p values as indicated. See **Table S1** for the full genotypes. **(F-J)** Quantifications from conditions in **Fig. 4N-O** of cell orientation relative to horizontal axis **(F)**, cell eccentricity **(G)**, Stbm orientation angle **(H)**, Stbm polarity magnitude **(I)** and microtubule orientation **(J)** relative to the average cell orientation, without (no repol.) or with Fz (repol.) repolarisation induction for 6h, in fixed pupal wings at 28h APF. Cells are grouped in rows relative to their location relative to Fz overexpression in Fz repolarisation condition or relative to longitudinal vein 4 without Fz repolarisation with row 0 in contact with Fz overexpression boundary and row 3 furthest away. Error bars are SD; n, number of wings. Mann-Whitney tests **(F**, **G** and **J)** and Unpaired Mann-Whitney tests **(H**, **I)** were used to compare in each condition with or without Fz repolarisation polarisation induction, p values as indicated. See **Table S1** for the full genotypes.

**Supplemental Figure 5.**
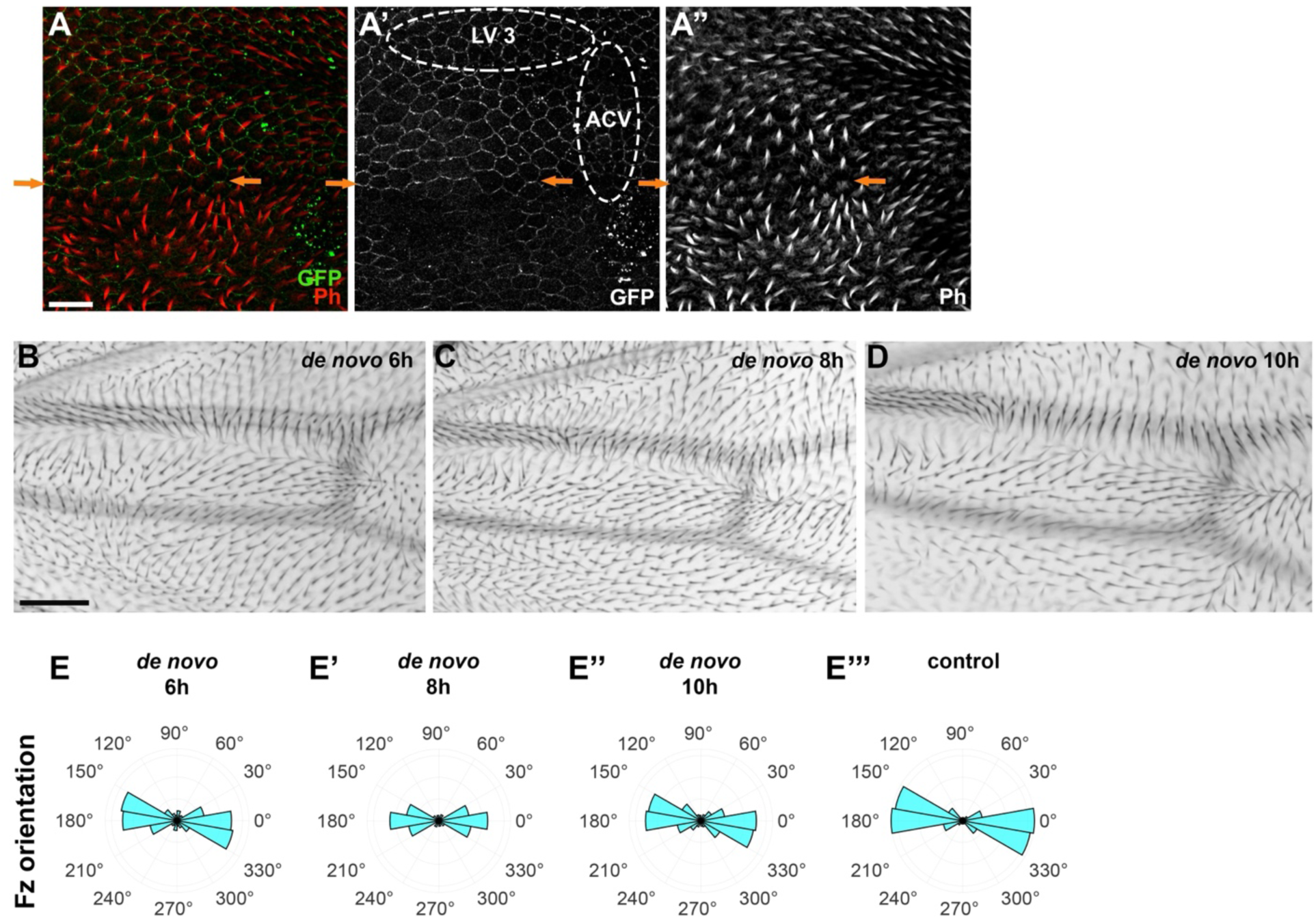
Trichome and core protein polarity in normal and *de novo* conditions. **(A)** Confocal microscope image proximal to the anterior cross vein (ACV) between the longitudinal vein 3 (LV 3) and the longitudinal vein 4 (LV 4) in a fixed wing at 33h APF. Wings stained for GFP revealing Fz localisation (green) **(A, A’)**, and for actin showing trichomes (red) **(A, A’’)**. In cells with Fz repolarisation on horizontal junctions, trichomes emerge repolarised along these same junctions with an antero-posterior orientation, observable in rows 0, 1, 2 and 3. Away from the Fz repolarisation area, trichomes emerges on distal cell edges with a proximo-distal orientation. Orange arrows indicate boundary 1, in row 0 which is the first repolarised row of cells. Scale bar 10 µm. For wing area localisation see **Fig.4M** and for cell and boundary localisations, see **Fig.6A**. See **Table S1** for the full genotype. **(B-D)** Microscope images of adult wings surrounding the experimentally analysed region in pupal wings proximal to the anterior cross vein (ACV), for *de novo* induction conditions with 6h **(B)**, 8h **(C)** and 10h **(D)**. Scale bar 50 µm. Note similar patterns seen under all induction conditions, with a swirling non-proximo-distal trichome orientations surrounding the proximo-distally oriented region just proximal to the ACV between LV 3 and LV 4. **(E-E’’’)** Polar histograms depicting binned Fz-mKate2 polarity orientation, based on mKate2 fluorescence in live pupal wings at 28h APF, with *de novo* condition with induction for 6h **(E)**, 8h **(E’)** 10h **(E’’)**, and in the control condition **(E’’’)**.

**Supplemental Figure 6.**
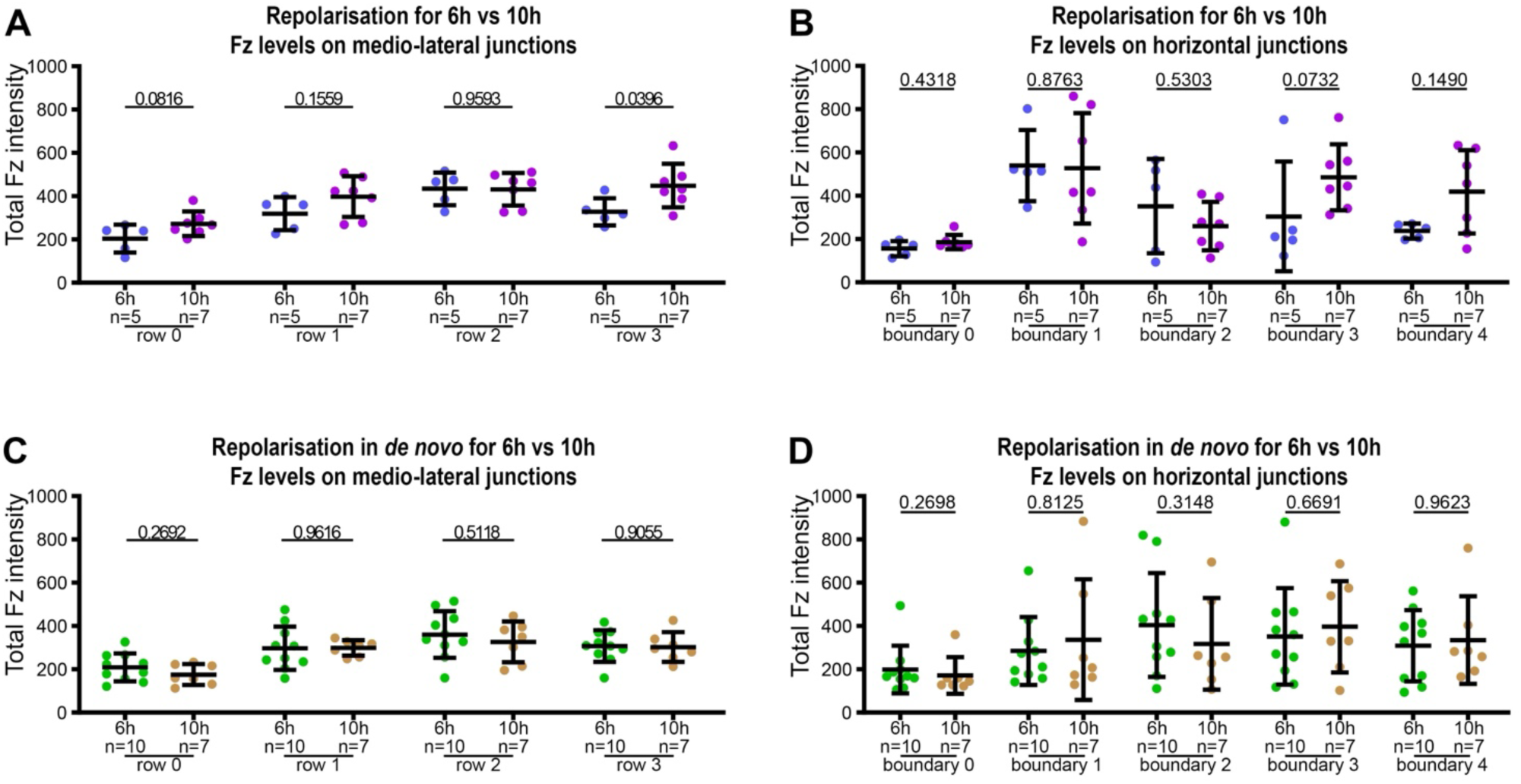
Distribution of Fz along medio-lateral and horizontal junctions in repolarisation and repolarisation under 6h and 10h *de novo* induction conditions. Quantitation of Fz intensity in fixed pupal wings at 28h APF, binned by row of cells relative to their location relative to Fz overexpression in Fz repolarisation and Fz repolarisation under *de novo* conditions, with row 0 in contact with Fz overexpression boundary and row 3 furthest away. See **Table S1** for the full genotypes. Error bars are SD; n, number of wings. **(A-B)** Comparison of Fz distribution in repolarisation condition with 6h or with 10h to repolarise, along medio-lateral junctions **(A)** and along horizontal junctions **(B)**. Unpaired t test **(A)** and Mann-Whitney test **(B)** were used to compare fluorescence intensity for 6h vs 10h conditions, p values as indicated. **(C-D)** Comparison of Fz distribution in repolarisation in *de novo* condition with 6h or 10h induction, along medio-lateral junctions **(C)** and along horizontal junctions **(D)**. Unpaired t test **(C)** and Mann-Whitney test **(D)** were used to compare fluorescence intensity at 6h vs 10h, p values as indicated.

**Table S1.**
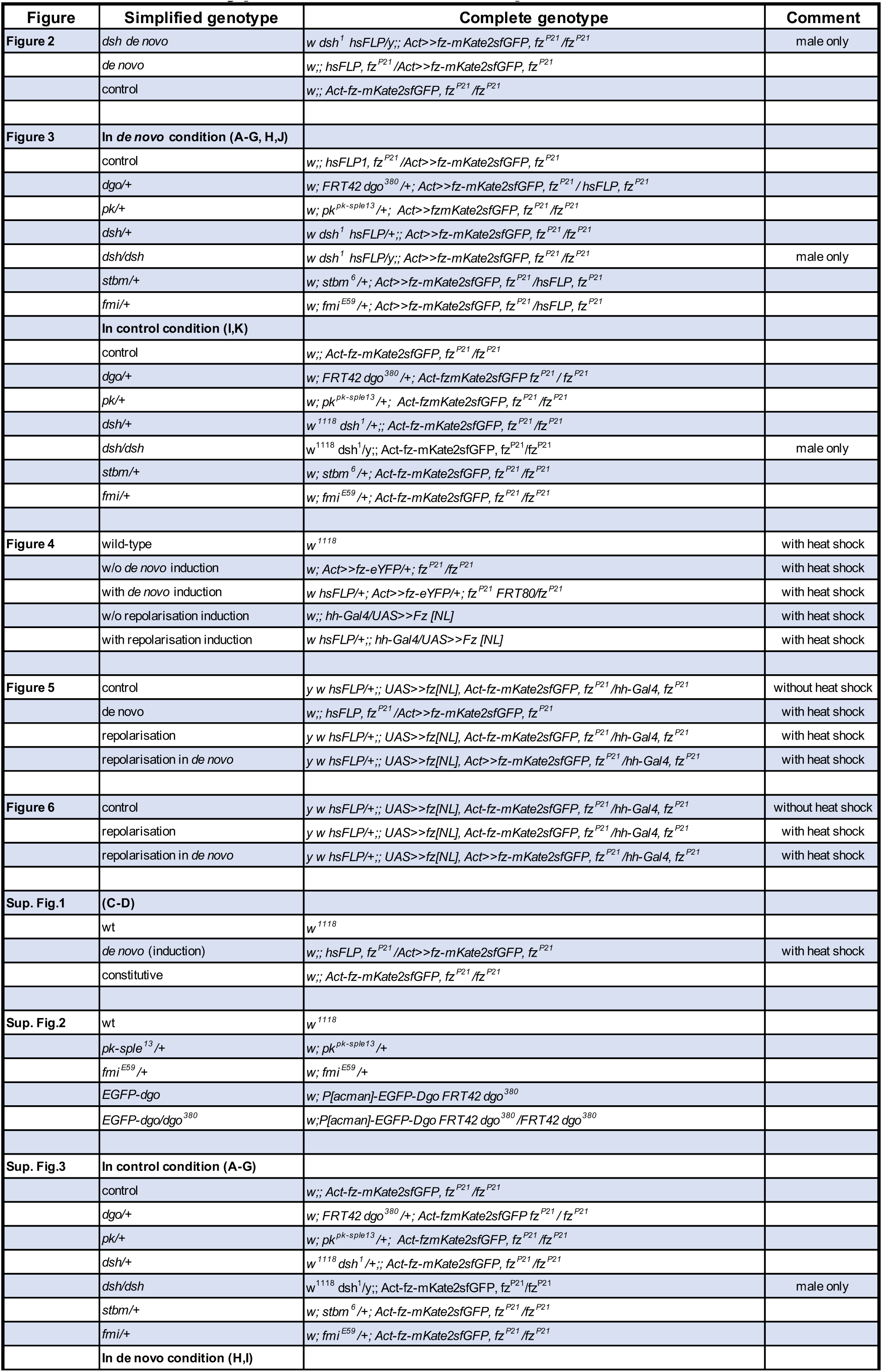
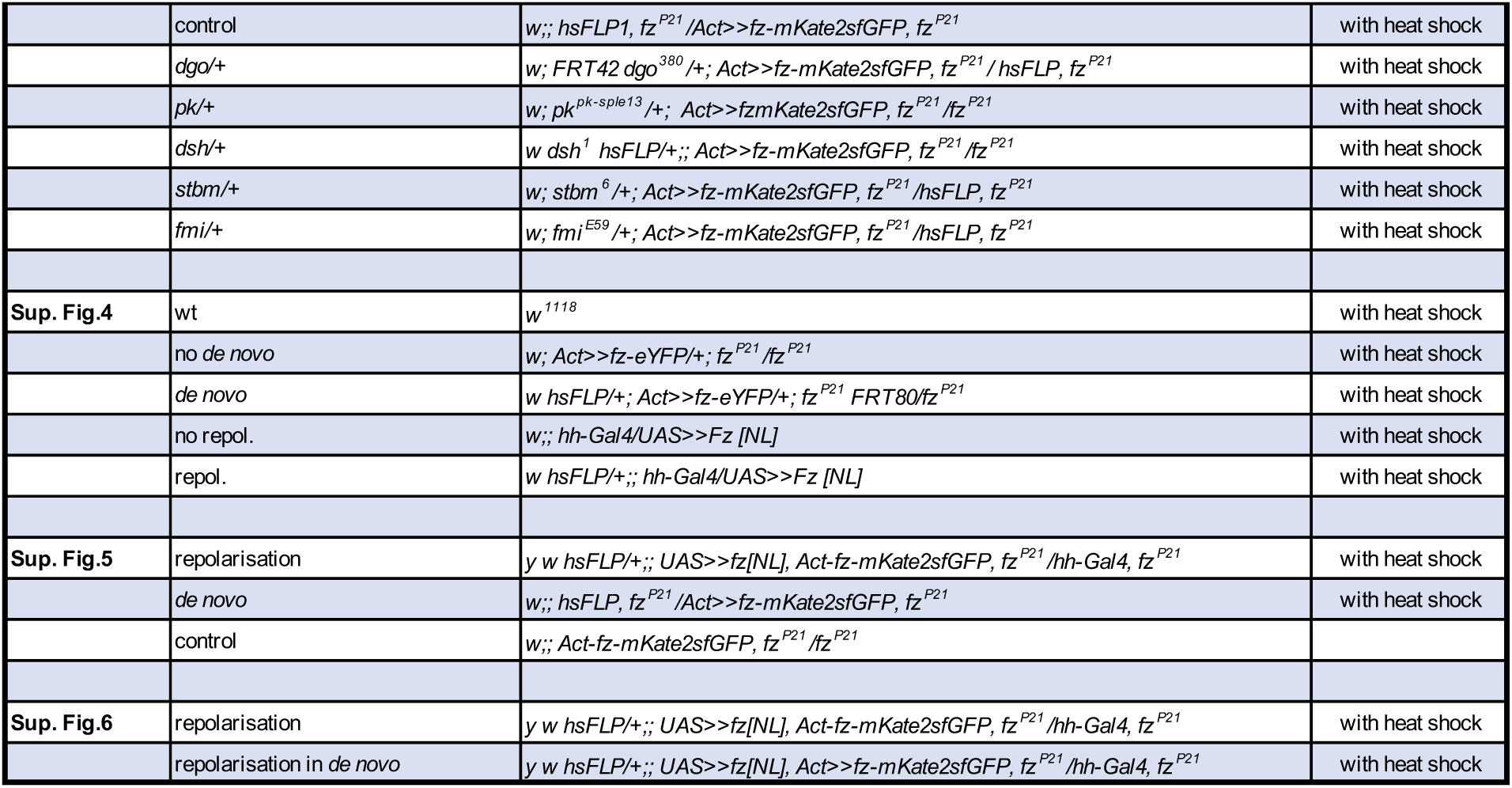
Genotypes used in each experiment.

**Table S2A.** Normality test for repolarisation from a boundary data (see Fig. 5I-L)

| Shapiro-Wilk normality test p-value | Control condition | *De novo* condition | Repolarisation condition | Repolarisation in *de novo* condition |
| --- | --- | --- | --- | --- |
| row 0 | 0.6254 | 0.2762 | 0.3410 | 0.0208 |
| row 1 | 0.3963 | 0.6336 | 0.0005 | 0.0657 |
| row 2 | 0.0673 | 0.2762 | 0.0015 | 0.5750 |
| row 3 | 0.0737 | 0.2762 | 0.0075 | 0.3361 |

Normality polarity angle combined with polarity magnitude
Before performing Hotelling's T2 test, confirm that data supports normality (using multivariate Shapiro-Wilks test) - p-value > 0,05 supports normality
Hotelling's T-square test is robust to violations of assumptions of multivariate normality
from https://online.stat.psu.edu/stat505/lesson/7/7.2/7.2.6

**Table S2B.** Hotelling’s T-square test for repolarisation from a boundary data (see Fig. 5I-L)

| Hotelling's T-square test p-value | row 0 | row 1 | row 2 | row 3 |
| --- | --- | --- | --- | --- |
| *De novo* vs control condition | 0.5751 | 0.1626 | 0.8424 | 0.3533 |
| Control vs repolarisation condition | 0.0014 | 0.2945 | 0.2258 | 0.0095 |
| Repolarisation vs repolarisation in *de novo* condition | 0.5096 | 0.4845 | 0.1650 | 0.7833 |
| Repolarisation in *de novo* vs *de novo* condition | 0.0011 | 0.0018 | 0.2634 | 0.0197 |

| Hotelling's T-square test p-value | Control condition | *De novo* condition | Repolarisation condition | Repolarisation in *de novo* condition |
| --- | --- | --- | --- | --- |
| row 0 vs row 1 | 0.5567 | 0.6200 | 0.1235 | 0.0731 |
| row 1 vs row 2 | 0.8061 | 0.2296 | 0.2164 | 0.1779 |
| row 2 vs row 3 | 0.6773 | 0.0037 | 0.0178 | 0.4499 |
| row 0 vs row 2 | 0.6014 | 0.6112 | 0.0041 | 0.0106 |
| row 1 vs row 3 | 0.2341 | 0.2341 | 0.1017 | 0.0167 |
| row 0 vs row 3 | 0.0804 | 0.1095 | 0.0333 | 0.0024 |

## Supplemental Materials – Analysis script 1

### Combined results for polarity (magnitude + angle)

This is a MATLAB script suitable for combining polarity magnitude and angle results generated by QuantifyPolarity software.

This script is designed to analyse files for 4 conditions (row1, row2, row3 and row4). In order to use it you have to strictly follow the following architecture:

- Create a folder, for row1 analysis, named ‘Analysed_sfGFP_row1’ containing analysed data from QuantifyPolarity software which own ‘Result’ subfolder. Do the same thing for row2 up to row4.
- Run the script and choose the appropriate folder location when prompted.

The script redistributes polarity angles in a range between −45° and 135° from −90° and +90° for each analysed cells per wing for all conditions. For each wing of the same condition (for example ‘row1’), those reoriented angles are averaged as polarity magnitude and added in a new saved .csv file ‘Analysed_sfGFP_row1’ in column ‘Average_Reoriented_Angle_Polarity_deg’. For each condition, the mean of polarity magnitude and angle is calculated and added in a saved new .csv file ‘Average_Reoriented_Angle_Polarity_row1’.

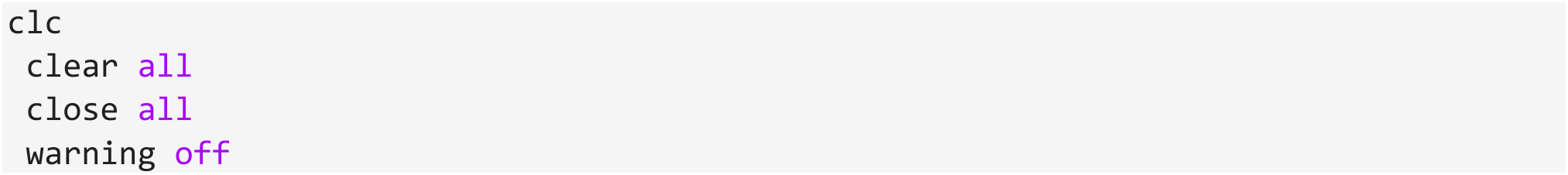

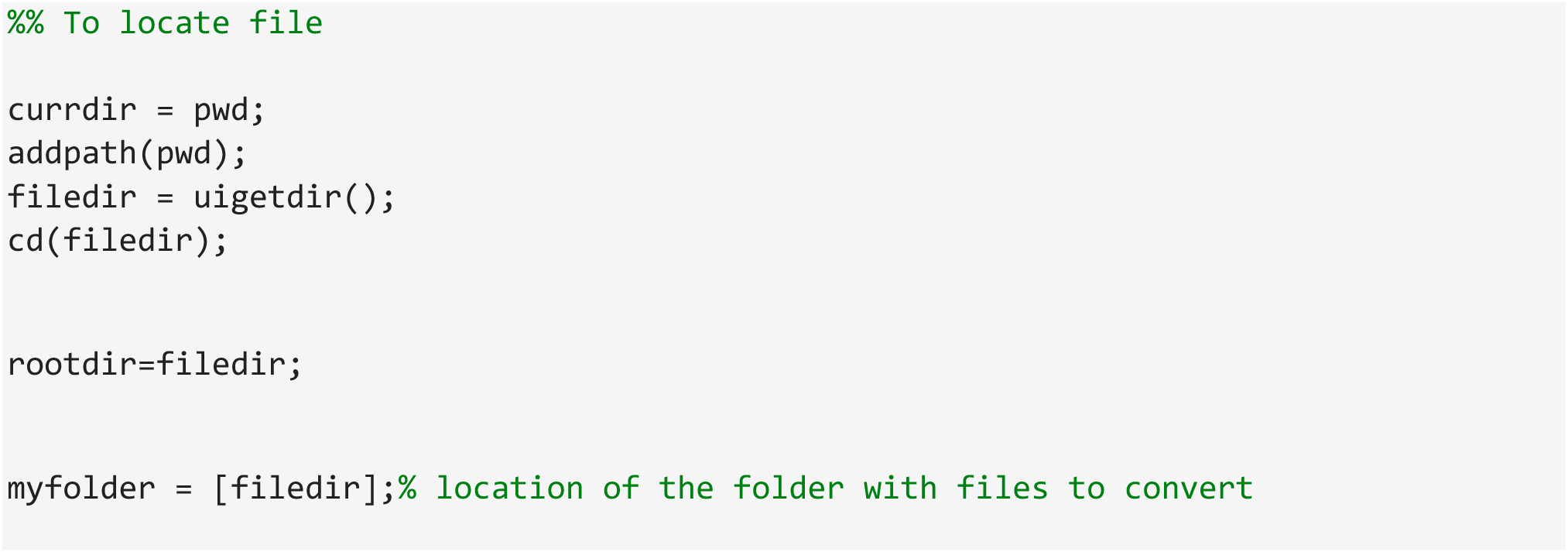

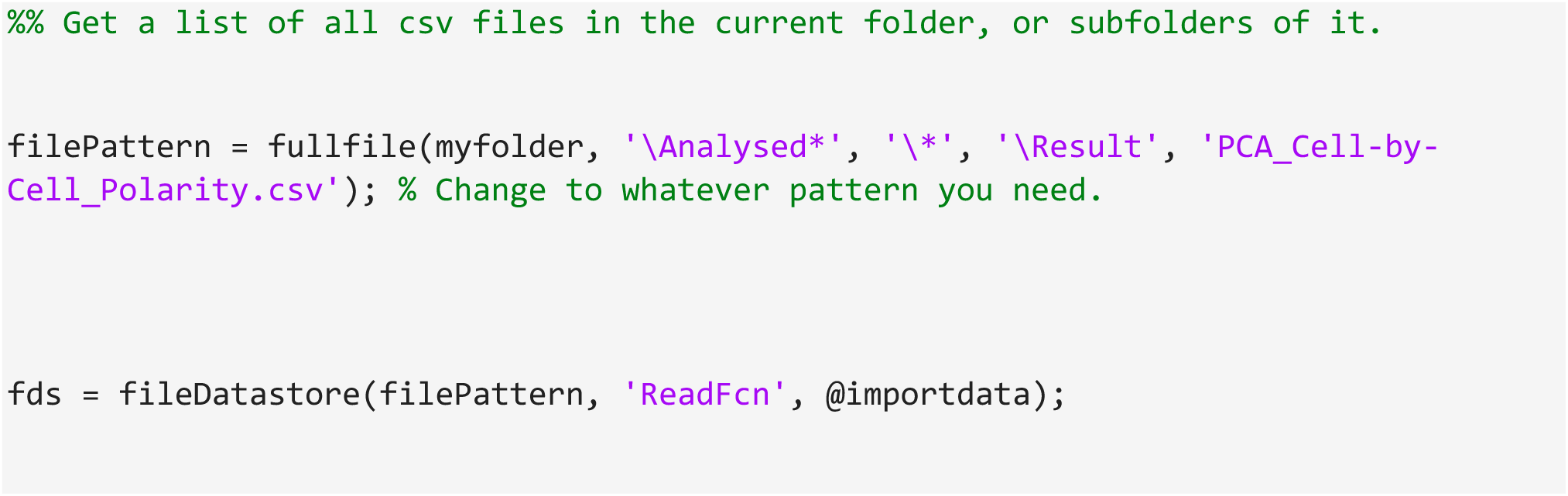
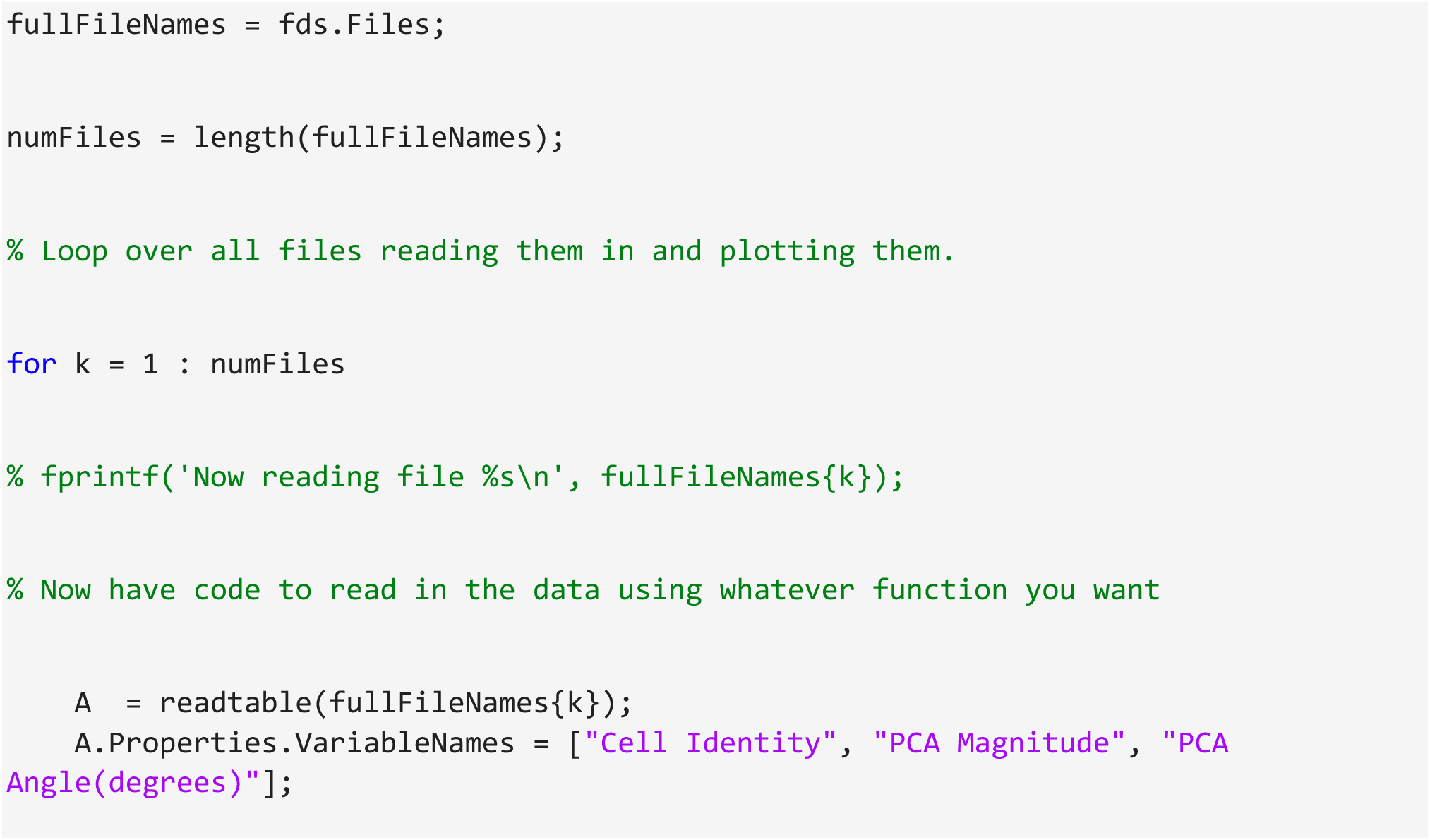

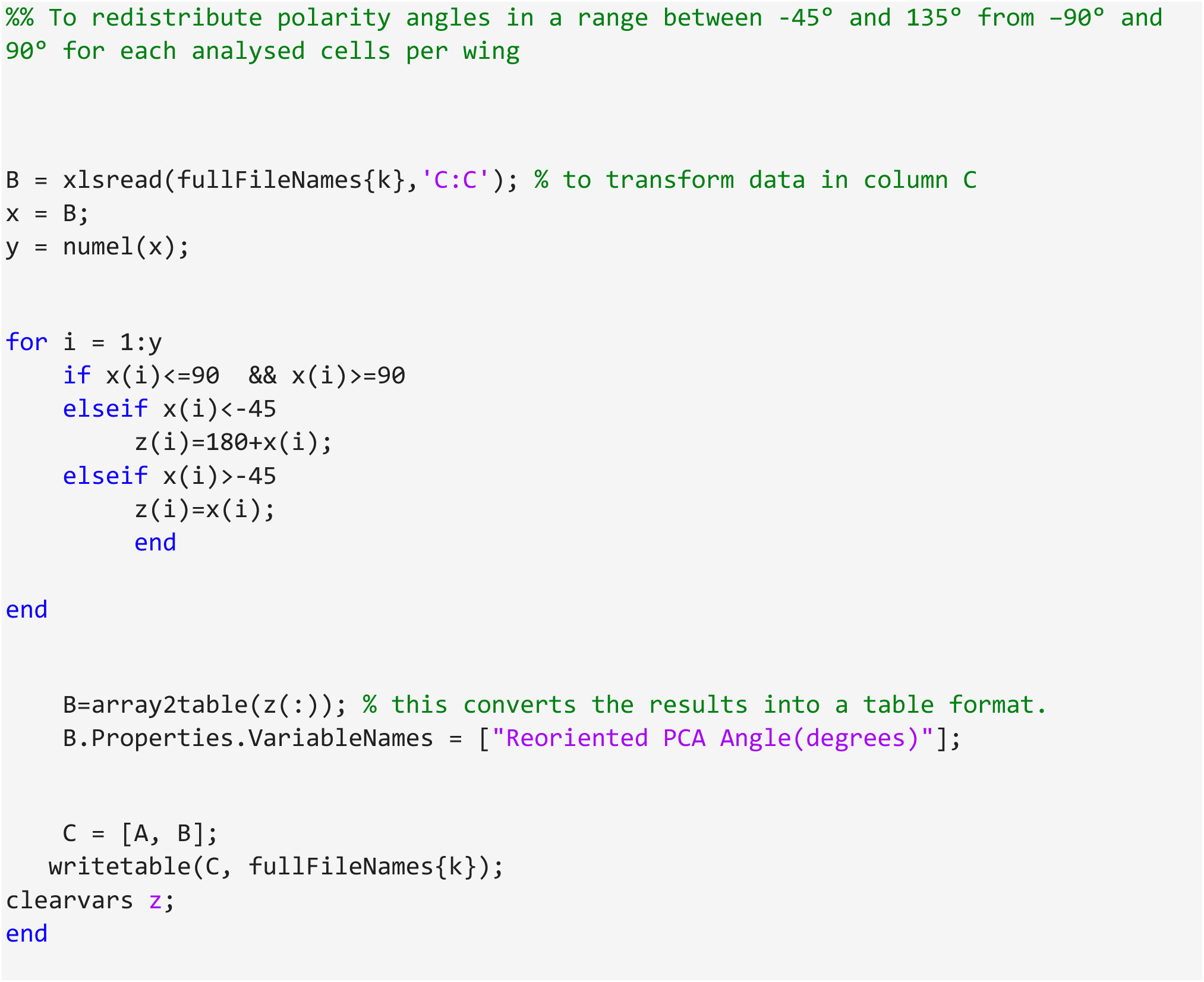
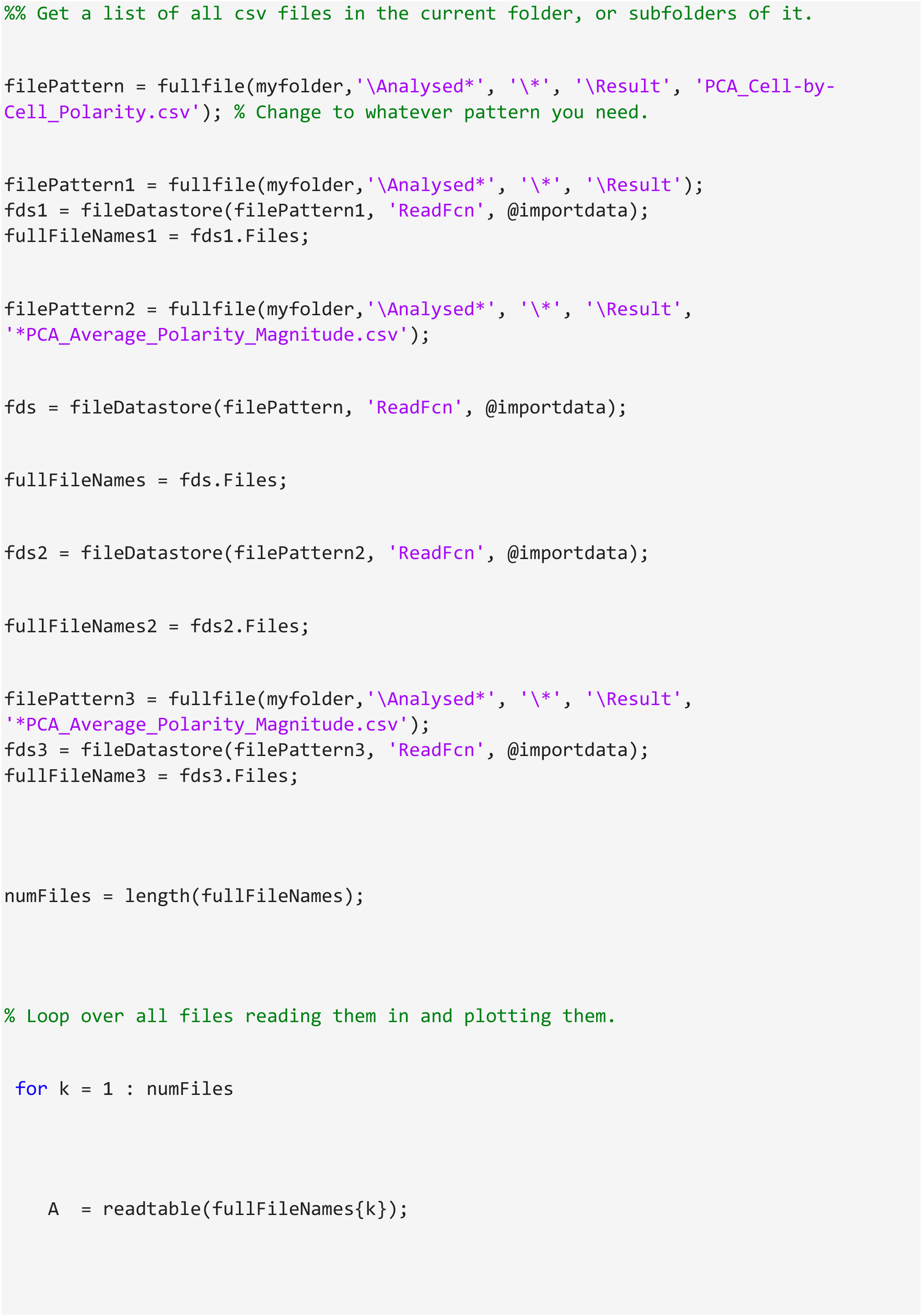
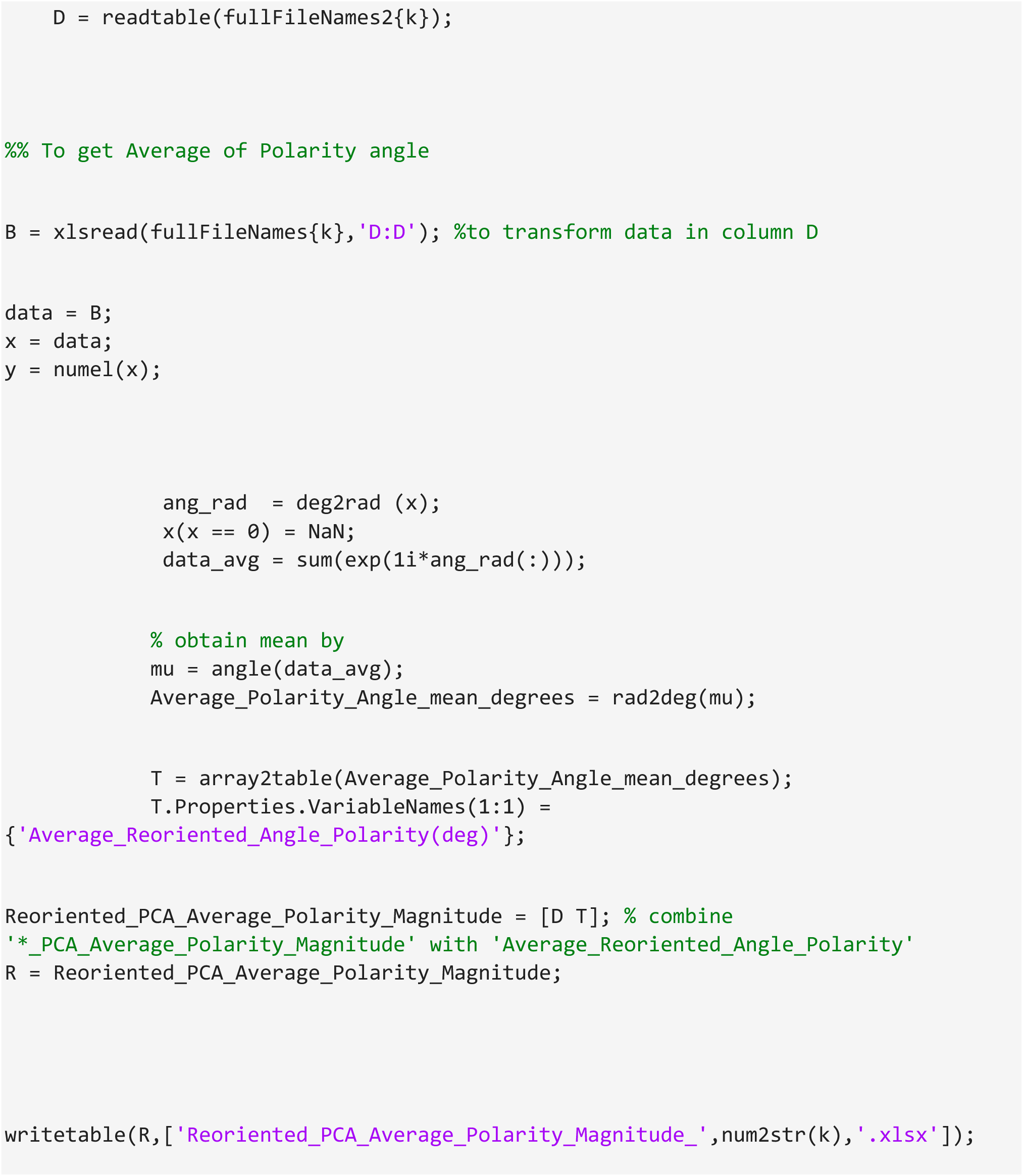

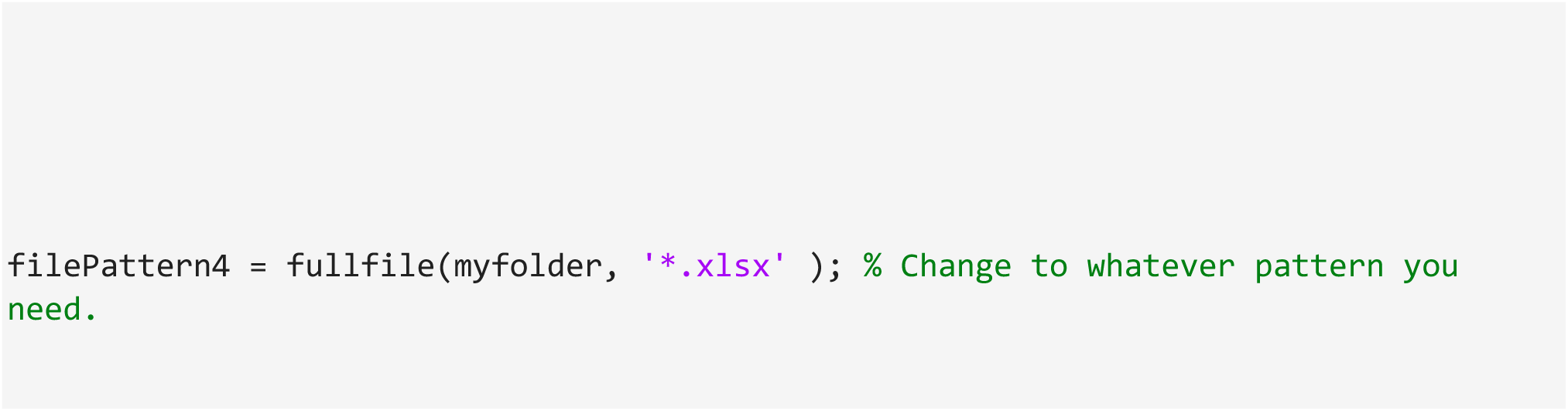
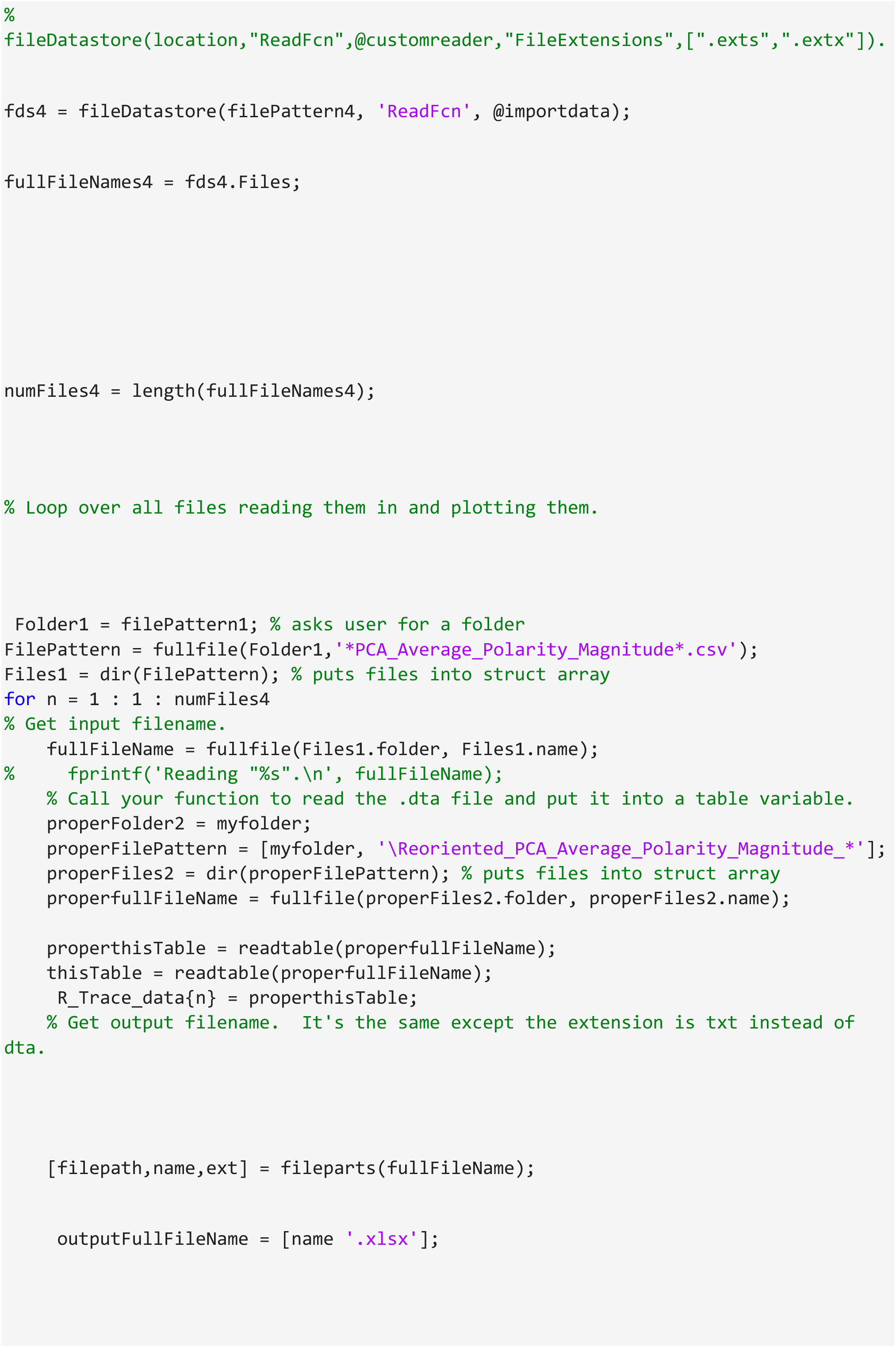
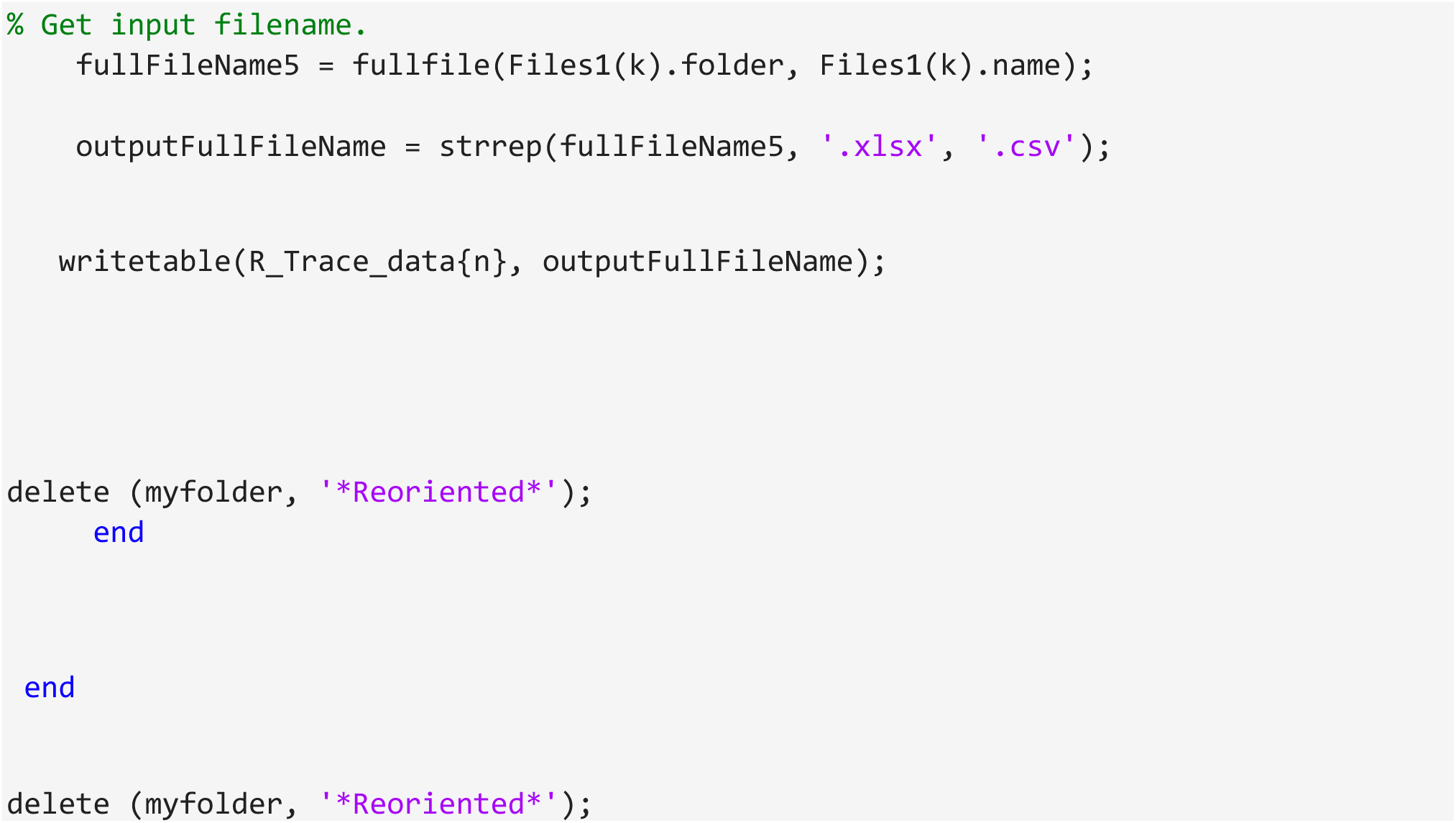

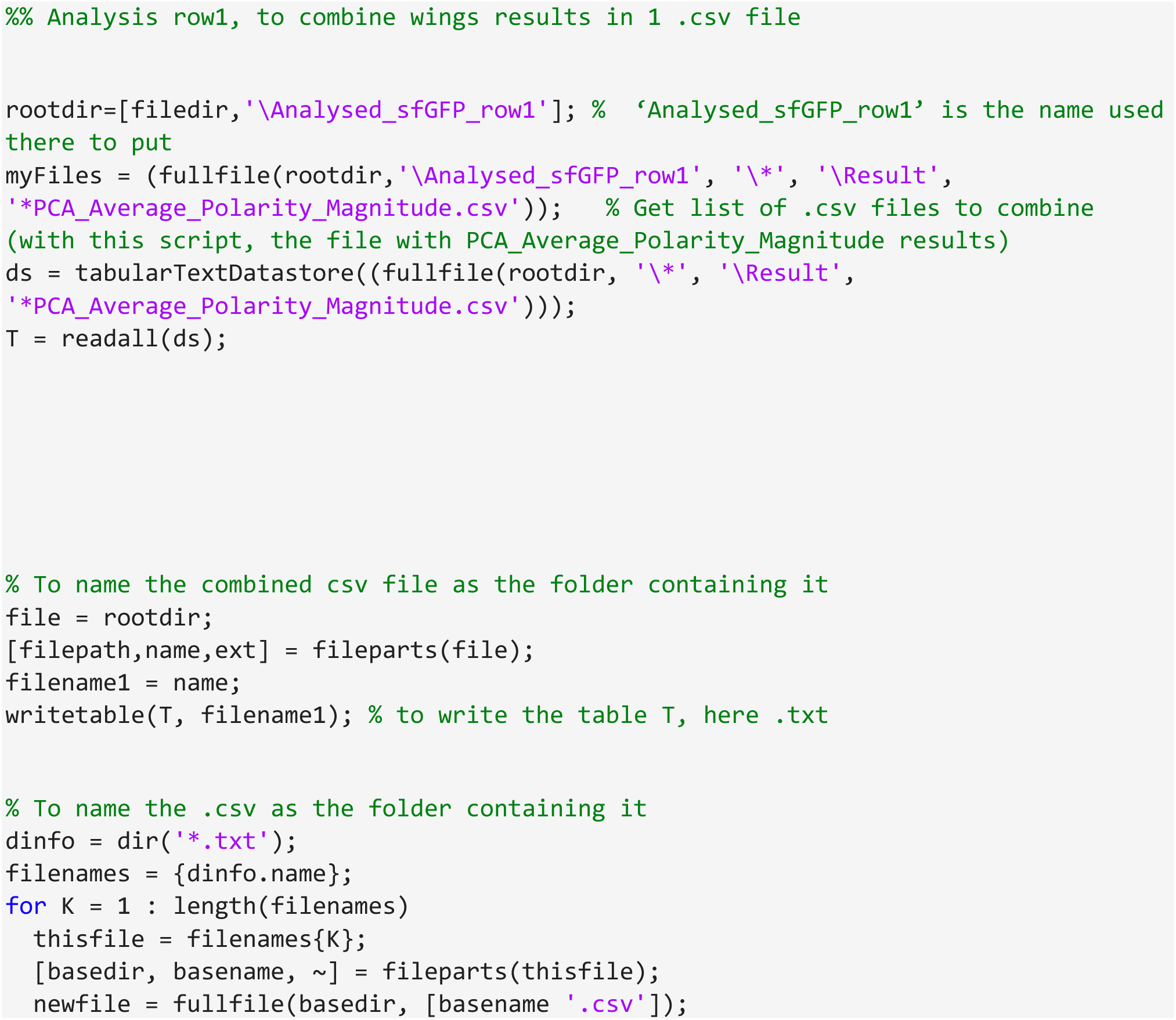
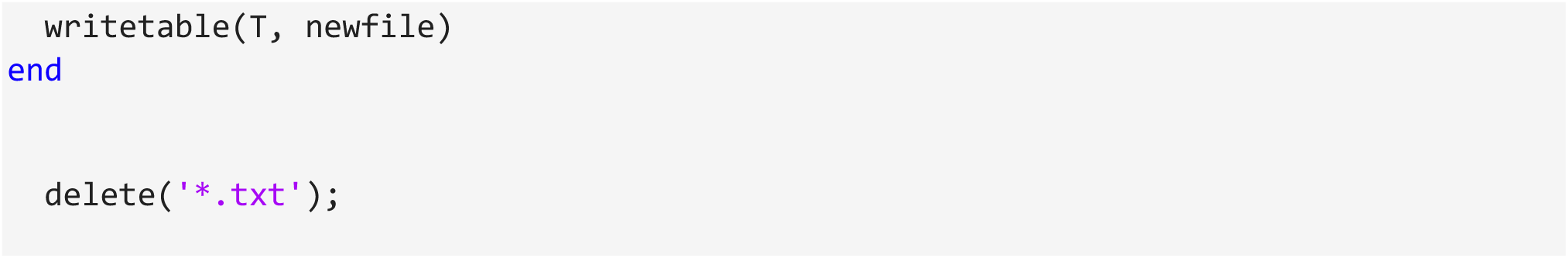

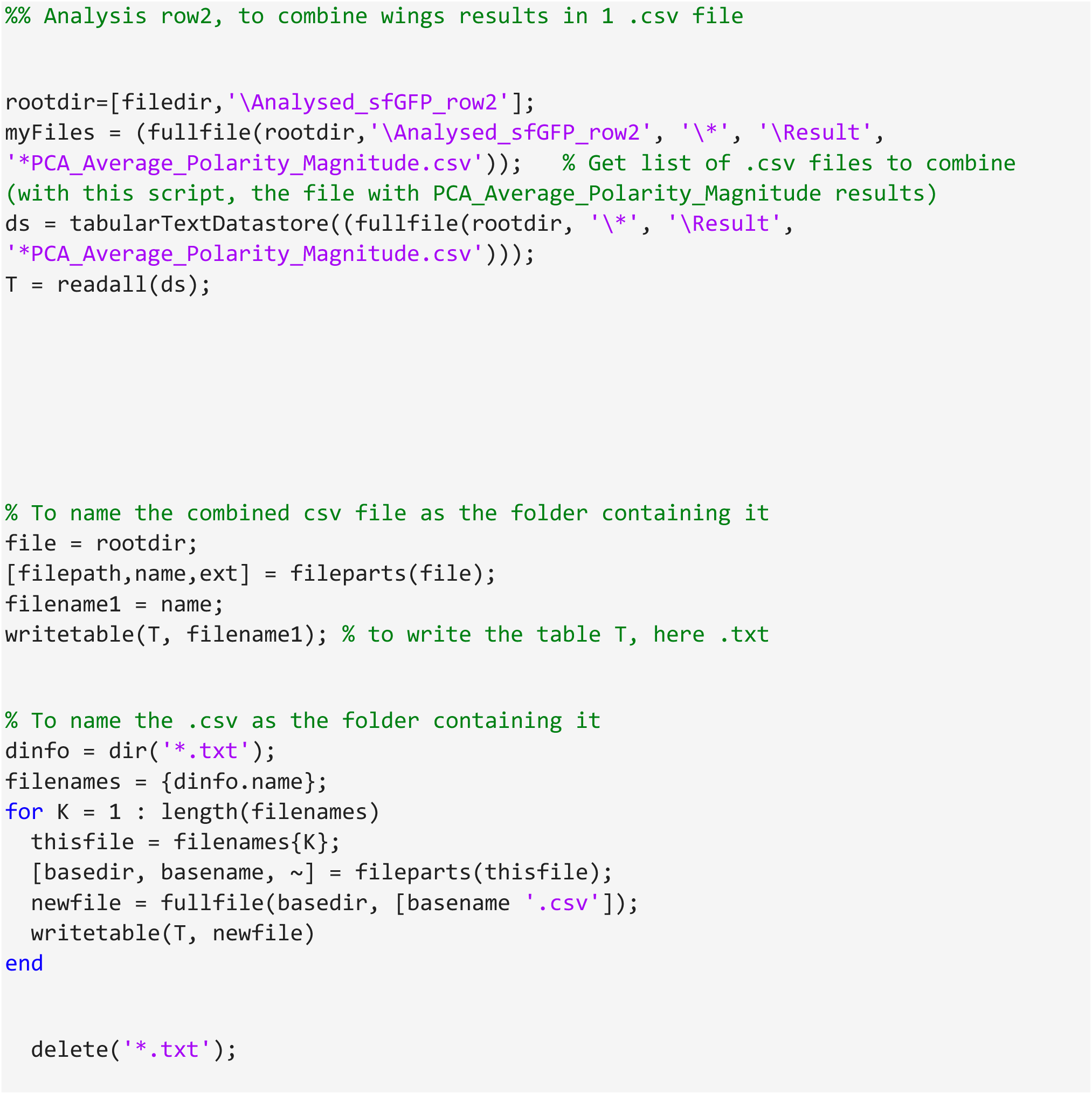

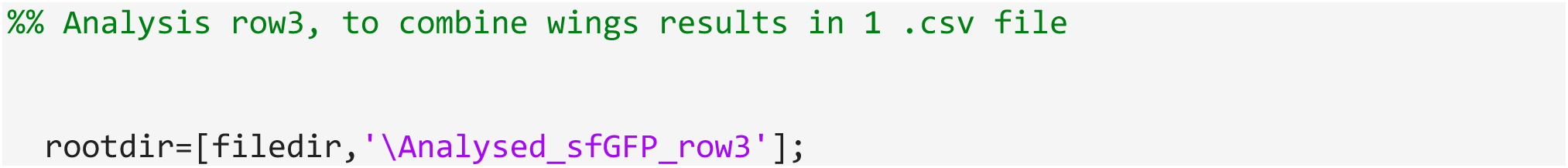
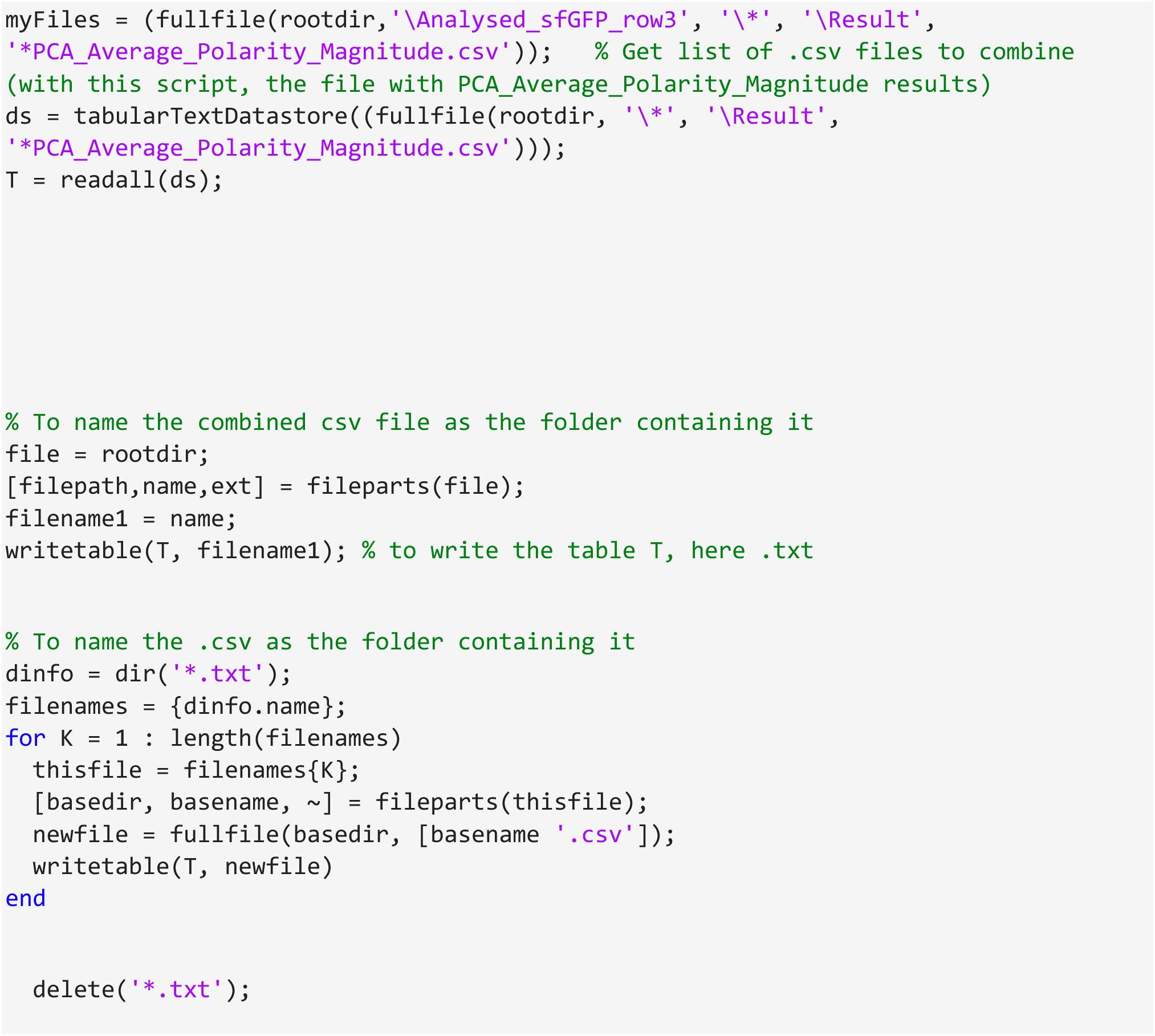

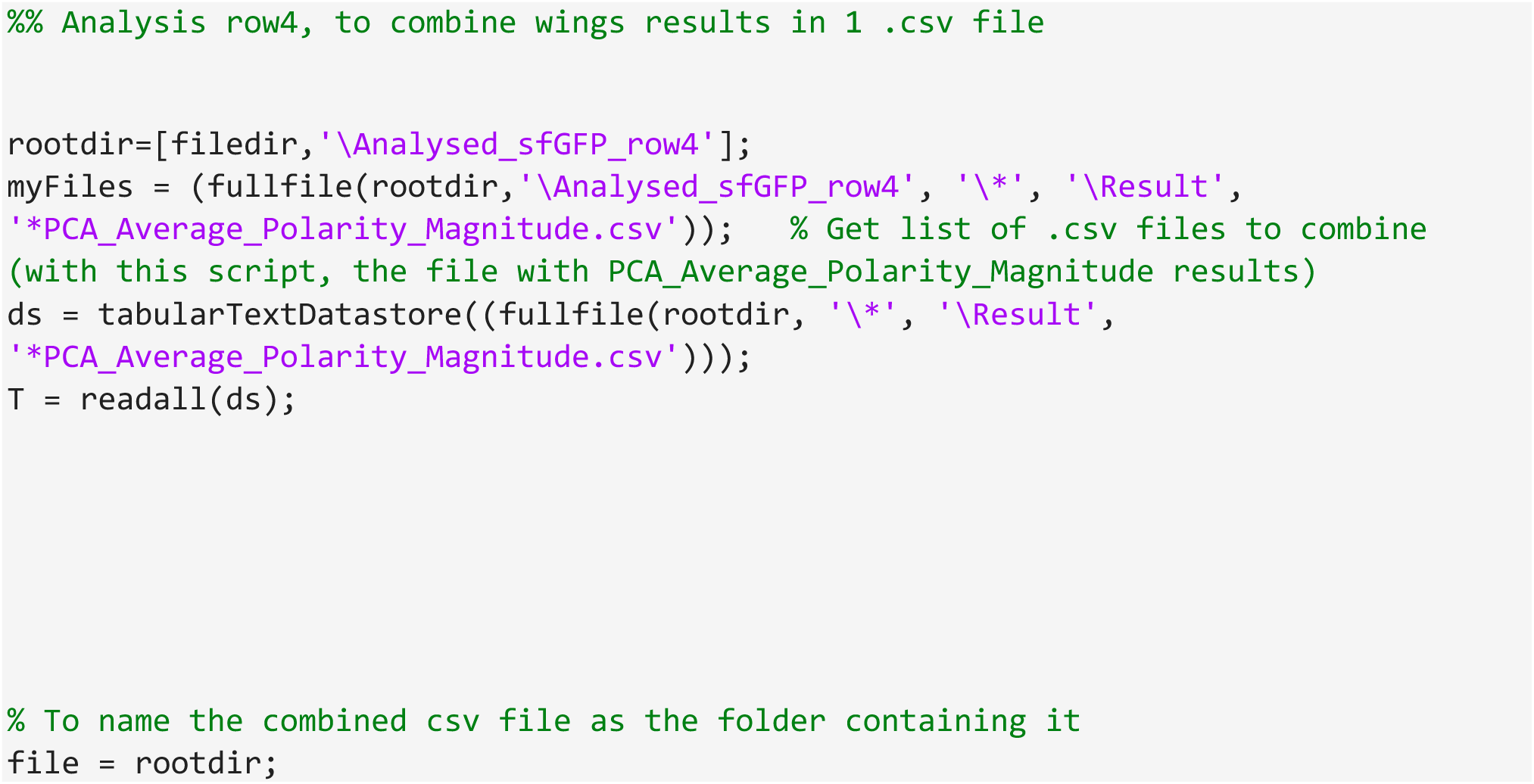
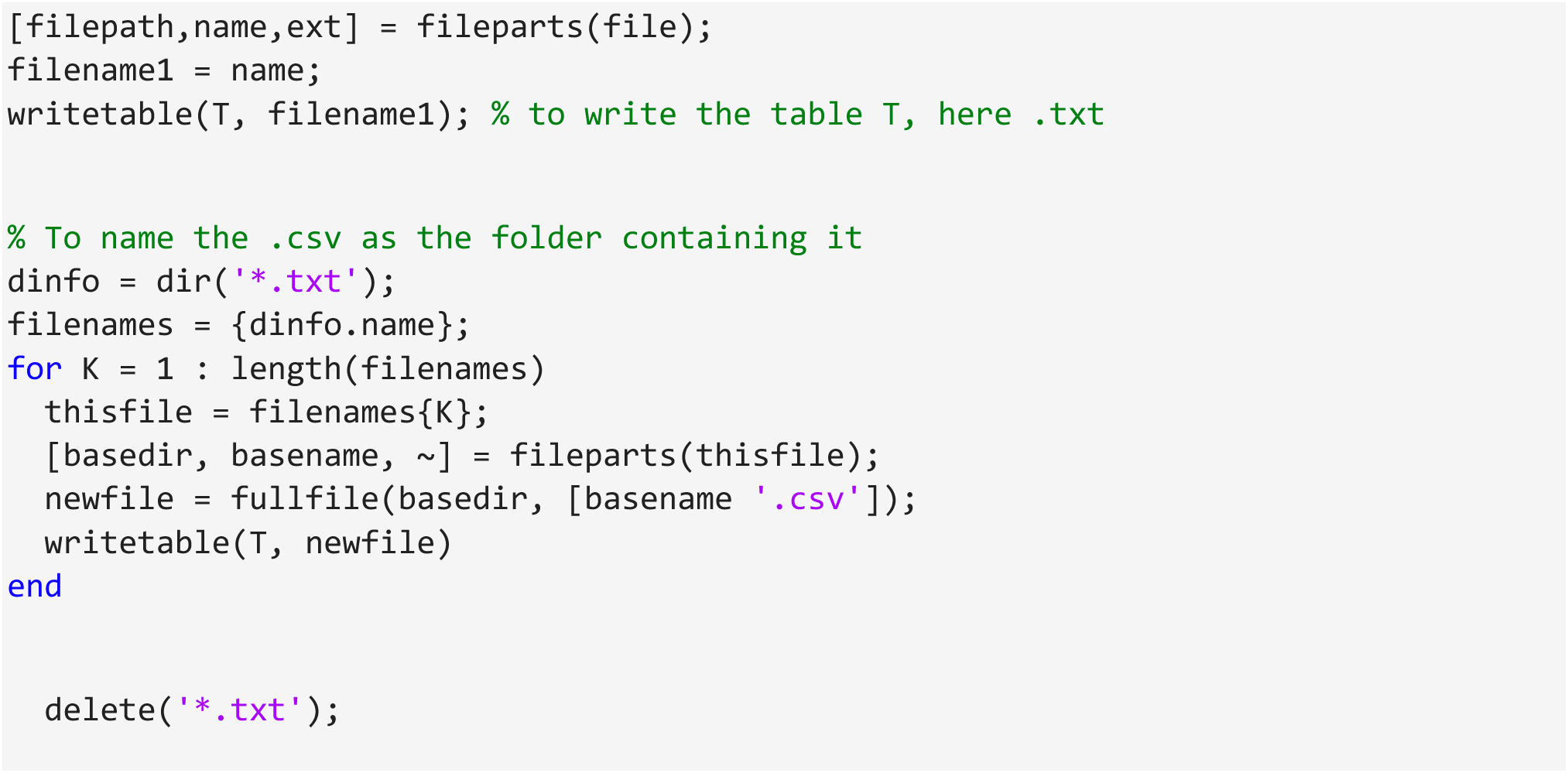

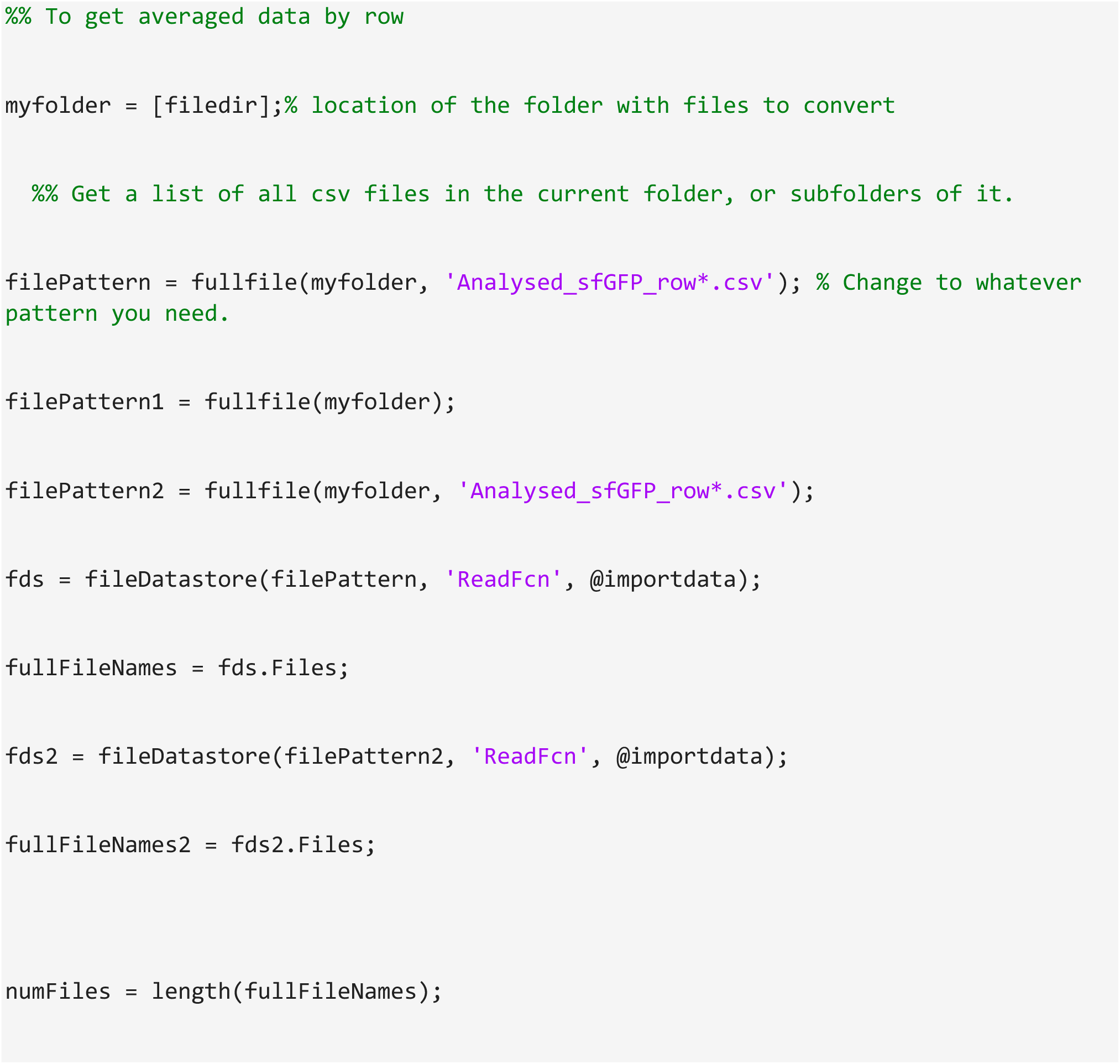
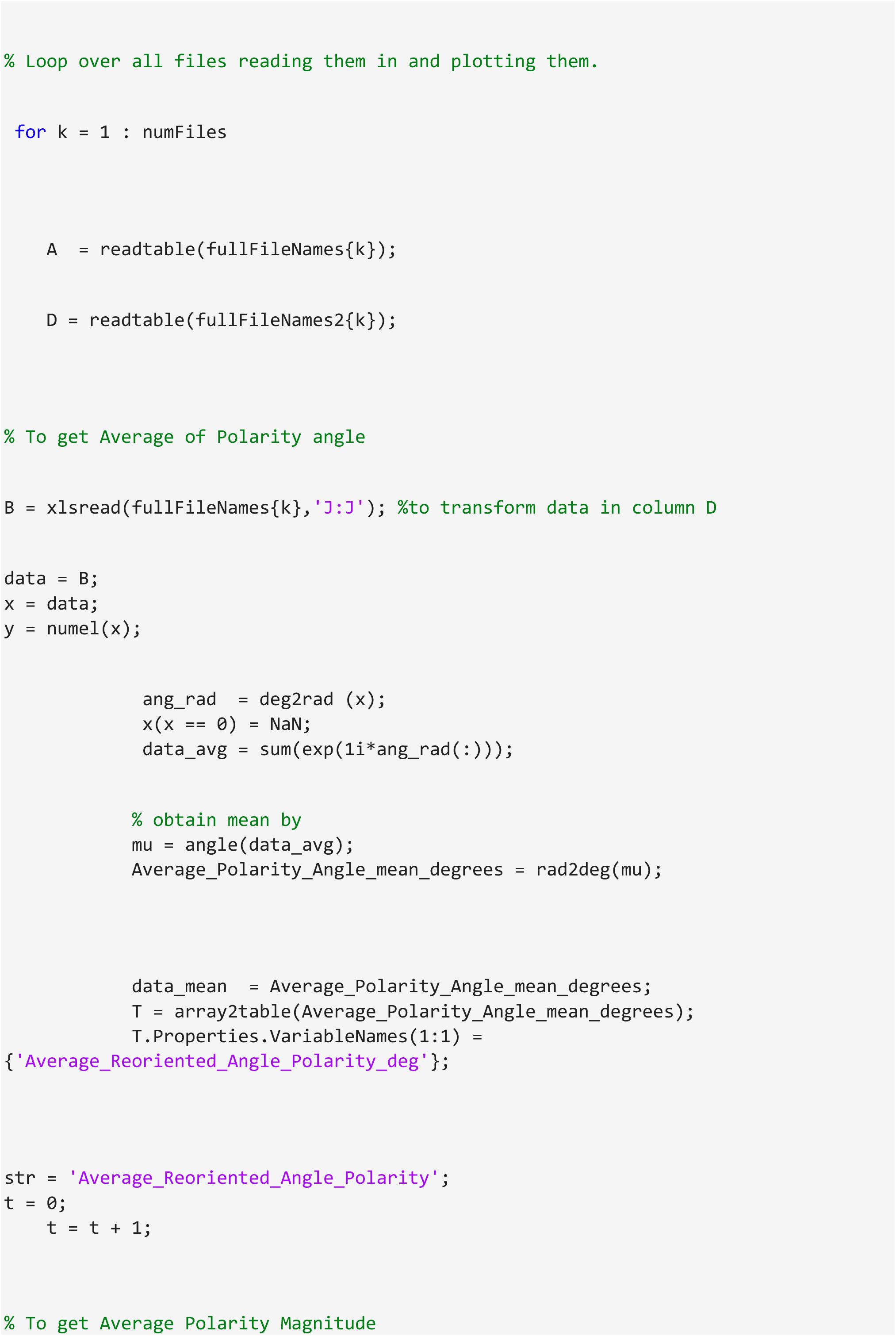
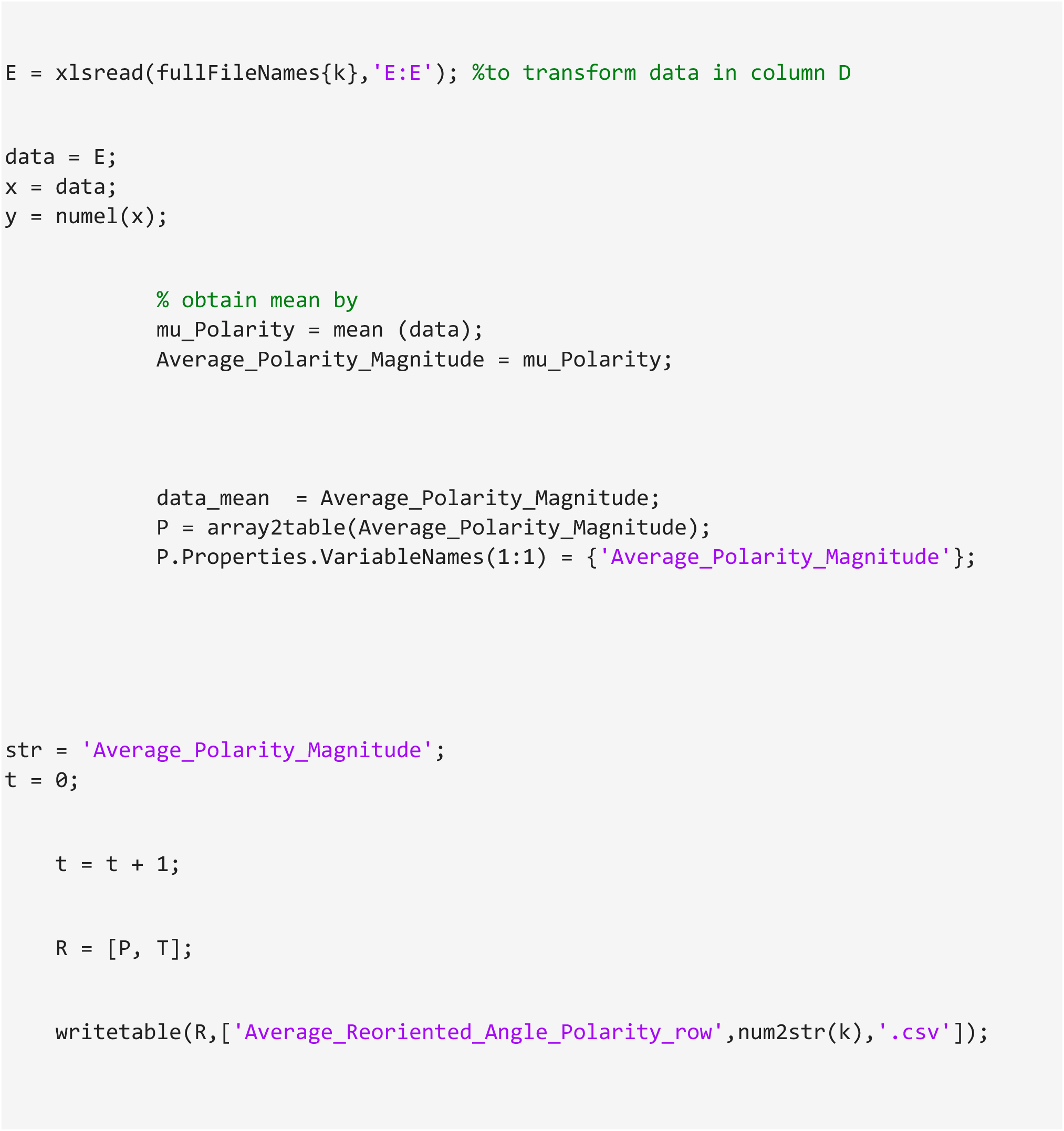

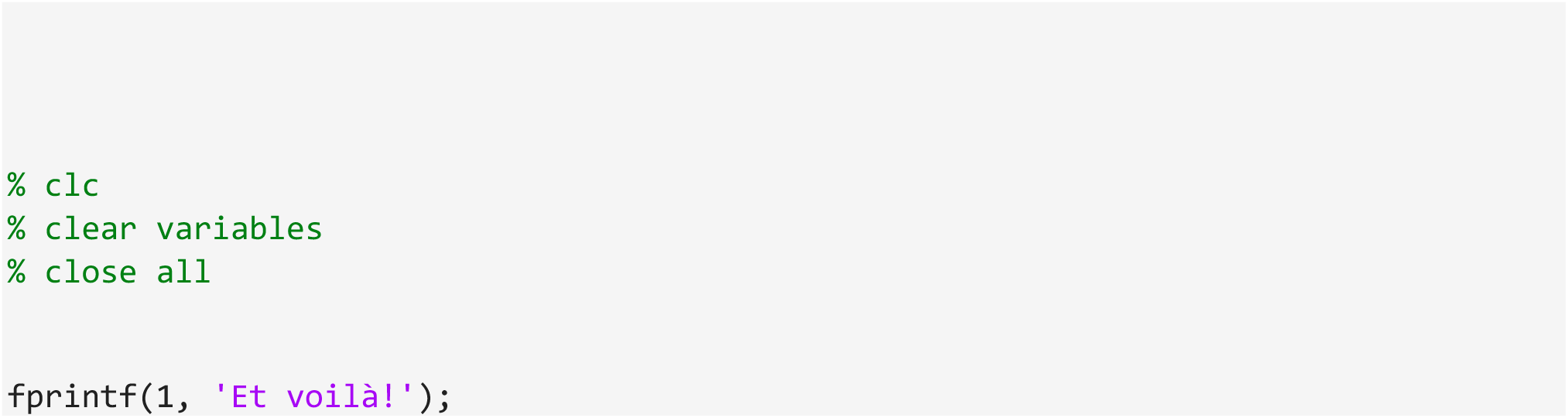

## Supplemental Materials – Analysis script 2

### Polar plot for polarity (magnitude + angle)

This is a MATLAB script suitable for generating polar plot showing polarity magnitude and angle, with results from ‘Combined results for polarity (magnitude + angle)’ MatLab script.

This script draw polar plot, with polarity magnitude and angle for each wing (empty circles) for one condition (example with row1) with results from ‘Analysed_sfGFP_row1’ that is file with averaged data per wing. It is added average polarity magnitude and angle for all wings from the same condition (filled circle) with results coming from ‘Average_Reoriented_Angle_Polarity_row1)

- Open the script, choose which condition to analyse by changing files name ‘’Analysed_sfGFP_row1.csv’’ and ‘’Average_Reoriented_Angle_Polarity_row1.csv’’ by proper files name.
- Run the script.
- Figure is saved in .pdf file with customised title.
- Circle colour and polar plot figure format can by change in the script following instructions.

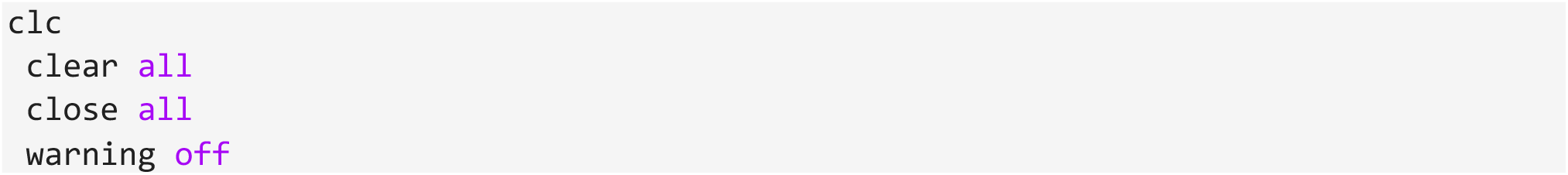
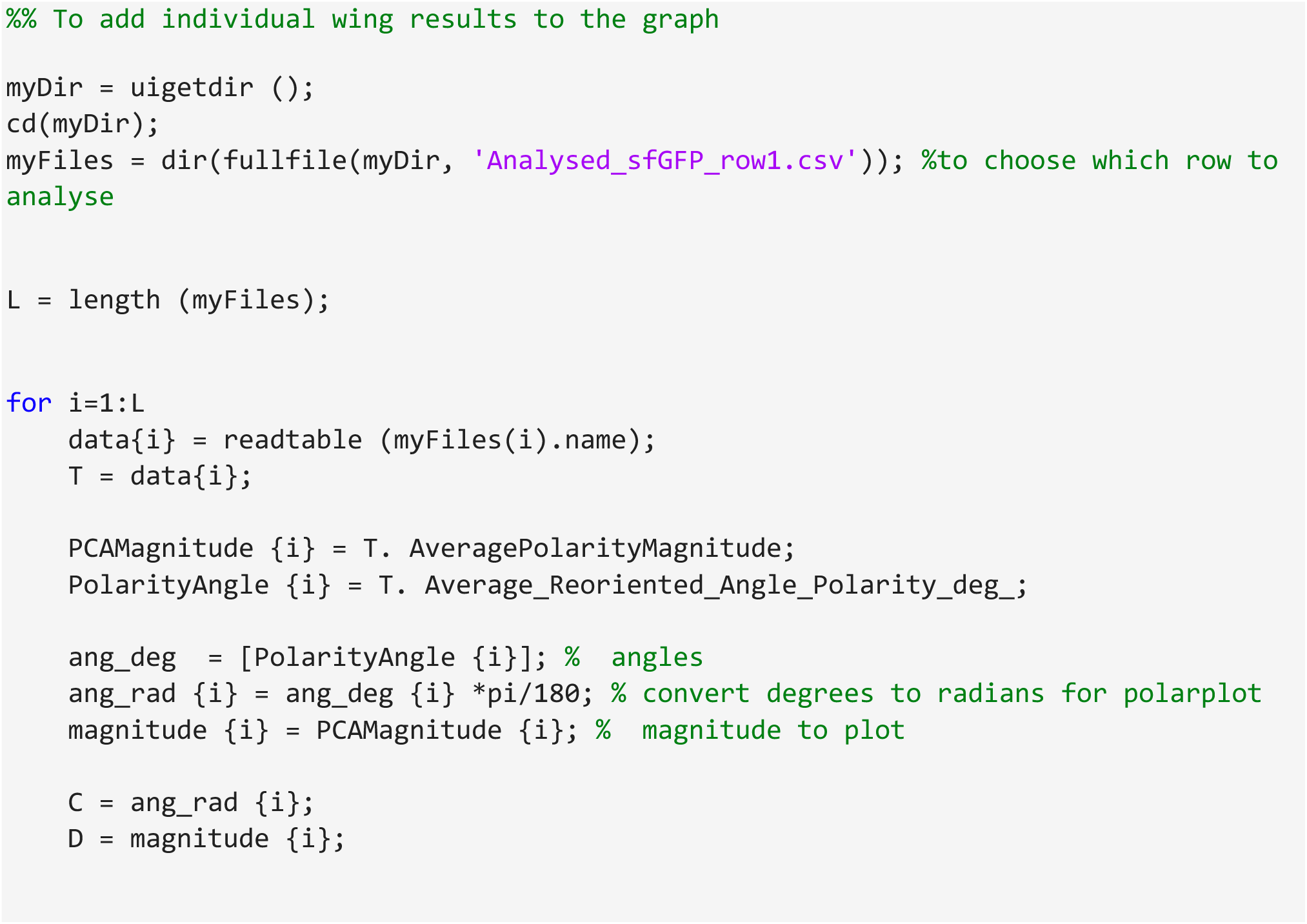
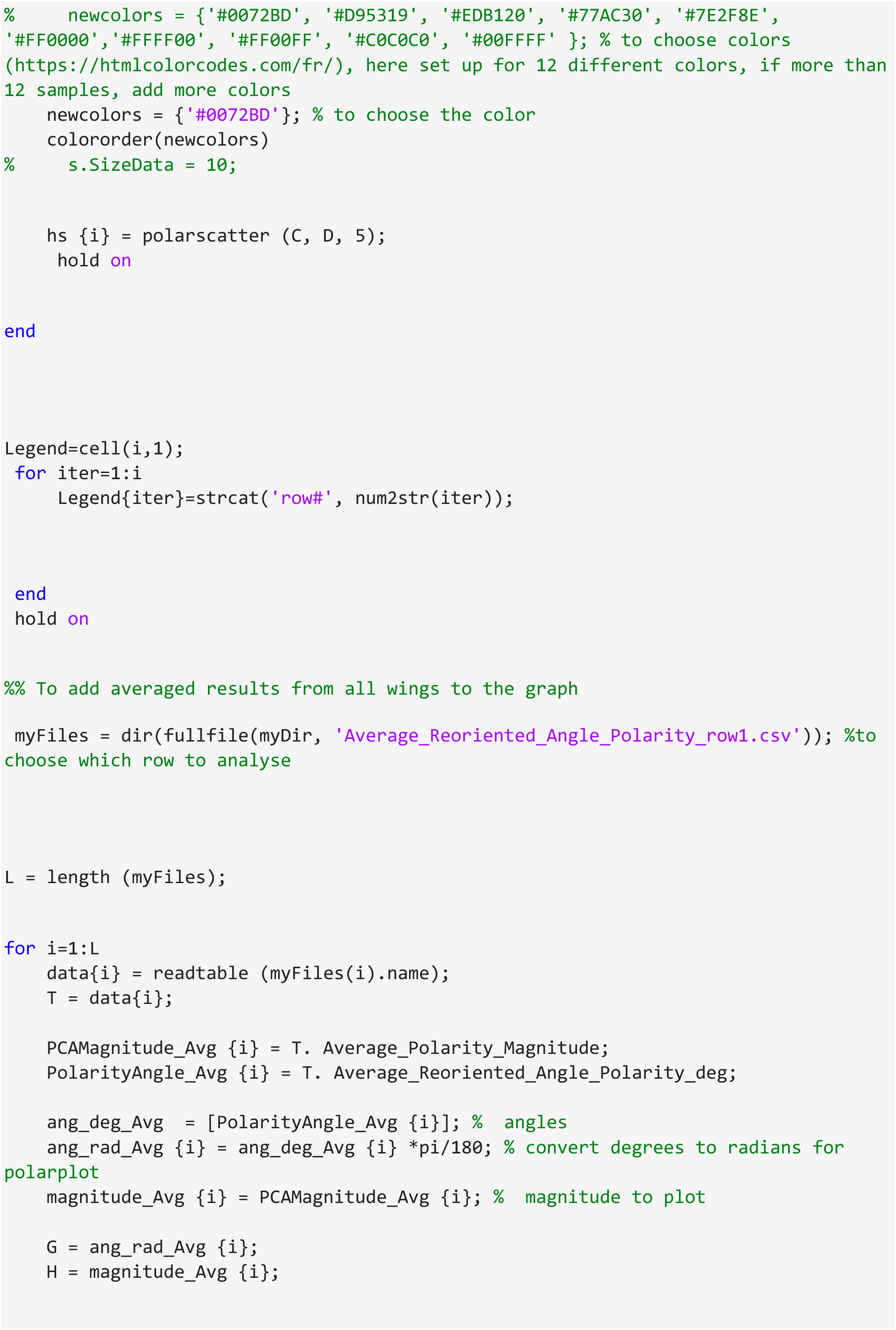
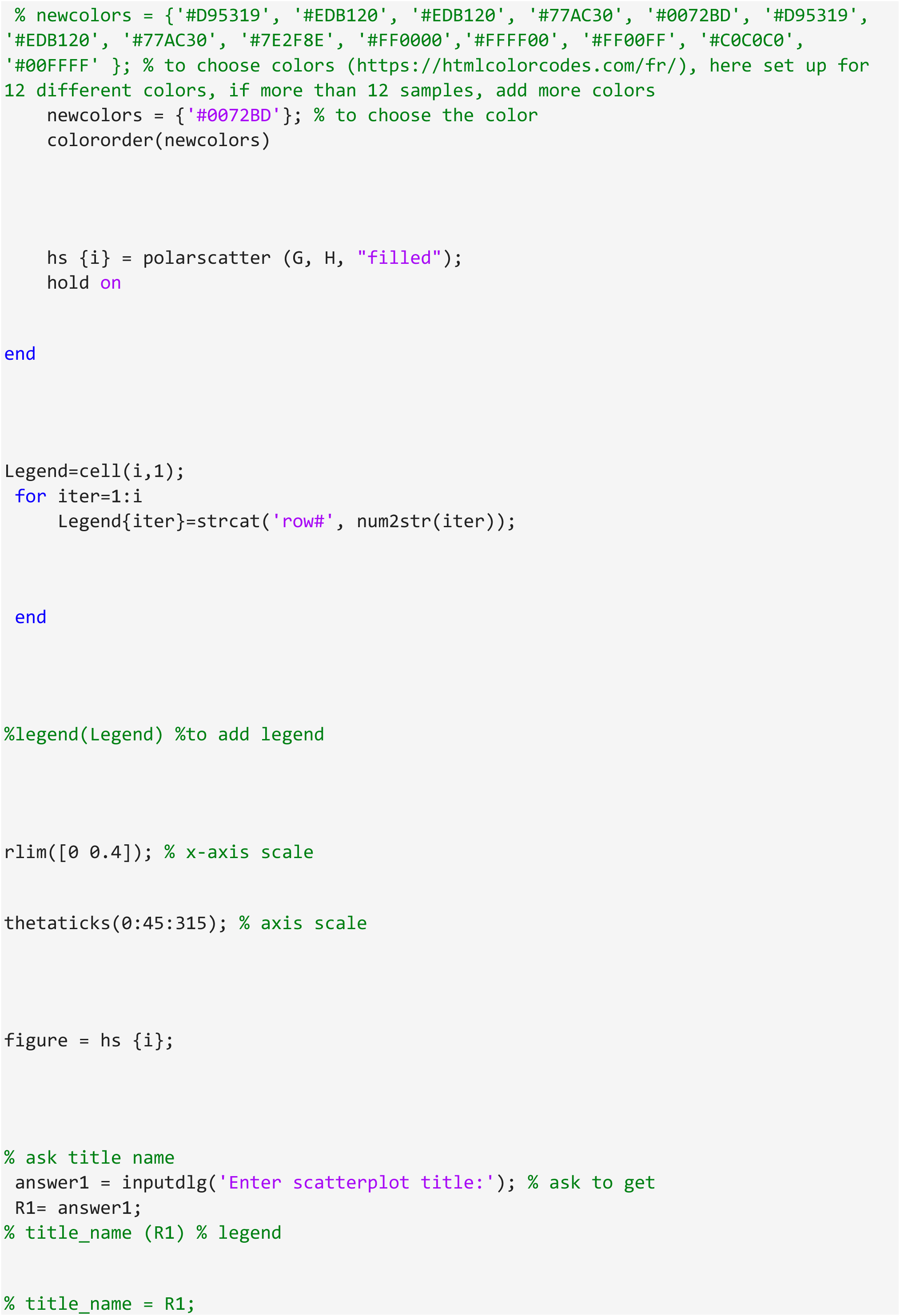
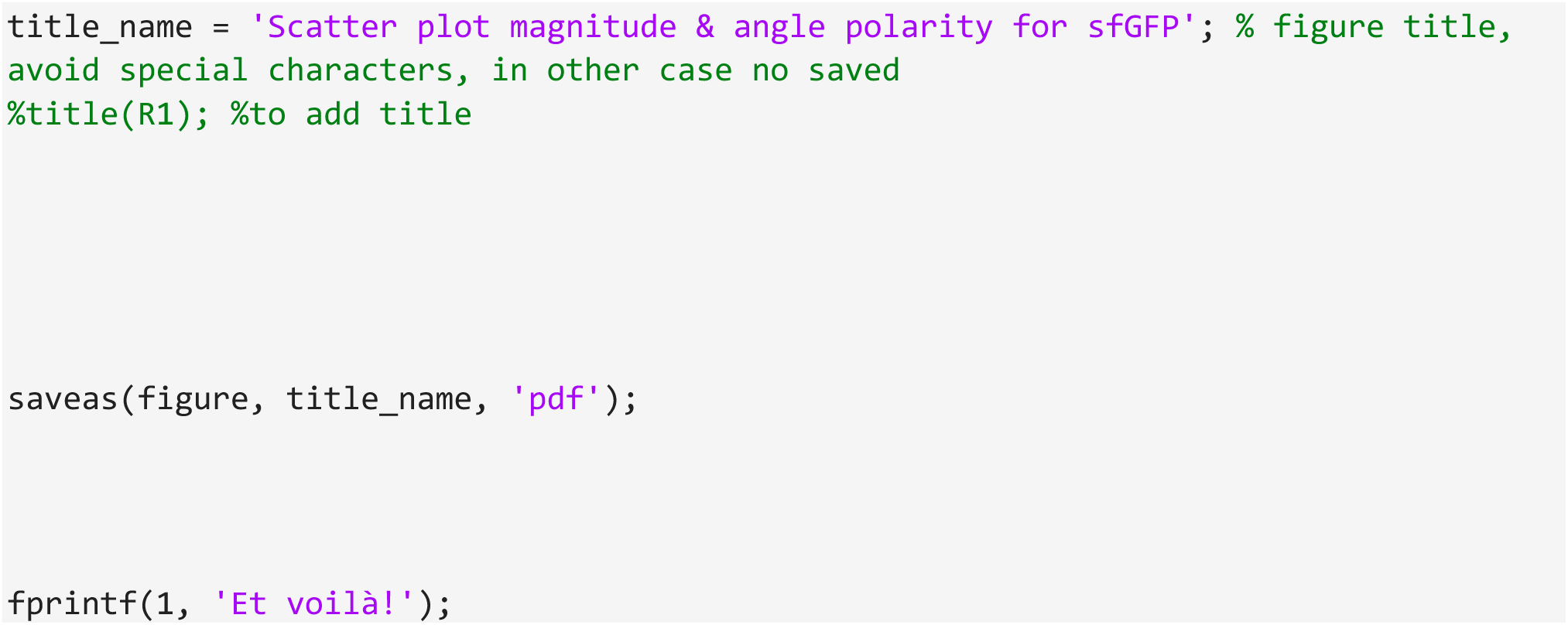

